# Genetic architectures of brain-related traits are shaped by strong selective constraints

**DOI:** 10.64898/2026.03.22.713538

**Authors:** Huisheng Zhu, Yuval B. Simons, Jeffrey P. Spence, Guy Sella, Jonathan K. Pritchard

## Abstract

Genome-wide association studies (GWAS) have identified hundreds of significant loci for psychiatric disorders, yet the strength of these associations remains modest compared to other human complex traits with similar numbers of hits. Whether this pattern reflects statistical artifacts or real biological differences — and, if the latter, what underlies it — remains unclear. In addition to psychiatric disorders, we find that other traits with functional enrichment in the central nervous system (CNS), whether binary or quantitative, also share similar genetic architectures, characterized by GWAS hits of limited statistical significance and generally higher allele frequencies. To robustly compare traits that differ in GWAS statistical power, we demonstrate how binarizing a quantitative trait reduces power. This loss of power can be replicated by a matched “effective sample size” on the liability scale. After matching “effective sample sizes”, we show that CNS-enriched traits have large mutational target sizes, with contributing variants and genes experiencing stronger selection than those for other traits. Our findings reveal heterogeneity among diseases and provide insights into traits that more effectively capture fitness-relevant processes. More broadly, our results suggest that the genetic architectures of complex traits are shaped by the tissues through which these traits are mediated.

## 1 Introduction

Many, if not most, traits in humans and in other species are genetically complex, with heritable variation spread over myriad variants throughout the genome. Complex traits have been studied for more than a century [1, 2], but only recently has it become possible to systematically dissect the genetic basis of their heritable variation. Over the past two decades, genome-wide association studies (GWAS) in humans have identified tens of thousands of robust associations between genetic variants and a wide array of quantitative traits and complex diseases [3, 4].

One intriguing observation from GWAS is that the genetic architectures of complex traits can vary widely. (Here, we use the term “architecture” to refer to the number of causal variants and their joint distribution of allele frequencies and effect sizes [4]. Often, we focus on the subset of approximately independent variants that exceed genome-wide significance in GWAS.) Notable trait variation in genetic architectures includes differences in the number and magnitude of significant associations detected within the same cohort [5, 6], as well as differences in the distributions of effect sizes [7] and allele frequencies [8] among significant variants.

Understanding the causes and determinants of this variation in architecture is of both practical and foundational importance. From an applied perspective, a trait’s genetic architecture determines the performance of studies aimed at mapping its genetic basis, as well as the accuracy of phenotypic predictions derived from these studies [9–12]. From a foundational perspective, the architecture of a trait reflects the evolutionary forces that shape heritable variation in natural populations and largely determines the genetic and phenotypic adaptive response to changes in selection pressures [13].

Recent work from Simons *et al*. tackled this problem from an evolutionary perspective [14, 15]. Motivated by extensive evidence that many quantitative traits are subject to stabilizing selection, with fitness declining with displacement from an optimal trait value [13, 16, 17], and that genetic variation affecting one trait typically affects many others [9, 18, 19], they modeled how selection on variants arises from stabilizing selection in a multi-dimensional trait space [14]. Their model also incorporated the effects of mutation and genetic drift. Given a demographic history and mutation rate per site, the genetic architecture of a trait in this model depends on two parameters and a distribution: the mutational target size (i.e., the number of sites in which a mutation affects the trait), the heritability per site in the target, and the distribution of selection coefficients for newly arising mutations in the target. Fitting this model to the 95 quantitative traits with more than a hundred approximately independent hits in the UK Biobank, they find that this model of pleiotropic stabilizing selection provides a better fit to the architectures of these 95 traits than previous ones [20–25] and that the distributions of selection coefficients are fairly similar across traits. Consequently, most of the variation in GWAS results across traits is driven by differences in the heritability per site in the target [15]. However, it remains unclear whether this result holds true for complex traits in general.

Several studies have suggested that the genetic architectures of psychiatric disorders and more generally of brain-related traits share characteristics that set them apart from other complex traits. We call a trait “brain-related” if variation in its value is largely mediated by genetic effects acting in cells of the central nervous system (CNS), and we identify such traits with stratified LD score regression (S-LDSC), testing for enrichment of SNP heritability in open chromatin regions of CNS cells [26, 27]. Brain-related traits are consistently found to be among the most highly polygenic traits examined in GWAS based on a variety of polygenicity measures [25, 28–30], suggesting that they have large mutational target sizes. Additionally, work by Holland *et al*. showed that under a point-normal model, non-null variant effect sizes for brain-related traits have variances that are orders of magnitude smaller than those for other complex traits [31]. When modeling the architecture with a flexible number of normal components, the effect sizes for brain-related traits were well approximated by a single normal distribution, whereas those for other complex traits typically required a mixture of two or more normals with different scales [32]. These lines of evidence suggest that brain-related traits arise from a large number of causal variants with relatively evenly distributed contributions to phenotypic variance.

Another consideration is that the traits Simons *et al*. examined were all quantitative; should we even expect complex diseases, including psychiatric disorders, to share similar properties in their genetic architectures? Intuition might suggest that genetic variation affecting complex diseases would be under directional selection, namely, that alleles that decrease the risk of disease would be selected for and alleles that increase this risk would be selected against. In contrast, the pleiotropic stabilizing selection model postulates that when mutations arise, they are always selected against, and for any given selection coefficient, they are equally likely to increase or decrease liability. Several recent studies support the latter view, indicating that, at least for common genetic variation, the architecture of complex diseases is predominantly shaped by pleiotropic stabilizing selection [7, 33, 34]. These findings therefore justify extending the Simons model to complex diseases.

Thus, we wanted to test whether the Simons model fits brain-related traits, including psychiatric disorders, and to explore how, and whether, these traits differ from other categories of traits. Do brain-related traits have unique genetic architectures? If so, what are the evolutionary causes that set brain-related traits apart? Here we set out to answer these questions.

We analyzed GWAS results across a wide range of complex traits and discovered that brain-related traits have unique features, with significant hits narrowly exceeding the significance threshold and enriched at higher allele frequencies. After adjusting for statistical power, which is influenced by sample size differences and the binary nature of diseases, we conclude that psychiatric disorders, along with other brain-related quantitative traits, have large mutational target sizes, and their underlying variants and genes are experiencing stronger selection.

## 2 Results

Substantial variation in genetic architecture revealed by GWAS can be exemplified with three contrasting traits: a non-brain-related quantitative trait — low-density lipoprotein cholesterol (LDL) [35]; a non-brain-related complex disease — coronary artery disease [36]; and a brain-related complex disease — schizophrenia [37]. While these GWAS yield similar numbers of approximately independent hits (Methods), the significant associations for schizophrenia surpass the genome-wide significance threshold by only a small margin (Figure 1A).

**Figure 1:**
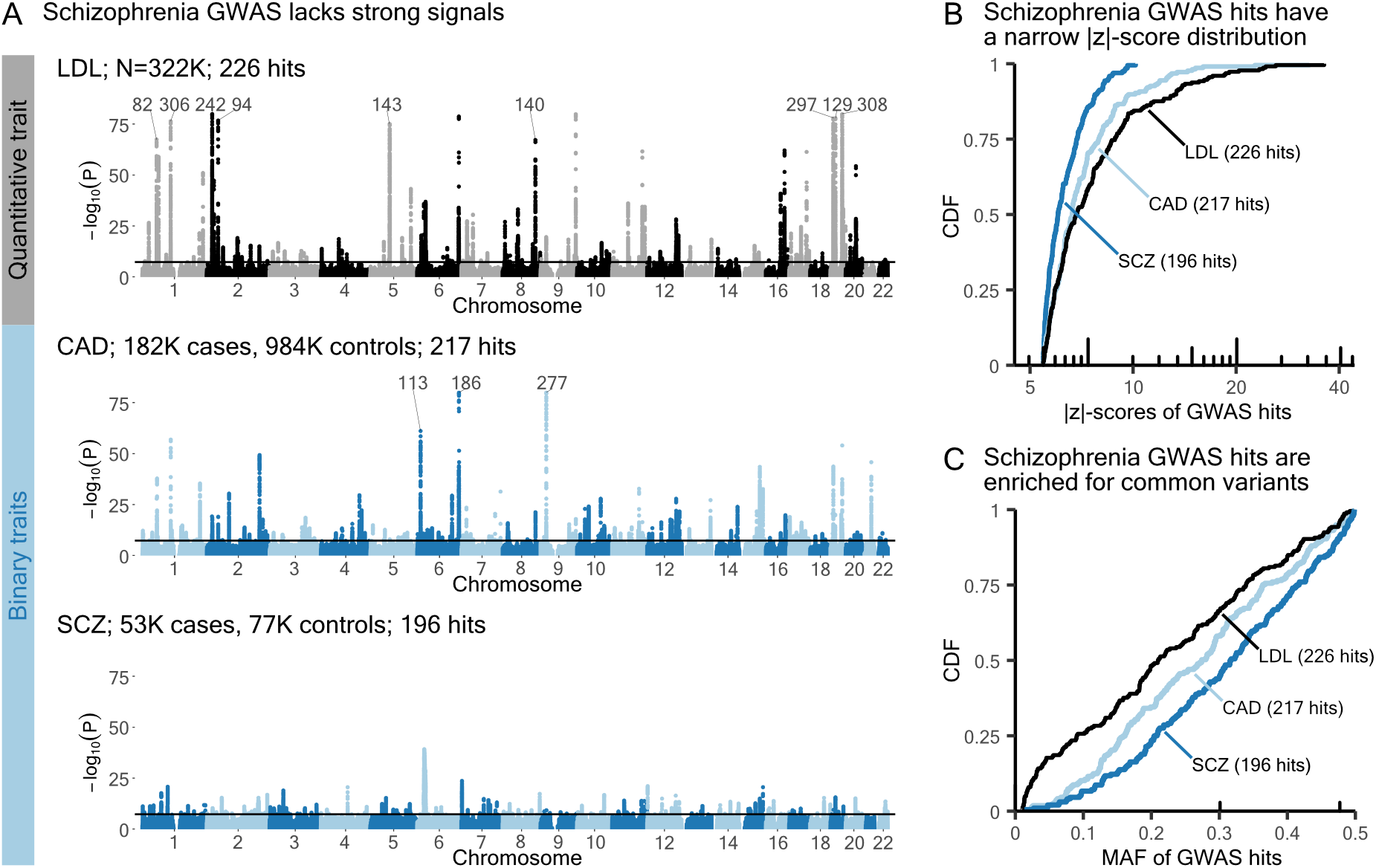
Schizophrenia GWAS hits narrowly exceed the significance threshold. ***A)** Manhattan plots for three example traits. The y-axes of all three plots are restricted to the range* [0, 80]. *Significant associations beyond this range are labeled with the most significant p-value. Independent hits are obtained from GCTA-COJO [38], and hits with MAF below 1%, in the HLA region, or of low imputation quality are excluded (Methods). **B)** Cumulative distributions of COJO hit absolute z-scores. **C)** Cumulative distributions of COJO hit MAF. CAD: coronary artery disease; SCZ: schizophrenia.*

Differences in strength of significant associations are more readily apparent in the cumulative distributions (CDFs) of |*z*| -scores of their approximately independent hits (Figure 1B). Schizophrenia, which lacks strong signals, has a steeper CDF curve. When we instead plot the cumulative distributions of MAF, a new piece of information emerges: schizophrenia GWAS hits are generally higher frequency (Figure 1C).

Plots in the style of Figures 1B and C provide useful summaries of genetic architectures across traits. Based on these results, we wanted to understand why schizophrenia has such a distinct genetic architecture.

### The distinct genetic architectures of brain-related traits

We asked whether these differences in architecture are seen more generally between brain-related and other complex traits. To this end, we considered the 151 quantitative traits in the UK Biobank with at least 20 approximately independent hits and SNP heritability above 5%, as well as 13 human complex diseases with publicly available case-control GWAS summary statistics that meet these same criteria (Supplementary Tables 1-2, Methods).

We classified these traits as brain-related or non–brain-related by applying a significance threshold of 0.05/(10 *×* 164) to the meta-analyzed S-LDSC [26, 27] p-values for CNS enrichment (Methods). This classification yielded 50 brain-related quantitative traits, including seven behavioral-cognitive traits, and 101 non-brain-related quantitative traits (Figures S4 and S5). We also identified four brain-related diseases — all of which are psychiatric disorders — and nine non–brain-related diseases (Figure S6). Among brain-related traits, psychiatric disorders and behavioral-cognitive traits show the strongest CNS-specific enrichment (Figure 2A).

**Figure 2:**
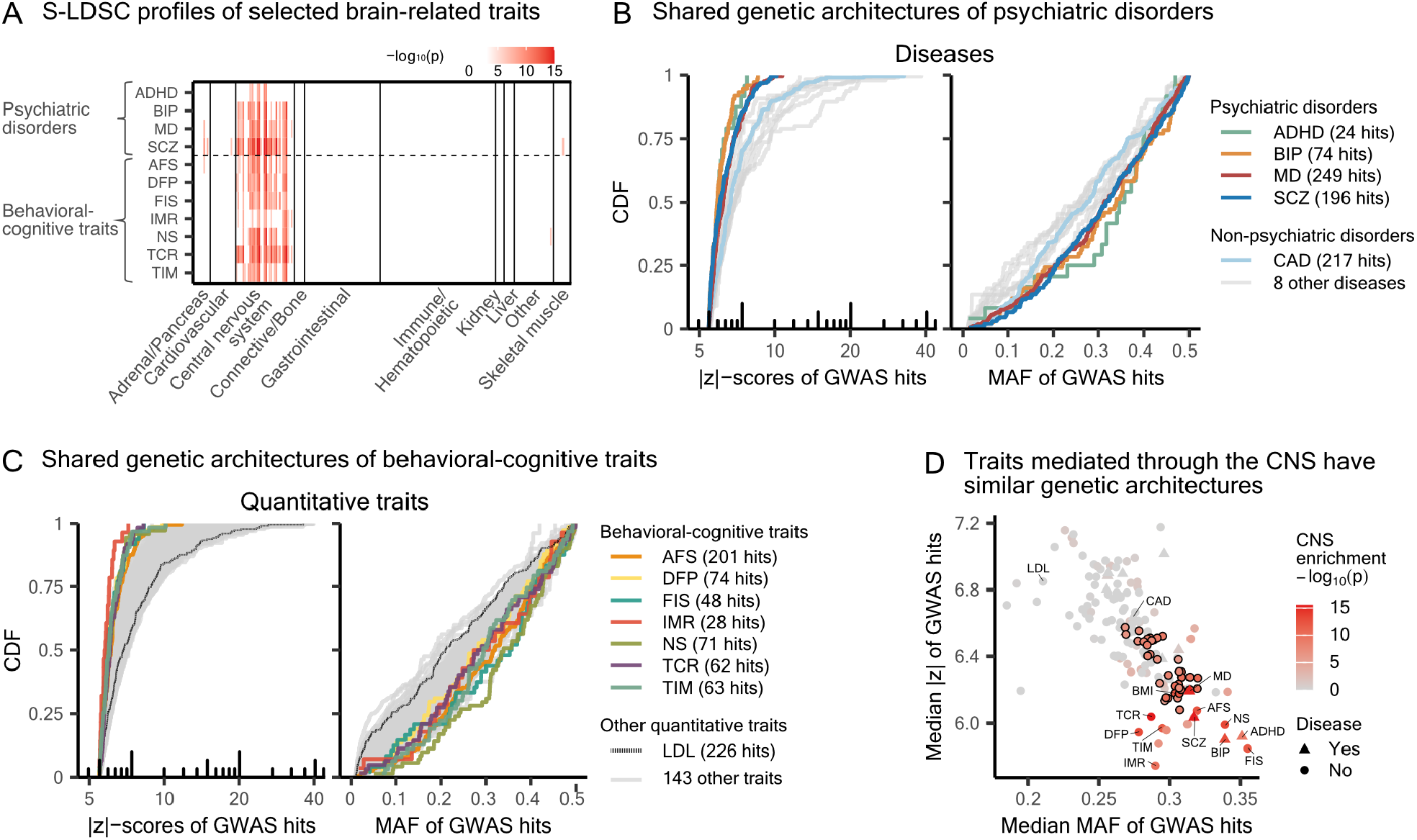
Traits with functional enrichment in the CNS share similar GWAS hit patterns. ***A)** Functional enrichment analyses with S-LDSC [26, 27] on selected traits. Each row consists of 220 cell types from 10 categories, colored by −log*_10_*(p) for the coefficient τ if significant after Bonferroni correction. The gradient scale is bounded above at 15. Results on all other traits are in Figures S4 to S6. **B)** Contrasting the GWAS hits for psychiatric disorders with those of nine other complex diseases. The full list of diseases can be found in Supplementary Table 2. **C)** Contrasting the GWAS hits for behavioral-cognitive traits with those of 144 other quantitative traits. The full list of quantitative traits is in Supplementary Table 1. **D)** GWAS hits for traits enriched for the CNS have higher median MAF and lower median |z|-score. Each trait included in this study is colored by the combined p-value for CNS enrichment, computed following the aggregate Cauchy association test (ACAT) [41] (Methods). The points corresponding to the 36 body composition–related traits are outlined with black borders. ADHD: attention deficit hyperactivity disorder; BIP: bipolar disorder; MD: major depression. AFS: age first had sexual intercourse; DFP: duration to first press of snap-button in each round; FIS: fluid intelligence score; IMR: number of incorrect matches in round; NS: neuroticism score; TCR: time to complete round; TIM: mean time to correctly identify matches. BMI: body mass index.*

As in the schizophrenia example, we find that brain-related traits, including both quantitative traits and diseases, generally have GWAS hits spanning a narrower range of statistical significance and are enriched for higher MAF compared with other complex traits (Figures 2B and C). By contrast, neurological diseases — including Alzheimer’s disease, Parkinson’s disease, and multiple sclerosis — are classified as non–brain-related, and do not exhibit the genetic architecture observed for brain-related traits (Figure S7).

At a fixed sample size, the expected strength of association for a given variant *i* in GWAS is determined by its MAF *p*_*i*_ and squared effect size 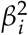; more precisely, 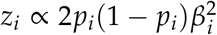 (see, e.g., supplementary information of [39]). Despite being enriched for common variants, GWAS hits for brain-related traits show narrower distributions of |*z*| -scores. This apparent contradiction is attributable to their narrower distributions of squared effect sizes relative to other complex traits (Figure S8), echoing previous findings [31, 32].

The narrow distributions of |*z*| -scores, elevated MAF, and small effect sizes place the genetic architectures of brain-related traits at an extreme among complex traits (Figures 2D, S9 and S10). Traits mediated through other tissues, including the adrenal gland and pancreas, kidney, and skeletal muscle, also cluster in this parametrization of architecture, but none are set apart from other traits as much as those mediated by the CNS. If a trait is mediated by the CNS as well as cells in other tissues, we might expect its genetic architecture to be intermediate between traits exclusively enriched for the CNS and non-brain-related traits. This is what we see for body composition-related traits (Figures 2D and S11B), which are partially mediated by the CNS through appetite control and preference for food intake [40], but are also mediated by other tissues (Figure S11A). Here, we focus on the pronounced differences between brain-related traits and other traits, but we revisit the potential roles of other tissues on trait genetic architectures in the Discussion.

Importantly, we wanted to ensure that these patterns were not driven by uncontrolled confounding in population-based GWAS of these traits. By comparing results from population-based and family-based association studies, previous studies have suggested that confounders including population stratification, indirect genetic effects, and assortative mating, can aggregate across the genome to bias heritability and genetic correlation estimates [42–45]. However, our analyses focus on GWAS hits, and we expected confounding effects at these strong-signal sites to be minor [45, 46]. To check this, we used a GWAS of birth coordinate as a negative control where most signals, except a few protective against allergies [47], are expected to be driven by confounding. MAF of GWAS hits for birth coordinate traits have a cumulative distribution that is distinct from any of the traits we considered (Figure S12).

While birth coordinates serve as a negative control trait, we were intrigued by urinary biomarkers, some of which were classified by our pipeline as brain-related and are less obviously prone to confounding from population stratification, assortative mating, or indirect genetic effects as compared to behavioral-cognitive traits. We noticed that creatinine and sodium levels in urine are classified as brain-related, while the level of creatinine in blood is not (Supplementary Table 1, Figure S13A). This is because in clinical practice, serum creatinine assesses the kidney’s normal function of eliminating waste products from muscle metabolism, whereas urinary creatinine is often used to normalize other urinary biomarkers, given that creatinine excretion is kept at a relatively constant rate [48]. Thus, unadjusted urinary creatinine and sodium concentrations are indicative of hydration status and urine flow rate, which involves coordination between the brain and bladder [49]. GWAS results of these urinary traits more closely resemble those of other brain-related traits than those of serum creatinine, which again supports that traits mediated through the CNS share similar genetic architectures (Figure S13B). Overall, this suggests that confounding is not driving the distinct patterns we see for the brain-related traits in our analyses, including the behavioral-cognitive ones.

### Brain-related traits retain distinct architectures after adjusting for statistical power

To ensure a rigorous comparison between brain-related and non-brain-related traits, we wanted to account for the binary nature of disease traits and for variation in sample size across traits. Both factors influence GWAS statistical power and, consequently, the learned genetic architectures.

We begin by examining binary traits, for which GWAS is typically conducted using logistic regression rather than the linear regression applied to quantitative traits. To study the impact of this difference, we converted quantitative traits into binary traits by applying a cutoff to the phenotypic values, a procedure we refer to as “binarizing” a trait. Our goal was to determine sample sizes that yield equivalent GWAS power between the binarized and the original continuous versions of essentially the same trait. Specifically, we binarized LDL levels using thresholds corresponding to varying population prevalences of the resulting binary trait (Figures 3A and S14A, Methods).

**Figure 3:**
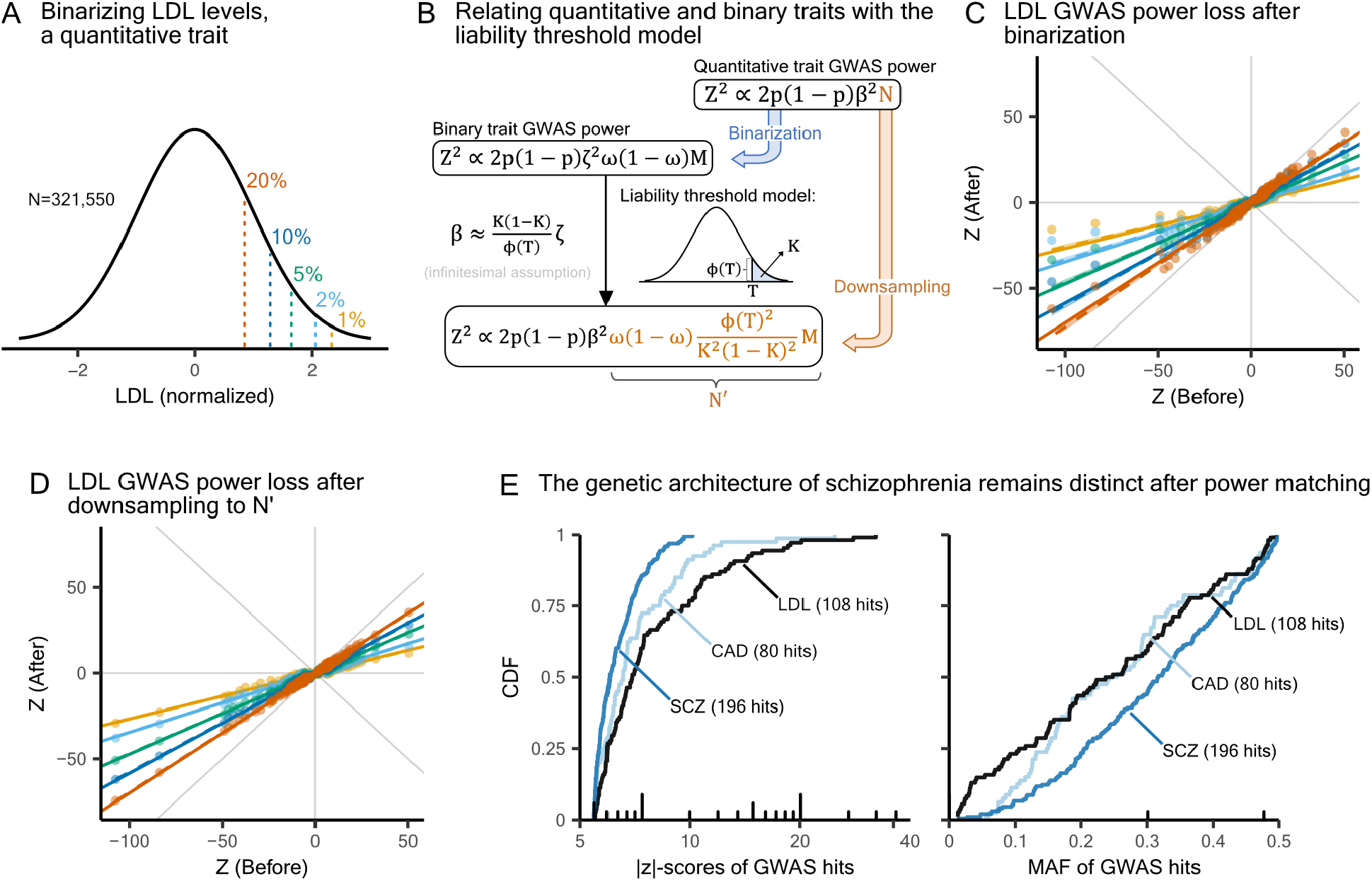
Calibrating GWAS statistical power between quantitative traits and binary traits. ***A)*** *Binarizing LDL with varying prevalence in the upper tail. LDL levels were measured in unrelated White British individuals from the UK Biobank. The original phenotypic distribution was standardized using rank-based inverse normal transformation*. ***B)*** *Derivation of the sample size needed on the liability scale to reach equivalent power of a case-control GWAS. MAF is denoted by p. We transferred effect sizes from log odds ratio scale (ζ) to continuous scale (β) using the liability threshold model. For details, see Supplementary Notes*. ***C)*** *Deflation of LDL GWAS hit z-scores after binarizing. Different colors reflect the threshold used to binarize traits, as shown in panel A. Solid lines represent predicted slopes, and dashed lines indicate fitted slopes. Gray lines in the background are y* = 0, *x* = 0 *and y* = *±x*. ***D)*** *Deflation of LDL GWAS hit z-scores after downsampling to matched sample sizes. Color choices, line styles, and background lines are identical to those in panel C*. ***E)*** *Reducing coronary artery disease and LDL GWAS power to match with the current effective sample size of schizophrenia GWAS, N*^*′*^ = 192,273 *(Methods)*.

Based on the classical liability threshold model [50], an individual’s liability is a quantitative trait that relates to their genotype through the standard additive model, and individuals with liability exceeding a threshold *T* develop the disease. This model allows us to map case-control GWAS results for complex diseases to the corresponding GWAS results on the underlying liability, which is typically unobserved. Assuming variant effect sizes are sufficiently small, the power to identify a variant in a case-control study with a sample size of *M* individuals, with a proportion *ω* of cases and (1 − *ω*) of controls, is approximately equal to the power to identify that variant in a GWAS of liability with effective sample size

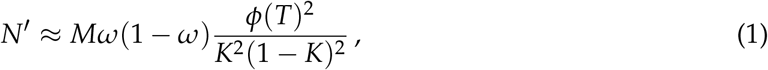

where *K* is the population prevalence of the disease, and *ϕ*(·) denotes the standard normal density (Figure 3B, Supplementary Note).

In our analysis, we can treat the original trait as the underlying liability for the binarized trait. Accordingly, we used independent hits from the original quantitative LDL GWAS and evaluated the extent to which their *z*-scores were deflated in case-control GWAS across different binarization thresholds (Figures 3C and S14B). We also performed quantitative LDL GWAS on a subsample whose size was determined by Equation 1 to match the reduction in power caused by binarizing for each threshold we used (Figure 3D). Both binarizing and downsampling are expected to approximately linearly deflate *z*-scores with

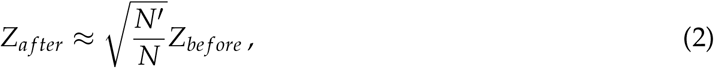

where *N* is the sample size of the original quantitative trait GWAS (Supplementary Note).

Figures 3C and D share the same set of predicted lines as given by Equation 2, and our theory accurately predicts the effects of both binarizing and downsampling. Variants of large effect deviate from the linear relationships in Figure 3C and S14B but not in Figure 3D, because Equation 1 relies on the infinitesimal assumption, whereas Equation 2 does not (Supplementary Note).

We repeated this analysis with five other medically relevant quantitative traits in the UK Biobank to ensure that these results were not unique to LDL (Figures S15 to S19). Unexpectedly, many traits exhibit asymmetrical power reduction when binarizing in the lower tail compared to the upper tail. For example, BMI shows more deflation than expected when binarizing in the lower tail, and less deflation in the upper tail (Figure S15). This pattern suggests that the same set of variants have, on average, stronger effects in high-BMI individuals than in low-BMI individuals. One possible explanation for this tail asymmetry is genetic epistasis, in which the magnitude of a variant’s effect varies depending on an individual’s genetic background, resulting in larger effects at one end of the liability distribution compared to the other. Gene-environment interaction is another plausible mechanism and has been proposed to explain why polygenic scores achieve higher predictive accuracy in higher BMI quantiles than in lower ones [51]. While intriguing, these deviations from our theory are relatively minor and determining the precise mechanism is beyond the scope of the current study.

Given the accuracy of our predictions, we can adjust for GWAS power by first computing *N*^*′*^ for each binary trait and then scaling *z*-scores according to Equation 2. We estimated that *N*^*′*^ is approximately 192*K* and 333*K* for the current schizophrenia and coronary artery disease GWAS, respectively (Methods, Supplementary note, Supplementary Table 3). Even after matching all three example traits in Figure 1 to the same *N*^*′*^, we continue to see differences in the CDFs of |*z*| -scores and MAF for independent GWAS hits (Figure 3E). This more rigorous comparison shows that differences in GWAS power do not explain why brain-related differ from other traits in genetic architectures, suggesting that other biological or evolutionary mechanisms are involved.

### Brain-related traits have large mutational target sizes and relevant variants are under strong selection

To formally investigate alternative explanations, we turned to the generative model for GWAS hits of a quantitative trait previously developed by Simons *et al*. [15]. We extended this model to accommodate binary traits using Equation 1 (Figure 4A). In this framing, mutations affecting the trait arise at a rate that depends on the trait’s mutational target size *L*, and are then assigned a selection coefficient *s* > 0 from a distribution *f* (*s*). The frequency of the mutation at the time the GWAS is performed, *p*, is sampled from a distribution that arises from under-dominant selection with the selection coefficient *s* and the effects of genetic drift during the demographic history of the population. We assume a demographic model estimated for the British population [52] and a mutation rate of 1.25 *×* 10^−8^ [14]. The effect size of the mutation, *β*, also depends on *s*: specifically, *β*|*s* ∼ *N*(0, *c* · *s*), where *c* is a trait-specific constant that depends on the heritability per site in the target, *h*^2^/*L*. The variant is then assigned a z-score sampled from a distribution that depends on its effect size, allele frequency, and GWAS sample size. When this z-score exceeds the genome-wide significance threshold, the variant is considered a GWAS hit. Below, we fit *f* (*s*) using a spline function with four knots (see Supplementary Section 4.8 in [15]), thus our full model includes six parameters per trait: four for *f* (*s*), as well as *h*^2^/*L*, and *L*.

**Figure 4:**
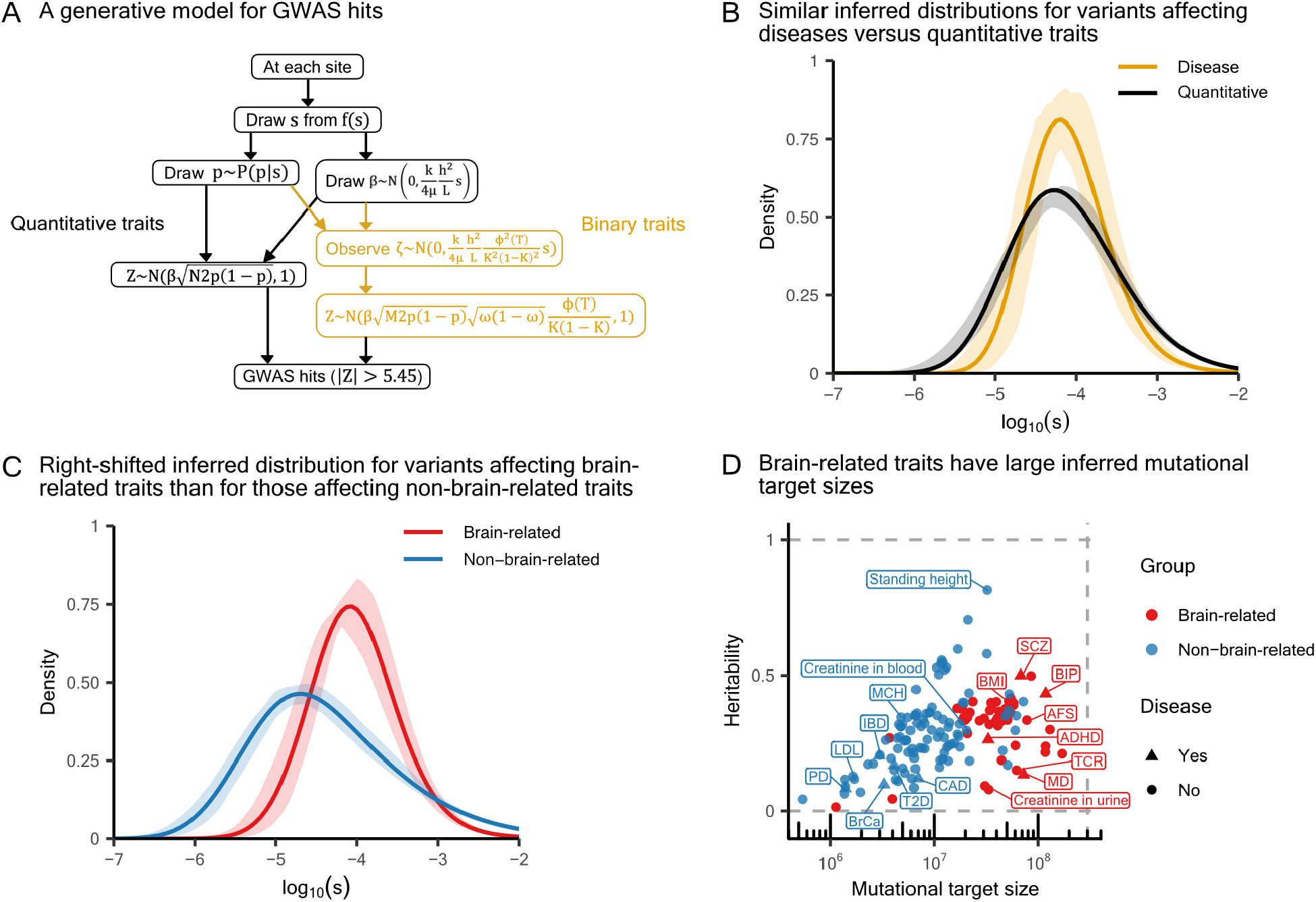
Stronger selection on brain-related variants and larger mutational target sizes for brain-related traits. ***A)*** *The model that generates MAF, effect size, and observed GWAS z-score at each genomic position, adapted from [15] to include binary traits. µ is the mutation rate, and k* ≡ 4*µ*/*E*[2*q*(1 − *q*)*s*] ≈ 1 *depends on f* (*s*) *and demography (see Supplementary Section 1*.*5 in [15]). h*^2^ *for a binary trait refers to heritability on the liability scale*. ***B)*** *Inferred distributions of selection coefficients estimated from 151 quantitative traits and 13 human complex diseases, respectively, with 90% confidence envelopes shown*. ***C)*** *Inferred distributions of selection coefficients when trait were instead grouped into 54 brain-related traits and 110 non-brain-related traits, with 90% confidence envelopes shown*. ***D)*** *Median estimates of heritability and mutational target size, inferred jointly with the respective f* (*s*) *shown in panel C. The gray lines are drawn at x* = 3 *×* 10^8^, *and y* = 0 *and* 1. *MCH: Mean corpuscular haemoglobin*.

First, we fit the extended Simons model to six complex diseases that have more than 100 approximately independent hits (trait-specific *f* (*s*) shown in Figure S20), as previous work suggests the inference of evolutionary parameters should be well powered in this case [15]. Two of the six diseases, schizophrenia and major depression, are brain-related. For comparison, we considered alternative models: fitting effect sizes with a normal distribution, and the *α*-model [21], which imposes a negative relationship between effect sizes and allele frequencies.

We quantified the goodness-of-fit of the different models by computing a residual p-value, defined previously as (Pr |*z*| > |*z*_*i*_||*z*_*i*_| > 5.45, *p*_*i*_, model) [15]. If we correctly model the distribution of z-scores among GWAS hits, then these residual p-values should be uniformly distributed between 0 and 1. To avoid overfitting, we split the genome into approximately independent blocks [53], each time inferring the model on 90% of the blocks and computing residual p-values for the held-out 10%. Across diseases, our pleiotropic stabilizing selection model provides a better fit to the residual p-values than the two simpler heuristic models (Figure S21). We next conducted Kolmogorov-Smirnov tests to determine whether the overall distribution of residual p-values for each trait matches the expected uniform distribution. At a false discovery rate of 0.05, we find that the simpler heuristic models provide a poor fit to the data; notably, we can reject the normal model for 5 of the 6 diseases, the *α*-model for 4, and the pleiotropic stabilizing selection model for none (Supplementary Table 4). These results confirm that the extended Simons model provides a robust fit for complex diseases and support the view that common variants affecting complex diseases are shaped by pleiotropic stabilizing selection [7, 33, 34].

To include traits with fewer hits in our inference, the Simons model further allows grouping traits to estimate a shared *f* (*s*) distribution, thereby reducing the number of trait-specific parameters from six to two — namely, *h*^2^/*L* and *L*. We therefore expanded our analysis to include all 151 quantitative traits and 13 diseases, each with more than 20 approximately independent hits. When stratifying traits into complex diseases and quantitative traits, the inferred distributions of selection coefficients have similar modes (Figure 4B).

By contrast, for traits with more than a hundred GWAS hits, the inferred trait-specific *f* (*s*) distributions differ markedly between brain-related and other complex traits (Figure S22). These findings motivated us to group traits into brain-related and non–brain-related categories. The two shared *f* (*s*) distributions reflect the stronger selective constraint acting on mutations affecting brain-related traits (Figure 4C). Moreover, the shared *f* (*s*) for brain-related traits is narrower than that for non–brain-related traits, which suggests a narrower distribution of effect sizes for brain-related traits, consistent with previous studies (see Introduction and [31, 32]).

We also found that brain-related traits tend to have substantially greater mutational target sizes than other complex traits (Figures 4D and S23), echoing previous studies that found brain-related traits to be more highly polygenic than other traits (see Introduction and [25, 28–30]). To contextualize these estimates, approximately 10% of the genome has been estimated to be constrained by selection and is therefore considered putatively functional [54–56]. For brain-related traits, our median inferred mutational target size is approximately 1.32% of the genome, compared with 0.27% for non-brain-related traits.

In contrast to selection and mutational target size, heritability estimates do not differ significantly between brain-related and other traits (Figures 4D and S23). In turn, estimates of *h*^2^/*L*, which is the key scaling parameter governing GWAS power and relating trait architectures within each category [15], are markedly lower for brain-related traits.

To validate our heritability estimates, we compared them with the recent whole-genome sequencing–based estimates 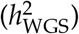 [57]. For the set of overlapping quantitative traits, our *h*^2^ estimates derived from brain-related and non-brain-related *f* (*s*) distributions show greater concordance with 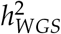 than estimates obtained under a single quantitative *f* (*s*) model, suggesting that grouping traits by CNS status improves model performance (Figure S24).

### Strong selection and large mutational target size explain the distinct architecture of brain-related traits

We next wanted to see if the inferred larger mutational target sizes and stronger selection on variants could explain the genetic architectures of brain-related traits. In particular, we used simulations to see the impact of mutational target size and selection on the MAF and |*z*|-score distribution of GWAS hits.

Under a fixed *f* (*s*) distribution — e.g., the one estimated from 151 quantitative traits — we simulated a trait with sample size 300*K* and trait heritability 0.5, varying only the mutational target size (Figure 5A). We found that larger values of *L* lead to steeper CDF curves for GWAS hit |*z*| -scores (Figure 5B). This occurs because GWAS power is linearly proportional to the scaling parameter *h*^2^/*L* (Figure 4A). The number of GWAS hits, on the other hand, scales nonlinearly with *L* (Figure 5C). While increasing *L* makes more genomic sites relevant to the trait [15], the power to discover each individual site declines. These opposing effects interact to produce the observed nonlinear trend. Nevertheless, the MAF distributions of GWAS hits are largely insensitive to changes in mutational target size, indicating that mutational target size variation cannot capture the observed enrichment of higher-MAF hits in brain-related traits (Figure S25).

**Figure 5:**
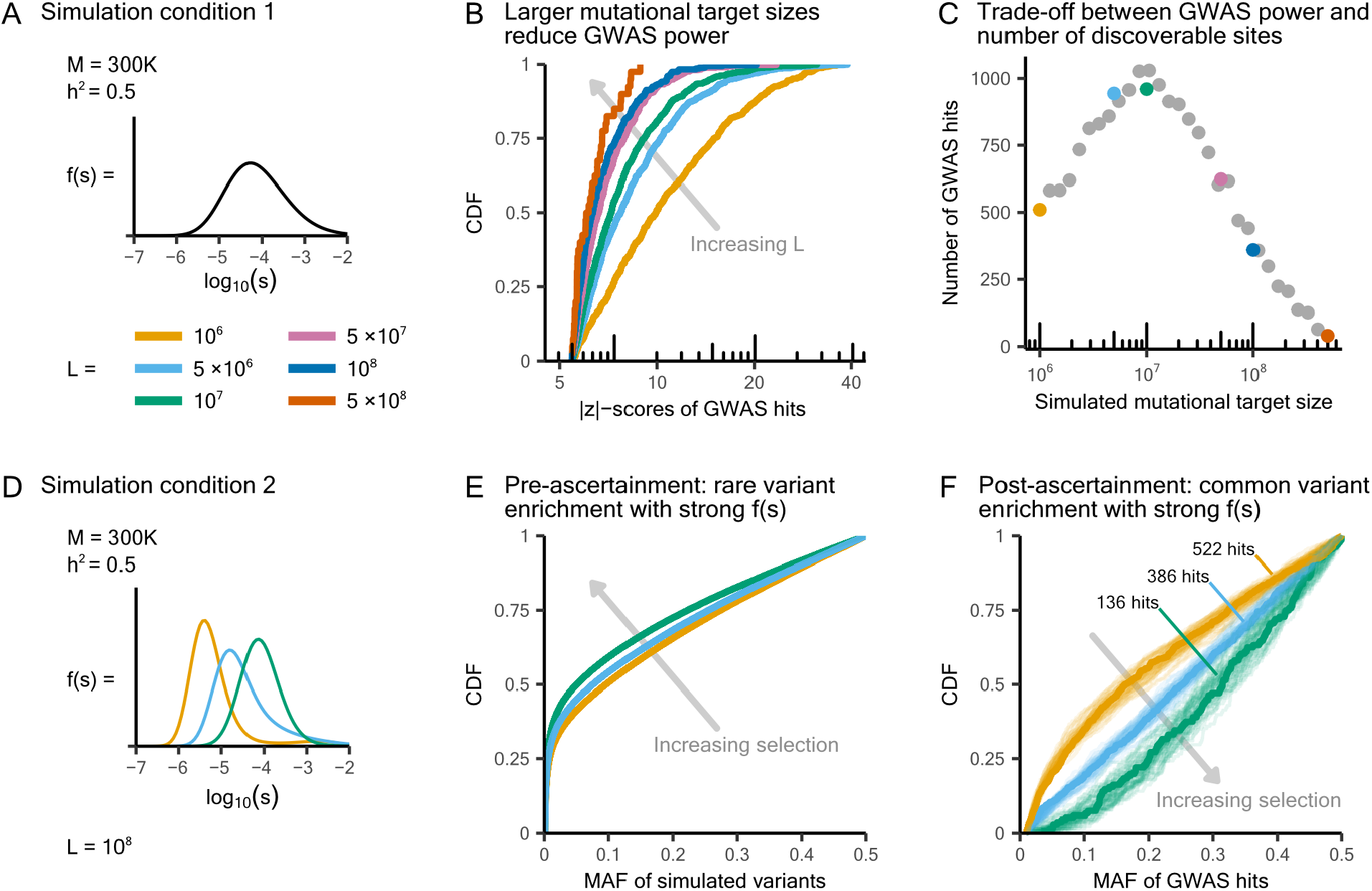
Recovering the genetic architectures of brain-related traits through simulations. ***A)*** *Simulation model for traits with varying mutational target sizes. We fixed f* (*s*) *to the shared distribution inferred from 151 quantitative traits as shown in Figure 4B. Colors in panels B and C correspond to the simulation settings shown in A*. ***B)*** *Larger L values lead to narrower distributions of GWAS hit* |*z*|*-scores*. ***C)*** *Nonlinear relationship between L and GWAS hit count. We ran additional simulations at finer increments of L and plotted them as gray dots*. ***D)*** *Simulation model for traits with varying distributions of selection coefficients. Colors in panels E and F correspond to the simulation settings shown in D*. ***E)*** *In the absence of GWAS ascertainment, variants simulated under stronger f* (*s*) *distributions tend to have lower MAF*. ***F)*** *After GWAS ascertainment (MAF* > 1% *and* |*Z*| > 5.45*), stronger f* (*s*) *simulations yield GWAS hits enriched for higher MAF. For each f* (*s*) *distribution, we ran 50 simulations, highlighted the median CDF curve of the MAF of simulated hits, and labeled the corresponding number of hits*.

We next sought to understand the impact of selection by holding the mutational target size at 10^8^ bp and varying the distribution of selection coefficients in our simulations (Figure 5D). In the absence of GWAS ascertainment, variants simulated under stronger *f* (*s*) distributions are generally rarer than those from weaker *f* (*s*), consistent with intuition (Figure 5E). Intriguingly, GWAS ascertainment — that is, restricting to common variants and selecting hits that surpass the genome-wide significance threshold — reverses this MAF relationship (Figure 5F). When trait-affecting variants are under stronger selection, only small effect variants can reach sufficiently high frequencies to be included in GWAS. Due to this ascertainment bias, an over-representation of small effect variants in turn leads to an enrichment of variants with higher MAF (Figure S26B). Additionally, stronger *f* (*s*) distributions also result in steeper |*z*| -score distributions and fewer hits (Figures S26 C to D).

### Rare variant burden tests suggest strong selection on brain-relevant genes

Although simulations show that a stronger *f* (*s*) distribution generates GWAS hits of higher MAF and lower |*z*| -scores — patterns consistent with what we observed for brain-related traits — these same GWAS hits are also the signals we used to infer *f* (*s*). To seek orthogonal evidence in support of stronger selective constraint on genetic variation influencing brain-related traits, we next turned to a genic perspective: are genes relevant to brain-related traits also under stronger selection compared to genes relevant to non-brain-related traits?

To that end, we leveraged previously reported estimates of the fitness effects of loss-of-function (LoF) variants, denoted *s*_*het*_ [58]. These estimates quantify selection against heterozygotes and serve as a metric of gene-level constraint. Moreover, from burden tests of these LoF variants, we obtained gene-to-trait effect estimates, 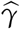, which we then used to derive unbiased estimates of the squared effects, 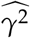 (Methods). One caveat to note is that our analysis did not include burden tests on binary traits, primarily due to limited data and the fact that statistical power is biased toward detecting risk-increasing effects in the rare variant regime.

Under a model of pleiotropic stabilizing selection, in which selection against LoF variants is mediated through their effects on traits, Spence *et al*. recently showed that on average 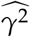 increases with increasing *s*_*het*_ across a set of independent quantitative traits [39]. In addition, multiple regression analyses showed that across 17 tissues, whether or not a gene is expressed in the “frontal cortex” is among the strongest predictors of estimated gene constraint, *s*_*het*_ (Figure S27). Together, these observations led to the hypothesis that brain-relevant genes are more constrained than other genes and that in constrained genes, the fitness effects of LoF variants should correlate more strongly with their effect on brain-related traits than on non-brain-related traits.

Binning genes by *s*_*het*_, we indeed found that genes that are more constrained have larger relative squared effect sizes, 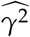, on brain-related than on non-brain-related quantitative traits, and vice versa for less constrained genes (Figure 6A). If these two trait categories had systematic differences in heritability or other factors, one curve would be consistently higher across the range. Instead, the two curves switch order, ruling out this possibility and supporting the hypothesis that effects on brain-related traits more strongly correlate with fitness consequences. In addition, we pruned the set of quantitative traits to 32 approximately uncorrelated traits, comprising 11 brain-related traits and 21 non-brain-related traits (Methods, Figure S28). Although the signal is noisier, we observed the same S-shaped trend (Figure S29).

**Figure 6:**
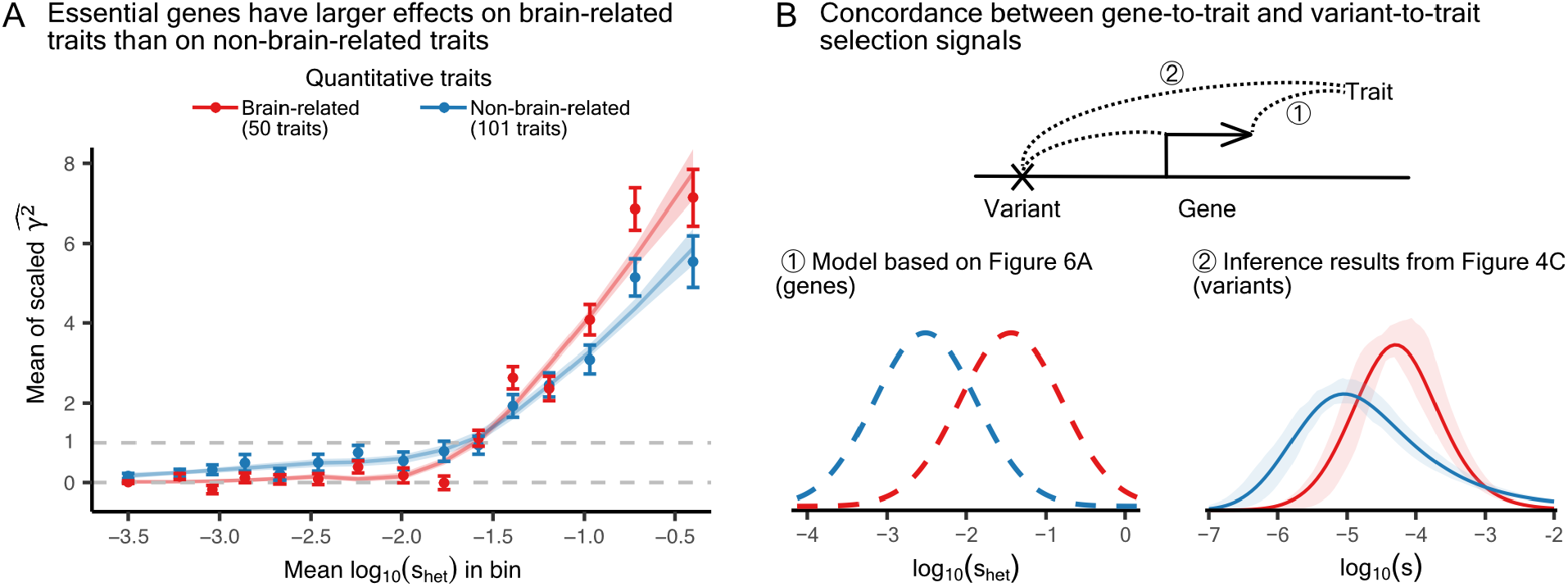
Burden tests provide aligned evidence of stronger selection on CNS-relevant genes. ***A)*** *Mean squared genic effects on brain-related versus non-brain-related quantitative traits, with genes grouped into 15 quantiles of s*_*het*_ *estimates [58]. The estimated* 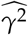 *values are scaled so that the mean across all gene-trait pairs in the tenth bin equals one. All estimates are shown with a* 95% *confidence interval computed from the standard error. We bootstrapped genes at the trait level and showed the mean and 95% confidence intervals of the fitted LOESS curves*. ***B)*** *Comparing selection signals from gene-to-trait and variant-to-trait associations*. ① *Brain-relevant genes have a stronger distribution of constraint values than genes relevant to other traits. The two curves are hypothetical and are intended to illustrate the relationship between their modes*. ② *Brain-relevant variants have a stronger distribution of selection coefficients than variants relevant to other traits*.

Figure 6A suggests a model in which brain-relevant genes have a stronger distribution of constraint values than genes relevant to non-brain-related traits (① of Figure 6B). If selection is stronger on changes in gene activity in the CNS relative to other tissues, this could help explain why we inferred a stronger distribution of selection coefficients for brain-relevant variants compared to variants relevant to other traits (Figure 4C, plotted again in ② of Figure 6B). Therefore, the strong selection signals we found for the brain-related traits are concordant at both the variant and gene levels.

## 3 Discussion

What are the processes that shape the genetic architecture of complex traits? Do certain groups of traits share common features, and if so, what causes them to collectively differ from other trait categories?

We found that brain-related quantitative traits and complex diseases share distinct genetic architectures, in that their GWAS hits have modest significance, narrow effect sizes, and are enriched at higher MAF. These features can be explained by the combination of brain-related traits generally having larger mutational target sizes than other complex traits and, importantly, experiencing markedly stronger selection on the genetic variation that affects them.

Our variant-level analyses were based on common variants, which are largely regulatory [59]. If selection on variants affecting brain-related traits is stronger than for other traits, then we would expect a greater proportion of their heritability to come from rare non-coding variants. In line with these expectations, recent studies using whole-genome sequencing data have found that brain-related traits exhibit relatively strong heritability enrichment from rare variants and the highest enrichment from non-coding variants across all traits examined [57].

As GWAS sample sizes continue to grow, we can use the inferred evolutionary parameters to explain past GWAS findings and predict future results. Focusing on the six complex diseases with more than a hundred GWAS hits, Figure 7A shows the simulated GWAS discoveries as a function of the effective study size (Equation 1). In particular, our findings for major depression recapitulate the historical trajectory: early studies showed a prolonged period with few or no significant findings [60–62], followed by a recent sharp increase in the number of detected loci [63–67]. Our inference provides a mechanistic explanation for these historical trends. To a good approximation, variants reach genome-wide significance when their contribution to phenotypic variance exceeds a threshold inversely proportional to the study’s sample size. Given the low heritability per site for major depression (Figure 7B), larger sample sizes are required to surpass this threshold, after which the number of discoveries sharply rises due to its large mutational target size (Figure 7C). More generally, we expect future GWAS discoveries for brain-related traits to increase more rapidly than those for non-brain-related traits, ultimately reaching higher saturation points.

**Figure 7:**
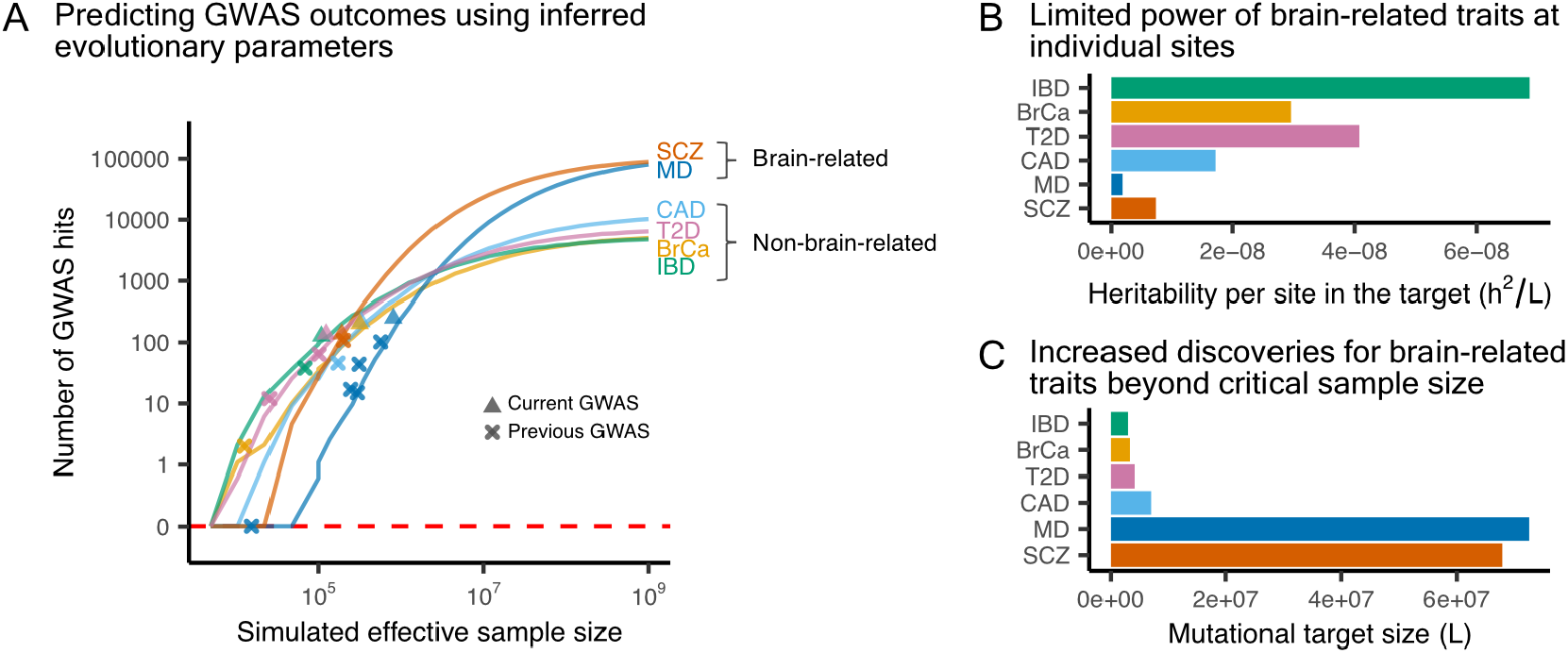
Explaining past and predicting future disease GWAS discoveries. ***A)*** *Expected disease GWAS outcomes as a function of sample size, computed using the inferred f* (*s*) *for brain-related and non-brain-related traits (Figure 4C), along with their corresponding estimates of h*^2^ *and L as shown in Figure 4D. We evaluated a range of sample sizes across 35 grid points and plotted the curves using the median values across 10 simulations at each point. Our inference was based on the current GWAS (see Supplementary Table 2). For reference, we overlaid the reported number of hits and sample sizes from selected earlier GWAS (IBD [73]; BrCa [74]; T2D [75, 76]; CAD [77]; MD [61, 63–66]; and SCZ [78])*. ***B)*** *Inferred h*^2^/*L estimates used in the simulations*. ***C)*** *Inferred L estimates used in the simulations*.

Importantly, our findings should not be interpreted as evidence that brain-related traits, especially the behavioral-cognitive traits, are themselves under stronger selection. Selection on brain-related variants and genes does not necessarily operate through the traits we considered. Rather, it can be posited that selection acts primarily on core aspects of brain development and functioning, more strongly than on other tissues. In support of this view, among constrained genes depleted of missense and LoF variants, 30% are highly expressed in the brain, compared with only 14% in lymphocytes, which rank second across tissues [68]. However, these processes are difficult to define as traits, let alone to quantify or measure. The brain-related traits we analyzed therefore serve only as proxies, tagging the variants and genes involved in these essential biological processes.

Additionally, it is within the Simons modeling framework that our findings provide the most plausible explanation for the genetic architectures of brain-related traits. We also examined potential ways in which brain-related traits might violate the modeling assumptions. For example, since brain-related traits are highly correlated [69], they could in principle collapse onto a single underlying trait, thereby undermining the assumption of a high-dimensional trait space. However, simulations assuming no pleiotropic variant effects cannot recover the observed MAF distributions of GWAS hits for brain-related traits (Methods, Figure S30).

Why would selection on genetic variation be stronger for brain-related traits compared to traits mediated by other tissues? Notably, brain tissues are distinct in that expression quantitative trait loci (eQTL) effects show high levels of sharing within the brain, yet remain largely unshared with non-brain tissues [70]. This suggests that variants regulating gene expression in the brain may be largely brain-specific. With most common variants being regulatory, we can envision the strong selection happening in three ways, where comparing CNS to other tissues: i) variants could typically have greater effects on the activity (e.g., expression) of genes; ii) changes in the activity of genes could typically have greater effects on traits; and iii) changes in the activity of genes typically affect more selected traits. The genic analyses we presented (Figure 6) suggest that changes to gene expression involve greater fitness effects in the CNS than in other tissues, which in turn supports the latter two models (without excluding the first).

Furthermore, our analysis suggests that traits that are mediated by the same tissues tend to have similar architectures and thus similar evolutionary parameters. Brain-related traits appear to occupy a “corner” in the architecture space, which is why their genetic architectures are more clearly set apart from the architecture of traits mediated by other tissues, but we also see a clustering of architectures for cardiovascular and skeletal muscle traits, for example (Figure S9 and S10). The omnigenic model [71, 72] suggests why trait architectures might cluster by tissues. If many of the regulatory variants that affect gene activity in given cell types contribute to heritable variation in traits that are mediated by these cell types, then we would expect these traits to share similar mutational target sizes and distributions of selection effects; the sharing of effect size distributions (scaled appropriately) follows from the shared distribution of selection effects under the pleiotropic stabilizing selection model [14]. Nonetheless, our inference suggests that there is substantial variation in mutational target sizes and distribution of selection effects among brain-related traits (Figures S22 and S23). This too is to be expected: even setting aside the coarseness in associating traits with tissues and the uncertainty in our inferences, traits that are mediated by the same tissue are expected to have different numbers of “core genes” and to be differentially affected by different cellular circuits.

The case of body composition-related traits is especially interesting from this point of view. As noted above, these traits are mediated by a mixture of tissues, including the CNS (Figure S11). Their architectures and inferred evolutionary parameters appear to fall between those of brain-related traits mediated primarily by the CNS and those of non-brain-related traits (Figures 4D and S23). More generally, this suggests that the architecture of common variation underlying complex traits can be approximated as a superposition of architectures associated with tissues, with some trait-specific effects in each tissue.

## 4 Methods

### Identifying approximately independent hits using GCTA-COJO

From GWAS summary statistics, we applied GCTA-COJO [38] to identify conditionally independent hits and re-estimated variant effect sizes 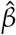 on each trait. We used parameters `–cojo-p 5e-8` and `–cojo-slct`, along with a LD reference panel consisting of genotypes from a subset of 10,000 White British individuals from the UK Biobank. We removed hits with a minor allele frequency (MAF) below 1%, those located in the HLA region, and those with INFO scores below 0.8.

### Data on 151 quantitative traits

#### GWAS summary statistics

We downloaded GWAS summary statistics for 305 quantitative phenotypes (Supplementary Table 1; same trait list as in [39]) from the Neale Lab (http://www.nealelab.is/uk-biobank, version 3). Specifically, these results were based on 361,194 individuals of White British ancestry in the UK Biobank. All the phenotypes were rank-based inverse normal transformed. Age, age^2^, *×* inferred sex, age×inferred sex, age^2^ *×* inferred sex, and principal components 1 to 20 were included as covariates.

#### SNP heritability

Estimates for the 305 quantitative traits were downloaded from the Neale Lab.

#### Number of significant GWAS hits

We followed the same procedure as described in the subsection for identifying approximately independent hits.

For this study, we focused on 151 quantitative traits, each with at least 20 significant hits and a SNP *h*^2^ greater than 5% (Supplementary Table 1).

### Data on 13 complex diseases

#### GWAS summary statistics

Since the UK Biobank cohort has been found to be healthier than the general population [79], we turned to publicly available latest GWAS summary statistics from case-control studies. We restricted our analyses to studies on cohorts of European ancestry, as the demographic model in [15] was specifically derived for the European population. For several diseases, the complete results require additional approval from 23andMe. We thus used only the publicly available version, excluding the 23andMe cohort.

#### Number of significant GWAS hits

We followed the same procedure as described in the subsection for identifying approximately independent hits.

Using the same cut-off of more than 20 genome-wide independent hits, we curated a list of 13 complex diseases (Supplementary Table 2). Although not exhaustive, this list defines the scope of this present study.

### Stratified LD score regression (S-LDSC)

GWAS summary statistics on all 164 traits were processed by the `“munge_sumstats.py”` script provided by LDSC developers (https://github.com/bulik/ldsc, v1.0.1) [80]. LD scores were computed based on European ancestry participants in the 1000 Genomes Phase 3 dataset [81]. To identify key cell types for each trait, we conducted cell type-specific analyses with S-LDSC [26]. Finucane et al. categorized 220 distinct cell types into 10 categories: *adrenal / pancreas, cardiovascular, central nervous system (CNS), connective / bone, gastrointestinal, immune / hematopoietic, kidney, liver, other*, and *skeletal muscle*. Each cell-type-specific annotation was then added to the baseline LD model (v1.2) to evaluate heritability enrichment. The resulting regression coefficient z-scores were converted to one-sided p-values, shown in Figures 2B and S4 to S6. To account for multiple testing, we applied a Bonferroni correction, setting the significance threshold to a p-value of 0.05/(220 *×* 164).

In Figures 2D, S9, and S10, the p-values for each cell-type category is obtained from metaanalyses using the aggregated Cauchy association test (ACAT), which is robust to the correlations among the constituent p-values [41]. Specifically, within each category, p-values were transformed into Cauchy variables and aggregated through 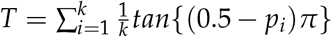. The resulting p-value can be obtained from the Cauchy distribution with 0.5 − *{arctan*(*T*)*}*/*π*.

### Thresholding and downsampling quantitative traits

For a given quantitative trait, we obtained recorded trait values from approximately 360K unrelated White British individuals in the UK Biobank. We first conducted GWAS on rank-based inverse normal transformed phenotypes using PLINK 2.0 [82], with parameters `–glm hide-covar` and `–covar-variance-standardize`. Age, age^2^, sex, age *×* sex, array, center and principal components 1 to 20 were included as covariates in the regression model. We then identified conditionally independent hits using GCTA-COJO as described above.

For the thresholding analysis, we dichotomized the continuous trait into ten binary traits (cases=2 and controls=1 following PLINK format), representing a disease prevalence of 1%, 2%, 5%, 10% and 20% on either the lower or the upper tail of the phenotypic distribution. To reduce computational burden, we ran case-control GWAS only on the list of previously identified hits with flag `–extract` and modified parameter `–glm firth-fallback hide-covar`.

For the downsampling analysis, we drew five subsamples, each with sample sizes calculated to match with the prevalence rates of 20%, 10%, 5%, 2% and 1% used to create previous binary traits (Supplementary notes). Smaller subsets were ensured to nest within larger ones. Phenotypes in each subset were again rank-based inverse normal transformed to run GWAS using the same variant list filter.

Lastly, z-scores from each of the ten thresholded or five downsampled GWAS were linearly regressed on z-scores from the original GWAS.

These procedures were repeated for six medically relevant traits, as they are commonly used in clinical settings to declare disease status based on specific cutoffs. These traits include LDL, BMI, hemoglobin A1c (HbA1c), diastolic blood pressure, systolic blood pressure, and heel bone mineral density T-score. The corresponding UK Biobank data fields are 30780, 21001, 30750, 4079, 4080, and 78, respectively.

### Estimating *N*^*′*^ for binary traits and re-scaling GWAS hit z-scores

For a given complex disease, we defined in Eq. 1 that 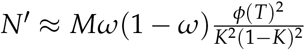 is the sample size needed on the underlying liability scale to reach equivalent power as of the original case-control study. Let *p*_*i*_ be the MAF of SNP *i* and 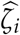 be its estimated effect size on the log odds ratio scale. Given that 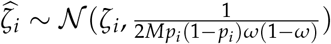 (Supplementary notes), we estimated 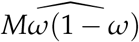 with the median value of 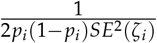 across all SNPs in the summary statistics file. Disease prevalence values, *K*, were taken from the GWAS source paper when available. Otherwise, we used prevalence estimates in the European population. *ϕ*(*T*) is the density at the liability threshold, assuming that the underlying liability follows a standard normal distribution.

When standardizing two traits to have comparable sample sizes, regardless of whether the trait is quantitative with reported sample size or binary with estimated sample size, we simply scale the GWAS hit z-scores of the trait with larger sample size with 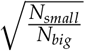.

### Inferring *f* (*s*), *h*^2^, and *L*; and simulating GWAS hits

We used the scripts developed by Simons *et al*. for analyzing quantitative traits [15]. The complete inference framework is described in detail in their Supplement. Here, we outline the relevant key concepts. With the significant GWAS hits for a given set of traits, we used a maximum likelihood approach to jointly infer a shared *f* (*s*) across traits, along with trait-specific parameters, *h*^2^ and *L*. The log likelihood of observing a GWAS hit *i* is

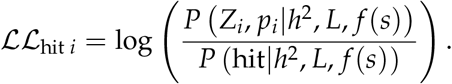

By the chain rule in probability, the numerator is *P*(*Z*_*i*_|*p*_*i*_, *h*^2^, *L, s*)*P*(*p*_*i*_|*s*). The site frequency spectra, *P*(*p*_*i*_|*s*), are estimated with forward simulations assuming a European demographic model. The z-scores then follow 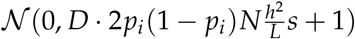 up to a constant term *D*. The denominator accounts for GWAS ascertainment and is a double integral of the numerator for |*z*| > 5.45 and *p* > 1%. Details can be found in Supplementary Section 3.3 of [15].

For a given *f* (*s*) distribution, we first find each trait *j*’s maximizing 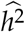 and 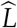 for

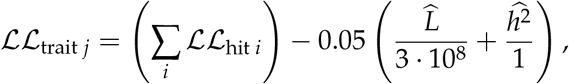

where the minus term penalizes estimates beyond reasonable thresholds. The shared *f* (*s*) across traits then maximizes (∑_*j*_ ℒℒ_trait *j*_). Confidence intervals are obtained by bootstrapping and resampling genomic blocks. Details can be found in Supplementary Sections 2.8, 3.5, and 3.7 to 3.9 of [15].

To extend the inference framework to accommodate binary traits, we simply transferred effect sizes from log odds ratio scale to the liability scale with Eq. 6. For binary traits, *h*^2^ denotes heritability on the liability scale, and *N* represents the corresponding effective sample size on that scale, *N*^*′*^.

We further adopted the scripts provided by Simons *et al*. [15] to simulate GWAS hits of a trait, with input parameters *f* (*s*), *h*^2^, and *L*. Briefly, we simulated *L* variants with MAF and z-scores drawn from *P*(*p*_*i*_|*s*) and *P*(*Z*_*i*_|*p*_*i*_, *h*^2^, *L, s*) and outputted the variants with *p*_*i*_ > 1% and |*Z*| > 5.45. Details can be found in Supplementary Section 5.1 of [15].

To modify the Simons model into a one-dimensional trait space model, i.e., assuming that variants do not have pleiotropic effects, we followed results in Simons et al. [14]. Contrary to a high-dimensional trait space in which 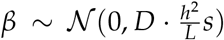, in a one-dimensional trait space *β* equals 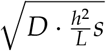. It follows that the z-scores for a given variant should instead be drawn from 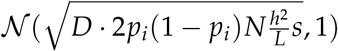.

### LoF burden tests on 151 quantitative traits

Summary statistics for LoF burden tests on 112 quantitative traits were downloaded from Milind *et al*. [83]. For the remaining 39 quantitative traits, we followed the scripts they provided and conducted gene-based burden tests with REGENIE [84]. A whole genome regression model is fit in the first step to make Leave One Chromosome Out (LOCO) phenotypic predictions. We filtered variants for MAF > 1%, missingness < 10%, Hardy-Weinberg equilibrium test p-value < 10^−15^ and applied linkage disequilibrium pruning with 1000 variant windows, 100 variant shifts, and *r*^2^ < 0.9. In the second step of REGENIE, LoF burden tests were performed with LOCO predictions included as an offset term. We filtered LoF variants to have MAF < 1% and misannotation probability (estimated in [58]) < 10%. At both stages, we applied rank-based inverse normal transformation to all phenotypes and included age, age^2^, sex, age *×* sex, genotyping PCs 1 to 15, rare variant PCs 1 to 15, and WES batch as covariates. With the estimated gene-to-trait burden effects, 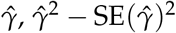 is an unbiased estimator of mean squared genic effects *γ*^2^ [39, 83].

### Distinct subsets of (approximately uncorrelated) brain-related and non-brain-related Traits

From the set of 164 complex traits, 50 quantitative traits and 4 complex diseases with meta-analysis p-values for CNS enrichment passing the Bonferroni threshold of 0.05/(10 *×* 164) are classified as CNS traits. The remaining 101 quantitative traits and 9 complex diseases are classified as non-CNS traits (Supplementary Table 1).

To reduce redundancy in the trait list, we pruned the set of 151 quantitative traits to 32 approximately uncorrelated traits as shown in Figure S28. Specifically, traits were sorted in decreasing order of meta-analysis p-values for CNS enrichment. Starting with the trait that had the strongest enrichment for CNS, *time to complete round*, we iteratively added traits whose pairwise genetic correlations with all previously selected traits did not exceed 0.5. Genetic correlation estimates were obtained from the Neale Lab. We excluded 27 biomarker traits that did not have genetic correlation estimates. In the end, we selected 32 approximately uncorrelated traits, of which 11 are CNS traits (Supplementary Table 1).

### Code availability

Codes used for this article are available at https://github.com/huishengz/cns-selection. Data and results are available at https://doi.org/10.5281/zenodo.19154957.

### URLs

PLINK 2.0 software: https://www.cog-genomics.org/plink/2.0/

Neale lab UKB data: http://www.nealelab.is/uk-biobank

LDSC software: https://github.com/bulik/ldsc

## Acknowledgments

We thank Alvina Adimoelja, Tami Gjorgjieva, Nikhil Milind, Roshni A. Patel, and members of the Pritchard Lab for helpful discussions and feedback. This research has been conducted using the UK Biobank resource under application number 52374. Analyses involving UK Biobank individual level data were conducted on the Research Analysis Platform (https://ukbiobank.dnanexus.com). We gratefully acknowledge the Psychiatric Genomics Consortium (PGC), the European Alzheimer & Dementia Biobank (EADB) consortium, the Breast Cancer Association Consortium (BCAC), the CARDIoGRAMplusC4D Consortium, the International Inflammatory Bowel Disease Consortium (IIBDGC), the International Multiple Sclerosis Genetics Consortium (IMSGC), the International Parkinson’s Disease Genomics Consortium (IPDGC), the Genetic Associations and Mechanisms in Oncology (GAME-ON) / Elucidating Loci Involved in Prostate Cancer Susceptibility (ELLIPSE) Consortium, and other research consortia for access to GWAS summary statistics files used in this study. H.Z. was supported by Stanford Biology Department’s graduate student assistantship. Y.B.S. was supported by grants from the National Science Foundation (DMS-2235451) and Simons Foundation (MPS-NITMB-00005320) to the NSF-Simons National Institute for Theory and Mathematics in Biology (NITMB). This work was supported by the National Institutes of Health grants R01HG014005 and R01HG008140 to J.K.P., R01GM115889 to G.S., and R01HL175076 to J.P.S..

## Author contributions

**H. Z**. Conceptualization, methodology, software, analyses, visualization, writing. **Y. B. S**. Conceptualization, methodology, software, writing, mentorship. **J. P. S**. Conceptualization, methodology, writing, mentorship. **G. S**. Conceptualization, methodology, writing, supervision, funding acquisition. **J. K. P**. Conceptualization, methodology, writing, supervision, funding acquisition.

## Supplementary Materials

### Supplementary Notes GWAS power

#### Quantitative trait GWAS power

In a linear regression model of the form *Y* ∼ *Xβ* + *ϵ*, the estimated effect sizes are approximately distributed as 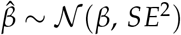 [85]. For a quantitative trait GWAS, *Y* is a vector of standardized phenotype values, *X* is a vector of genotype values at a focal SNP, *β* is the additive effect size per copy of the effect allele, and the residual term *ϵ* ∼ *N*(0, *σ*^2^) captures the background genetic and environmental noise. If *H*_1_ is true, i.e., *β≠* 0, *Z*^2^ approximately follows a non-central chi-square distribution, 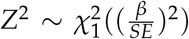. GWAS power is determined by this non-centrality parameter 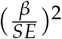. Furthermore, if *N* is the total number of individuals in the study, and *X*_*i*_ is centered so that 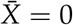,

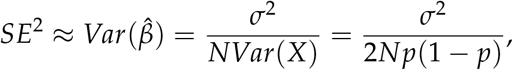

where the last step assumes Hardy-Weinberg equilibrium at this locus. Assuming *σ*^2^ is approximately the phenotypic variance, *V*_*p*_, and the trait is standardized to have unit variance, we get

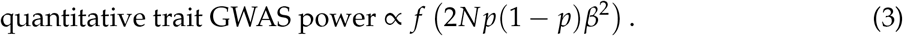

#### Binary trait GWAS power

A case-control GWAS typically assumes a logistic regression model of the form

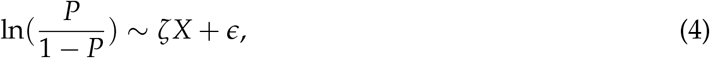

where *P* = *Prob*(*Y* = 1), and *ζ* is the effect size of allele 1 measured on the log odds ratio scale.

Given a contingency table as shown in Table 1 and assuming that all cell counts are non-zero,

**Table 1:**
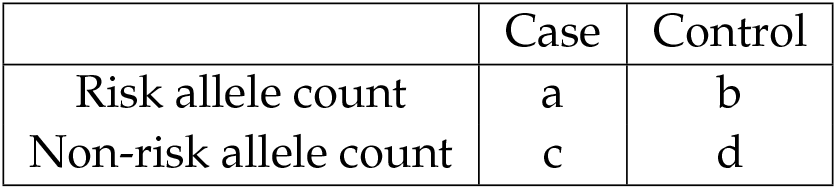
Contingency table.

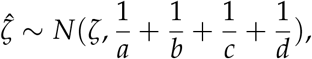

approximately. Under the null hypothesis that *ζ* = 0, we have *E*(*a*) = 2*Mωp* given that *M* is the number of cases plus the number of controls, and *ω* is the sample prevalence. Similarly, *E*(*b*) = 2*M*(1 − *ω*)*p, E*(*c*) = 2*Mω*(1 − *p*), and *E*(*d*) = 2*M*(1 − *ω*)(1 − *p*). Plugging these expected values into the previous formula, it follows that 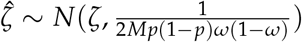. Therefore, we arrive at

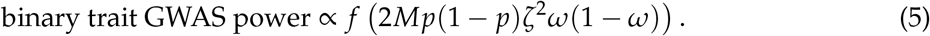

### Relating effect sizes

Note that the effect sizes obtained from a quantitative trait GWAS, *β*’s, are measured in standardized units on an additive scale, whereas the effect sizes obtained from a binary trait GWAS, *ζ*’s, are measured on a log odds ratio scale. Previous studies found that if effect sizes are sufficiently small, then the relationship between these two effect sizes is

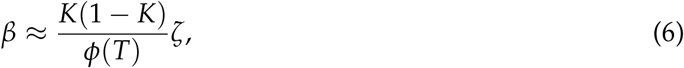

where *K* is the prevalence of the disease in the population and *ϕ*(*T*) is the standard normal density at the liability threshold [11, 86].

Here, we provide a derivation of this relationship. In a population carrying a variant with effect size *β*, the disease liability distribution is shifted to the right compared with a population without the variant (Figure S1). When the effect size, *ζ*, is sufficiently small, using Equation 4 we find that

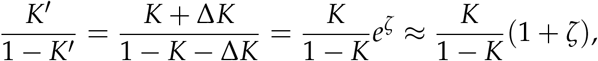

which implies that

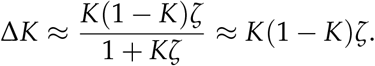

In turn, when *β* is sufficiently small, the increase in prevalence can be approximated by

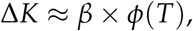

where *ϕ*(*T*) is the density at the liability threshold. The relationship between effect sizes on the liability scale and on the log odds ratio scale (Equation 6) follows from these expressions. When effect sizes are sufficiently large, this approximation breaks down, and Δ*K* depends on the shape of the liability distribution, which is typically assumed to be standard normal.

**Figure S1:**
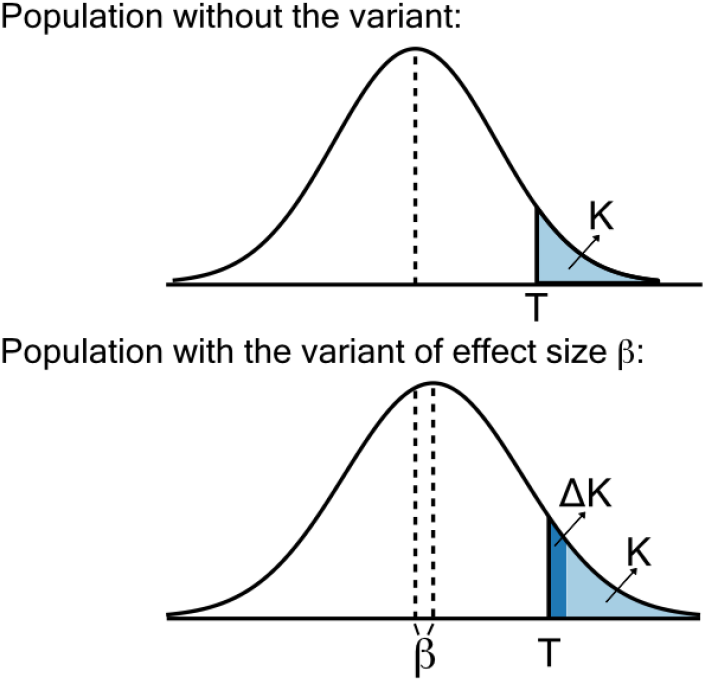
Relating effect sizes with the liability threshold model. Our derivation here assumes that effect sizes (both *β* and *ζ*) are small.

### Binarizing and downsampling a quantitative trait

Plugging Equation 6 into Equation 5, we can re-write binary trait GWAS power as

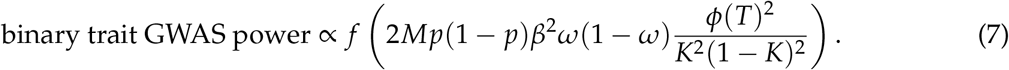

To better understand how to relate the GWAS power of a quantitative trait and a binary trait, we next binarized a quantitative trait and ran a case-control GWAS. In the simplest case, we sampled the full set of cases and controls after binarization, that is, *N* = *M* and *K* = *ω*. According to Equation 7 and Equation 3, we see that binarizing a quantitative trait reduces GWAS power, with the case-control GWAS retaining 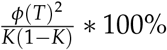 of the original GWAS power (Figure S2).

**Figure S2:**
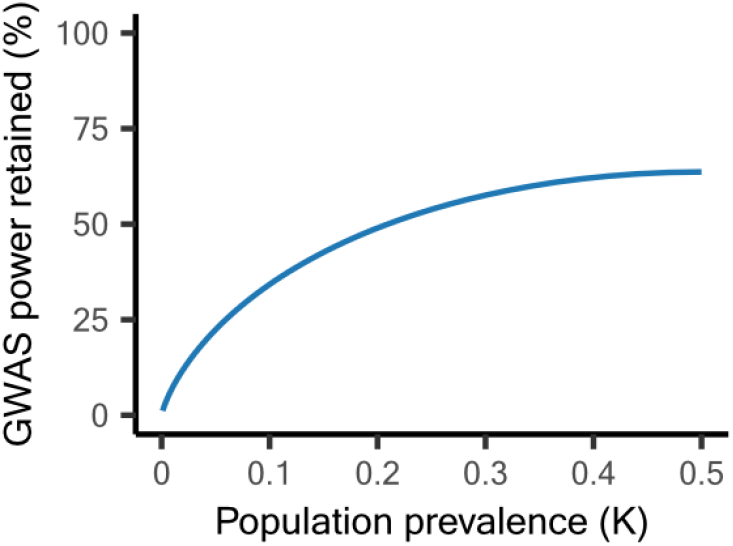
Reduction in GWAS power after converting a quantitative trait to a binary trait. We assume that the sample size remains the same and that the population prevalence matches the sample prevalence. *ϕ*(*T*) denotes the density at the liablity threshold, assuming liability follows a standard normal distribution.

Furthermore, the loss in GWAS power after trait binarization can be replicated by down-sampling the original trait to a new sample size *N*^*′*^, given by Equation 1. The obtained Z-scores after binarization and downsampling both will be deflated by a factor of

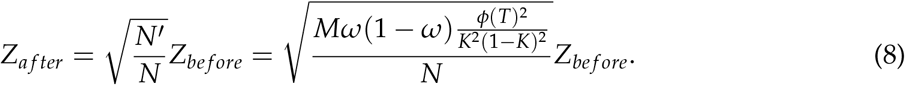

Equation 8 gives the predicted slopes, shown as solid colorful lines in Figures 2C and D.

### Insights from the formula for estimating *N′*

The formula we derived for transferring the sample size of a case-control GWAS to the effective sample size on the liability scale, 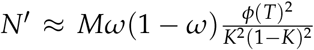, can be decomposed into two parts. From the first factor, *ω*(1 − *ω*), it is evident that the power of a case-control GWAS is maximized when the sample prevalence is 50%. The second factor, 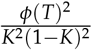, is the main reason why the estimated *N*^*′*^ for the current schizophrenia GWAS, 192,273, is larger than its original sample size of *M* = 130,644 (Methods, Supplementary Table 3). For a given case-control study, i.e., when *Mω*(1 − *ω*) is fixed, a higher statistical power is achieved when the disease is less prevalent (Figure S3). One can also regard this as rewarding the extra data collection effort to recruit the same number of cases for a rarer disease.

Combining these elements, we arrived at a strategy for optimizing GWAS design. Suppose we want to conduct a GWAS on a quantitative trait, but with a constraint on budget. In other words, there is a limit on the number of individuals we can genotype or sequence. Under these circumstances, the optimal approach that maximizes GWAS power is to sample *X* individuals in the upper *K*% of the phenotypic distribution, match with an additional *X* individuals sampled from the lower (1 − *K*)% of the distribution, and conduct a case-control GWAS. As long as *K* is below 8.5%, the second factor will be larger than 4 (Figure S3), and therefore the resulting case-control GWAS will have greater statistical power than conducting a quantitative trait GWAS with 2*X* individuals.

**Figure S3:**
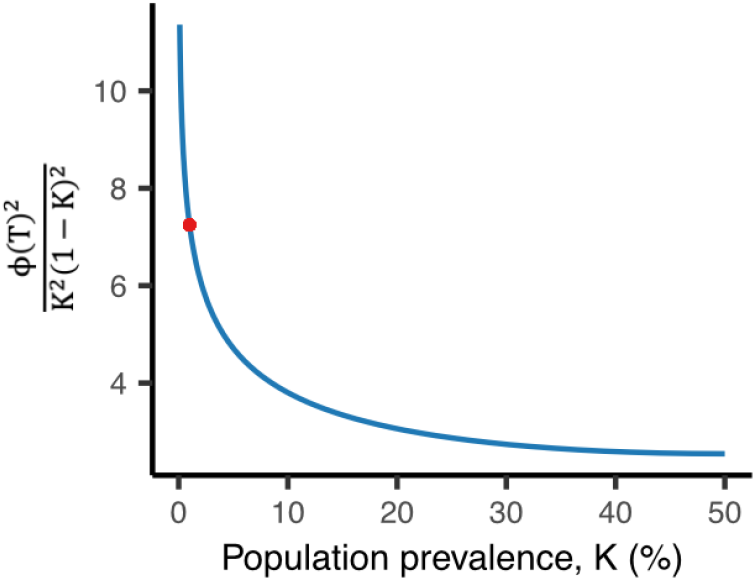
A larger sample size is needed on the liability scale when the disease is rarer. The red point is placed at *K* = 1%, the assumed population prevalence for schizophrenia [37].

## Supplementary Tables

Supplementary Table 1: **Details of quantitative traits included in this study.**

**Supplementary Table 2:** Details of complex diseases included in this study. Table of 13 complex diseases included in this study, with commonly used abbreviations, source of case-control GWAS summary statistics, and the number of genome-wide independent associations identified with GCTA-COJO [38] (Methods).

| Disease | Abbreviation | GWAS source | Number of hits |
| --- | --- | --- | --- |
| Alzheimer's disease | AD | [87] | 62 |
| Attention deficit hyperactivity disorder | ADHD | [88] | 24 |
| Bipolar disorder | BIP | [89] | 74 |
| Breast cancer | BrCa | [90] | 202 |
| Coronary artery disease | CAD | [36] | 217 |
| Inflammatory bowel disease | IBD | [91] | 127 |
| Major depression | MD | [67] | 249 |
| Multiple sclerosis | MS | [92] | 87 |
| Parkinson's disease | PD | [93] | 22 |
| Prostate cancer | PrCa | [94] | 134 |
| Rheumatoid arthritis | RA | [95] | 97 |
| Schizophrenia | SCZ | [37] | 196 |
| Type 2 Diabetes | T2D | [96] | 138 |

**Supplementary Table 3:** Key parameters for complex diseases included in this study. *M*: total sample size in the case-control study, including both cases and controls; *ω*: sample prevalence; *K*: population prevalence commonly used or estimated for the European population, taken from the GWAS source paper unless annotated with *; *N*^*′*^: effective sample size required on the liability scale to reach equivalent power, calculated with Equation 1.

| Disease | Reported $M$ | Reported $\omega$ | $M\omega(1 - \omega)$ | $\widehat{M\omega(1 - \omega)}$ | $K$ | $N'$ |
| --- | --- | --- | --- | --- | --- | --- |
| AD | 487,511 | 17.6% | 70,701 | 29,699 | 5% [97]* | 140,016 |
| ADHD | 225,543 | 17.2% | 32,121 | 22,449 | 5% [88] | 105,835 |
| BIP | 840,309 | 7.1% | 55,426 | 35,255 | 2% [89] | 215,143 |
| BrCa | 247,173 | 54.0% | 61,398 | 52,115 | 2% [98]* | 318,030 |
| CAD | 1,165,690 | 15.6% | 153,479 | 78,272 | 7% [36] | 332,959 |
| IBD | 59,957 | 41.8% | 14,586 | 12,836 | 0.5% [73]* | 108,431 |
| MD | 2,000,702 | 20.6% | 327,243 | 244,161 | 15% [67] | 816,506 |
| MS | 115,803 | 41.0% | 28,013 | 7,327 | 0.1% [92] | 83,236 |
| PD | 482,730 | 7.0% | 31,426 | 4,018 | 0.5% [93] | 33,945 |
| PrCa | 140,254 | 56.4% | 34,489 | 29,087 | 2% [98]* | 177,504 |
| RA | 97,173 | 23.0% | 17,209 | 8,159 | 0.5% [95] | 68,921 |
| SCZ | 130,644 | 40.9% | 31,579 | 26,529 | 1% [37] | 192,273 |
| T2D | 659,316 | 9.5% | 56,685 | 32,547 | 10% [96] | 123,757 |

**Supplementary Table 4:** Evaluating model fit of the normal distribution, the alpha model, and the Simons model. For each trait-model combination, we conducted Kolmogorov–Smirnov (KS) tests to assess whether the residual p-values follow a uniform distribution. We then applied the Benjamini-Hochberg (BH) procedure to the KS p-values for each model to control the false discovery rate at 0.05.

|  | **Normal distribution** |  | **Alpha model** |  | **Trait-specific f(s)** |  |
| --- | --- | --- | --- | --- | --- | --- |
| **Disease** | **KS p-value** | **BH decision** | **KS p-value** | **BH decision** | **KS p-value** | **BH decision** |
| BrCa | $9.521 \times 10^{-10}$ | **Reject** | $4.666 \times 10^{-8}$ | **Reject** | 0.641 | Accept |
| CAD | $1.756 \times 10^{-12}$ | **Reject** | $6.384 \times 10^{-8}$ | **Reject** | 0.242 | Accept |
| IBD | $3.992 \times 10^{-3}$ | **Reject** | 0.201 | Accept | 0.904 | Accept |
| MD | 0.765 | Accept | 0.764 | Accept | 0.538 | Accept |
| SCZ | $6.797 \times 10^{-3}$ | **Reject** | 0.0164 | **Reject** | 0.0498 | Accept |
| T2D | $5.133 \times 10^{-9}$ | **Reject** | $1.405 \times 10^{-9}$ | **Reject** | 0.191 | Accept |

**<sup>1</sup> Supplementary Figures.**

## Supplementary Figures

**Figure S4:**
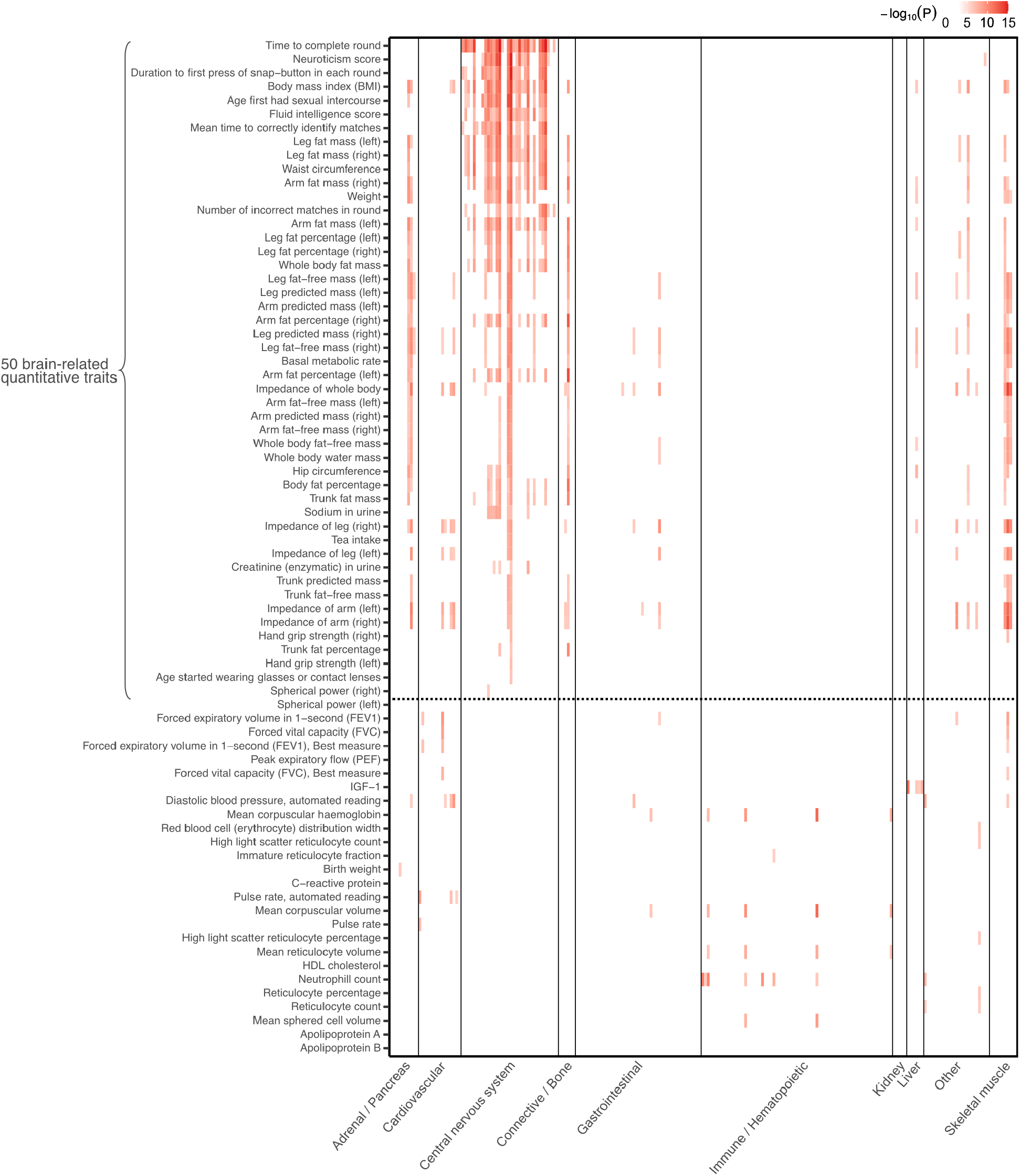
S-LDSC functional enrichment analyses on 151 quantitative traits (first half). *Each row consists of 220 cell types from 10 categories, colored by* −*log*_10_(*p*) *for the coefficient τ in the S-LDSC model [26, 27] if* −*log*_10_(*p*) *passes the Bonferroni-corrected threshold* −*log*_10_(0.05/(220 *×* 164)). *Traits are ordered by decreasing significance of the meta-analyzed p-values for CNS enrichment (Methods). The top 50 traits pass the significance threshold of* −*log*_10_(0.05/(10 *×* 164)) *and are therefore classified as brain-related*.

**Figure S5:**
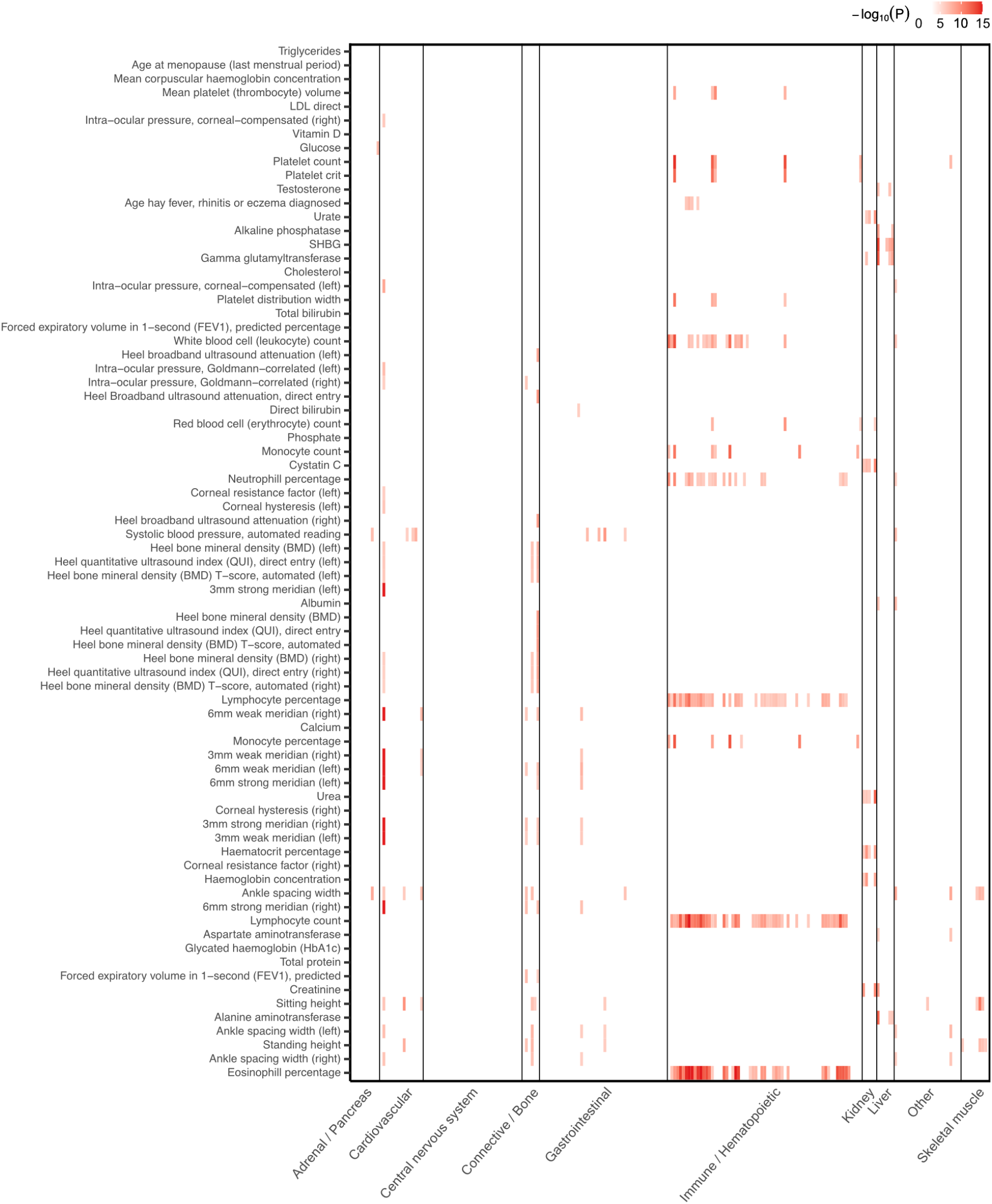
S-LDSC functional enrichment analyses on 151 quantitative traits (second half). *Each row consists of 220 cell types from 10 categories, colored by* −*log*_10_(*p*) *for the coefficient τ in the S-LDSC model [26, 27] if* −*log*_10_(*p*) *passes the Bonferroni-corrected threshold* −*log*_10_(0.05/(220 *×* 164)). *Traits are ordered by decreasing significance of the meta-analyzed p-values for CNS enrichment (Methods). All traits in this figure are classified as non-brain-related*.

**Figure S6:**
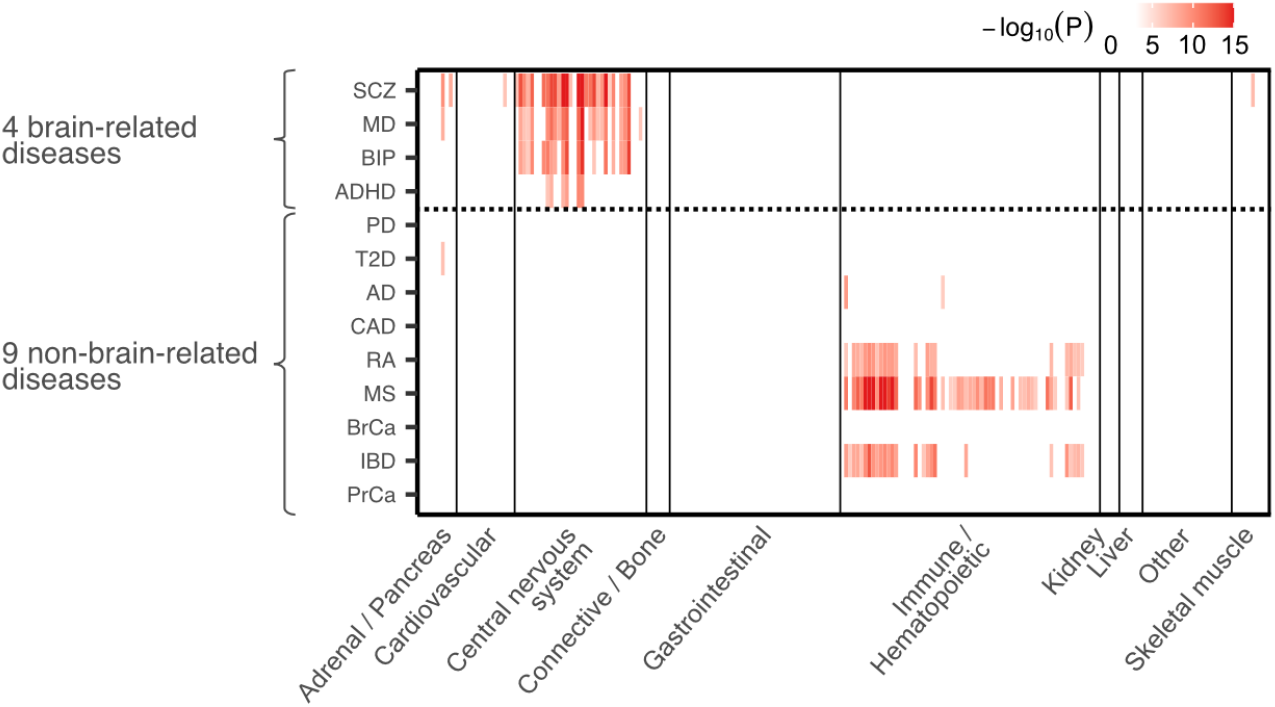
Functional enrichment analyses with S-LDSC on 13 human complex diseases. *Each row consists of 220 cell types from 10 categories, colored by* −*log*_10_(*p*) *of the coefficient τ in the S-LDSC model [26, 27] if* −*log*_10_(*p*) *passes the Bonferroni correction threshold* −*log*_10_(0.05/(220 *×* 164)). *Traits are ordered by decreasing significance of the meta-analyzed p-values for CNS enrichment (Methods). The top 4 traits pass the significance threshold of* −*log*_10_(0.05/(10 *×* 164)) *and are therefore classified as brain-related. Disease full names can be found in Supplementary Table 2*.

**Figure S7:**
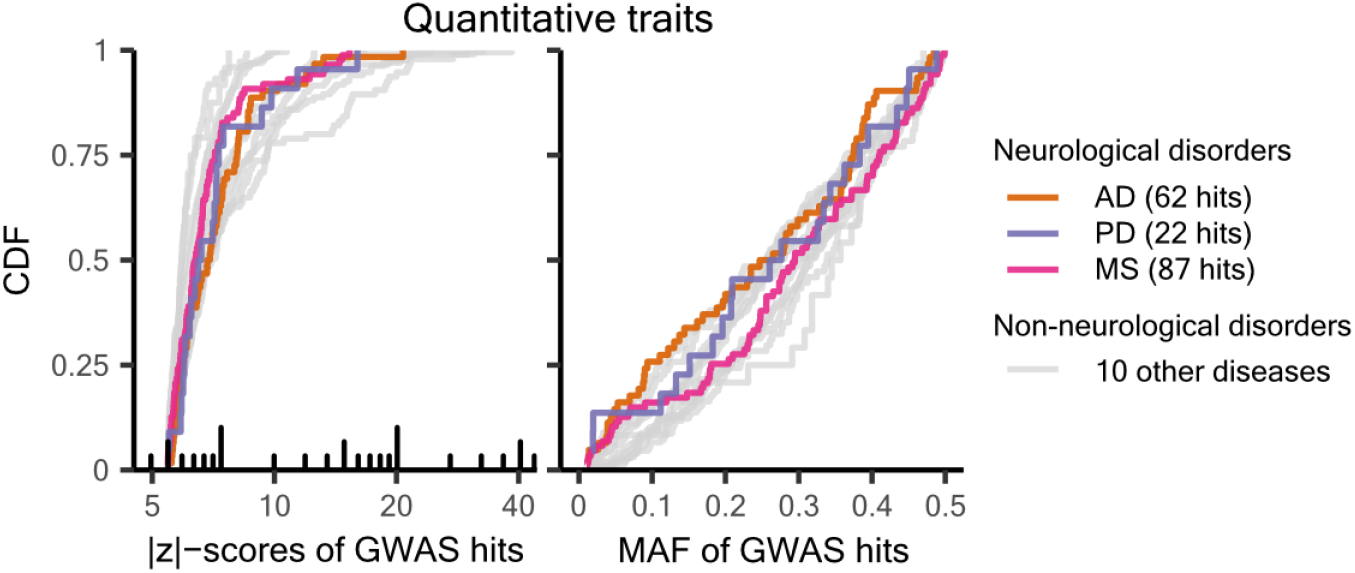
Contrasting the GWAS hits for neurological diseases with those for 10 other human complex diseases. *The full list of diseases can be found in Supplementary Table 2. AD: Alzheimer’s disease; PD: Parkinson’s disease; MS: multiple sclerosis*.

**Figure S8:**
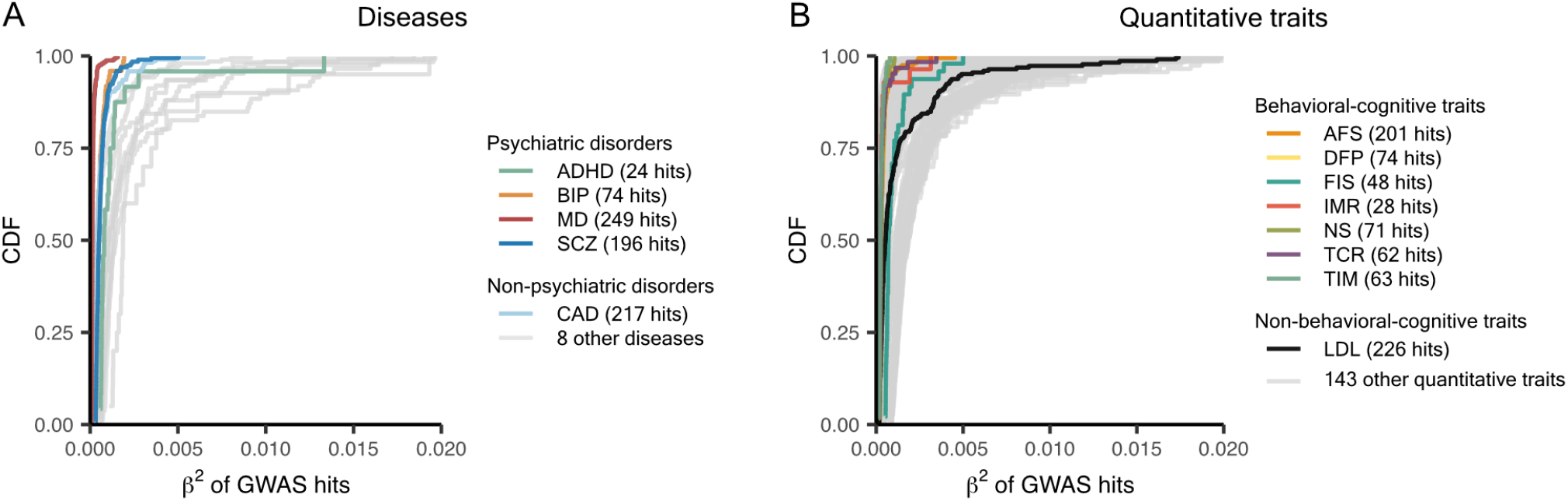
Squared effect sizes of GWAS hits for brain-related versus non–brain-related traits. ***A)*** *Psychiatric disorders compared with nine other complex diseases. Effect sizes reported on the log odds ratio scale (ζ) are transformed to the liability scale (β) with Equation 6. Details of the derivation are provided in the Supplementary Notes. When applying Equation 6, we assumed disease prevalence as reported in Supplementary Table 3*. ***B)*** *Behavioral-cognitive traits compared with 144 other quantitative traits*.

**Figure S9:**
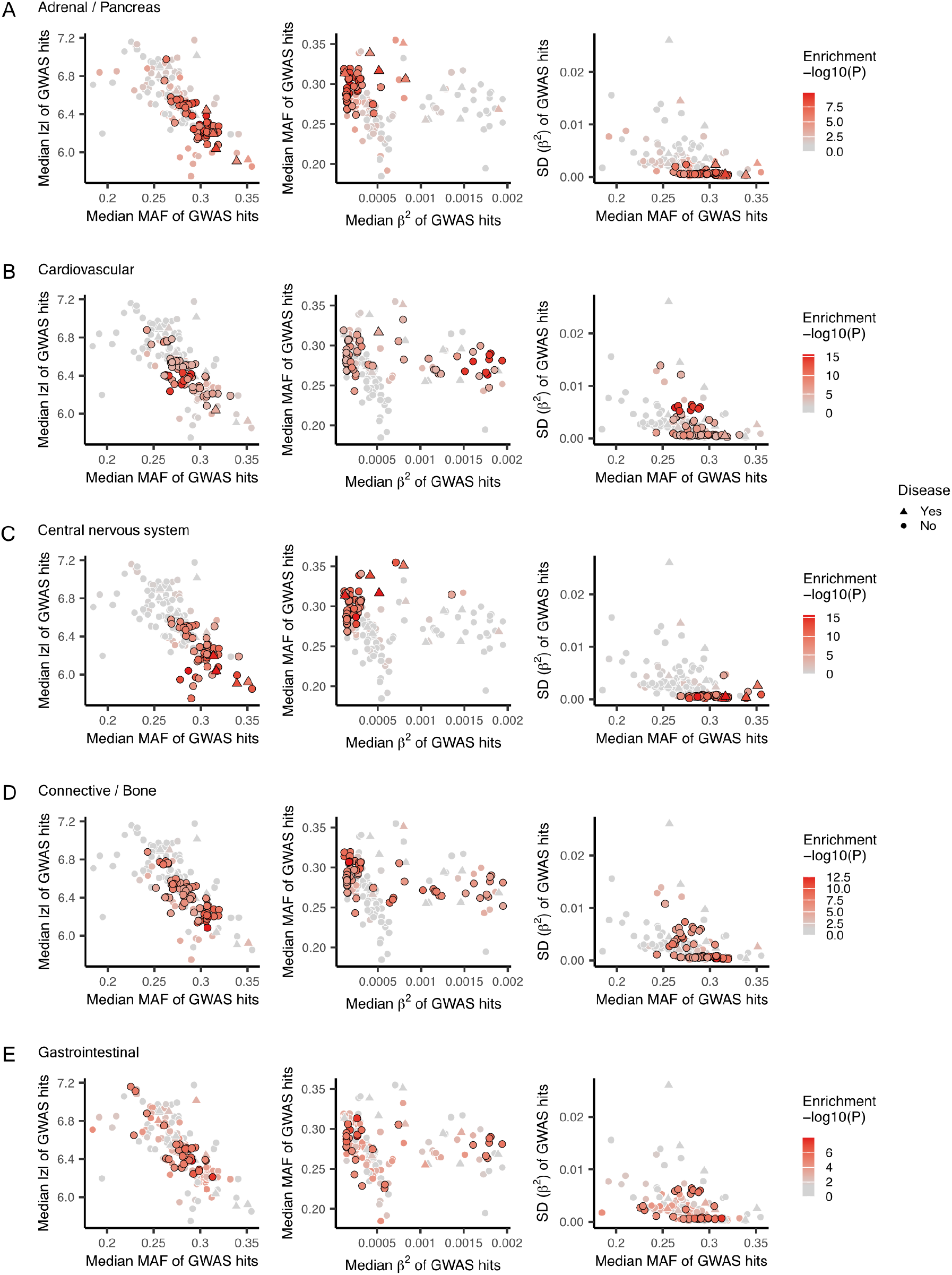
Parameterizing the genetic architectures of complex traits (first half). *Pairwise combinations of four GWAS hit metrics — median MAF, median* |*z*|*-score, median β*^2^, *and standard deviation of β*^2 —^ *were used to parameterize trait architectures. All 164 traits included in this study are colored by meta-analyzed S-LDSC p-values for functional enrichment in the* ***A)*** *Adrenal / Pancreas* ***B)*** *Cardiovascular* ***C)*** *Central nervous system* ***D)*** *Connective / Bone* ***E)*** *Gastrointestinal category. Traits with p-values exceeding the significance threshold of* 0.05/(10 *×* 164) *are outlined with black borders*.

**Figure S10:**
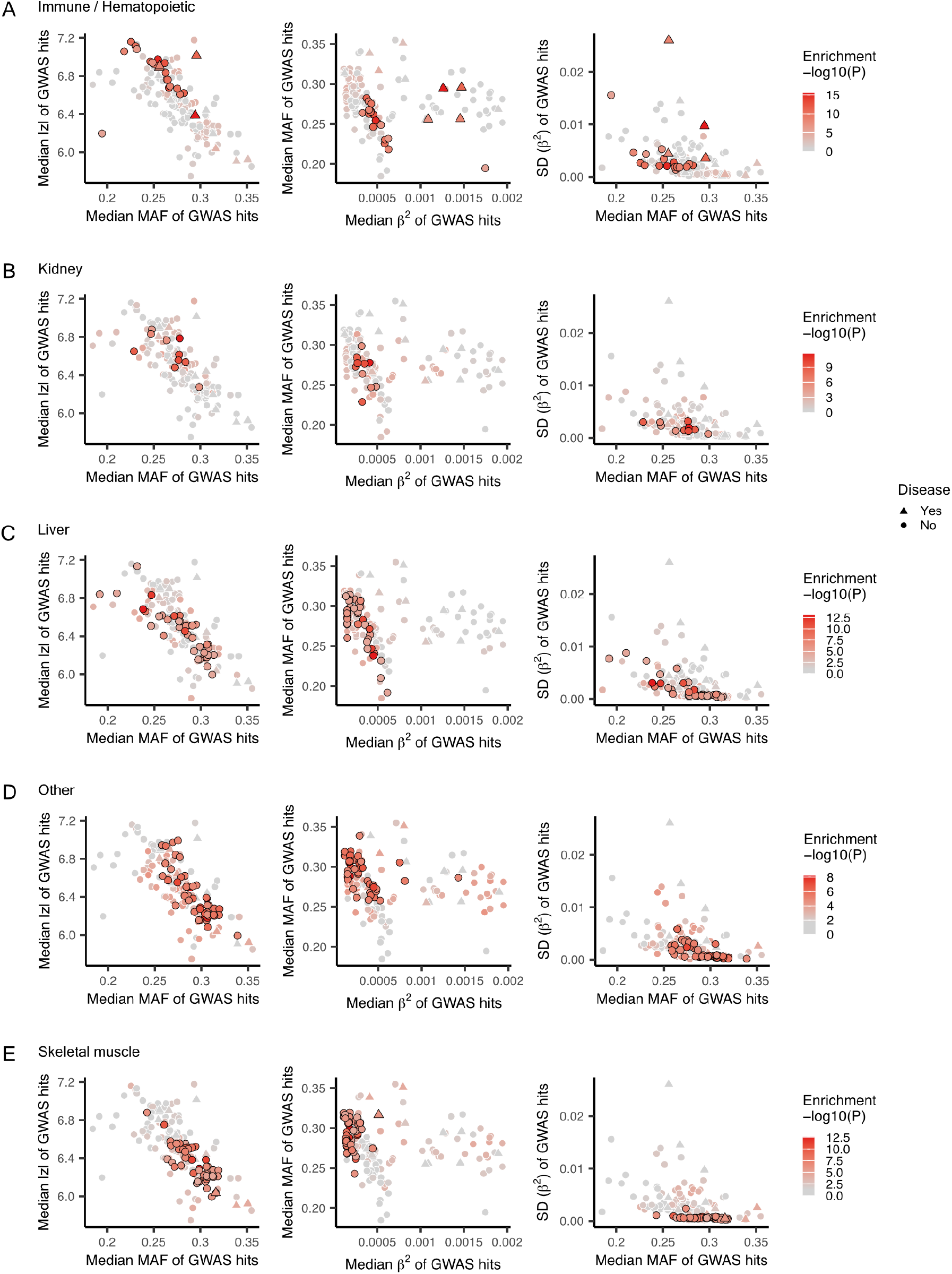
Parameterizing the genetic architectures of complex traits (second half). *Pairwise combinations of four GWAS hit metrics — median MAF, median* |*z*|*-score, median β*^2^, *and standard deviation of β*^2 —^*were used to parameterize trait architectures. All 164 traits included in this study are colored by meta-analyzed S-LDSC p-values for functional enrichment in the* ***A)*** *Immune / Hematopoietic* ***B)*** *Kidney* ***C)*** *Liver* ***D)*** *Other* ***E)*** *Skeletal muscle category. Traits with p-values exceeding the significance threshold of* 0.05/(10 *×* 164) *are outlined with black borders*.

**Figure S11:**
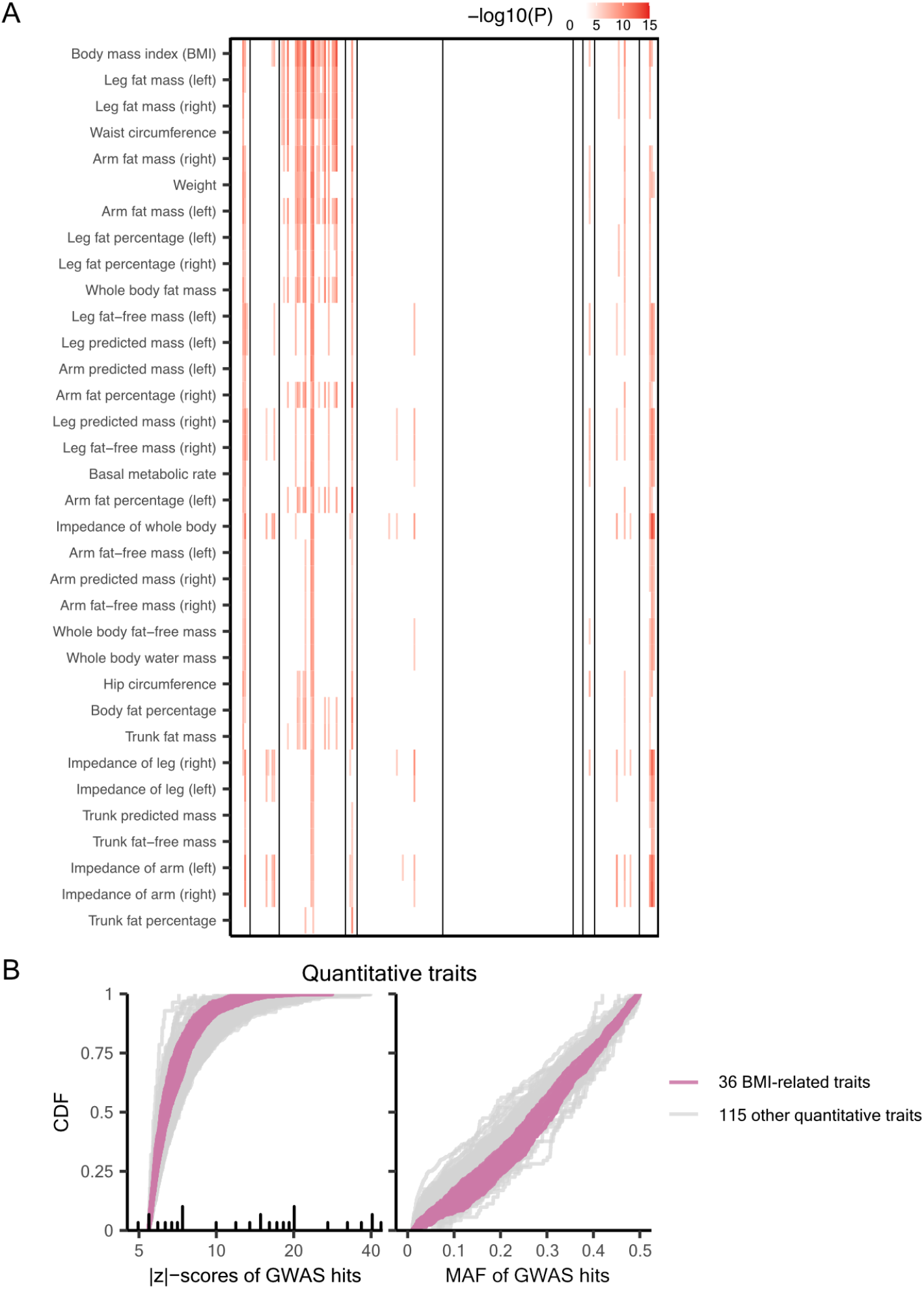
Contrasting body composition-related traits with other quantitative traits. ***A)*** *S-LDSC profiles of the 36 body composition-related traits extracted from Figure S4*. ***B)*** *CDF curves for GWAS hits. The UK Biobank data fields for the 36 body composition–related traits are 48, 49, 21001, 21002, 23098–23102, and 23104–23130. Gray lines correspond to the rest of the 115 quantitative traits listed in Supplementary Table 1*.

**Figure S12:**
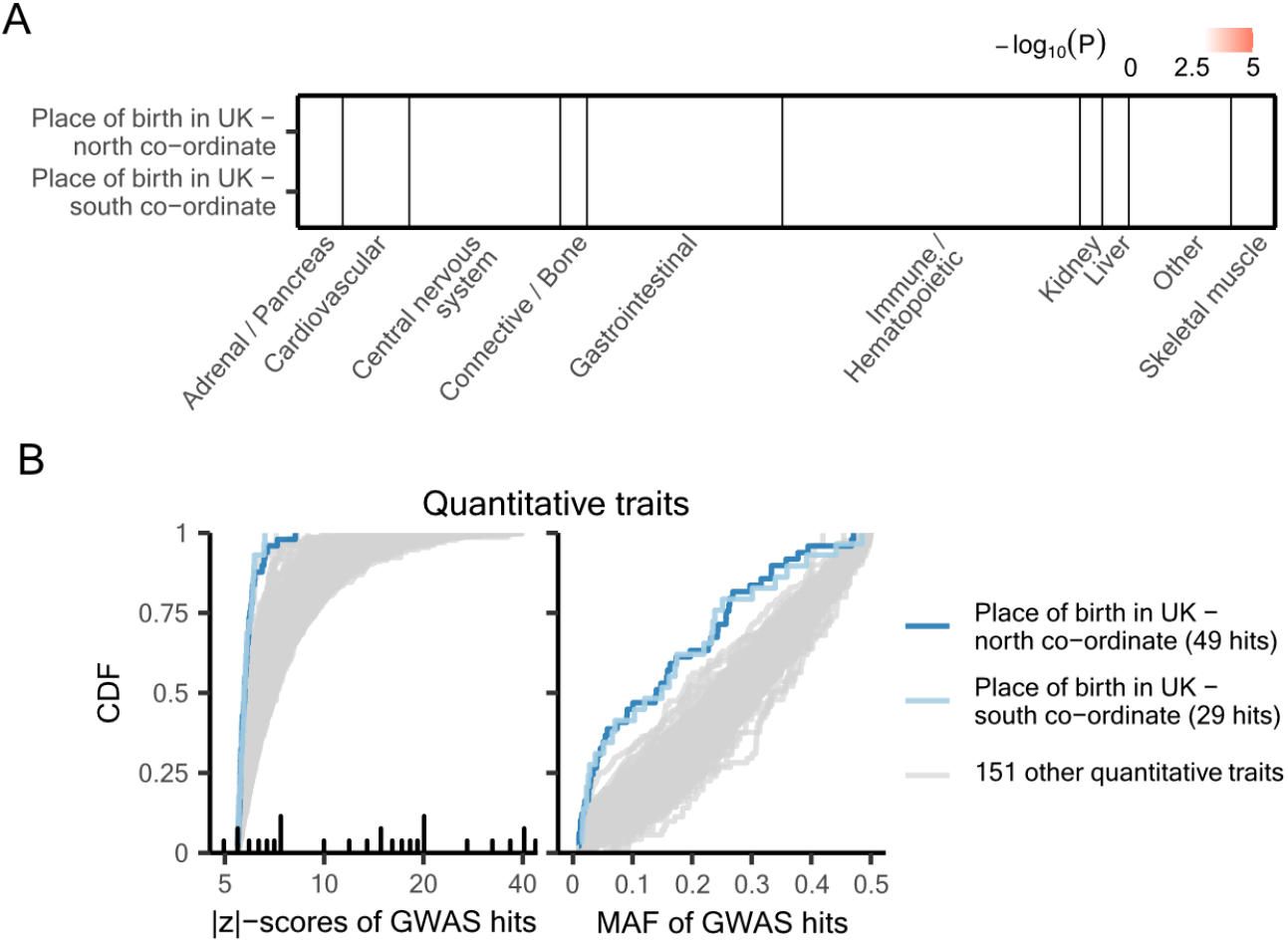
Contrasting two birth coordinate traits with 151 selected quantitative traits. ***A)*** *S-LDSC enrichment results*. ***B)*** *CDF curves for GWAS hits. UK Biobank data fields are 129 for Place of birth in UK - north co-ordinate and 130 for Place of birth in UK - south co-ordinate. Same as the other quantitative traits, GWAS summary statistics for these two birth coordinate traits were downloaded from the Neale Lab [35], with 20 genotyping PCs included as covariates. The full list of quantitative traits can be found in Supplementary Table 1*.

**Figure S13:**
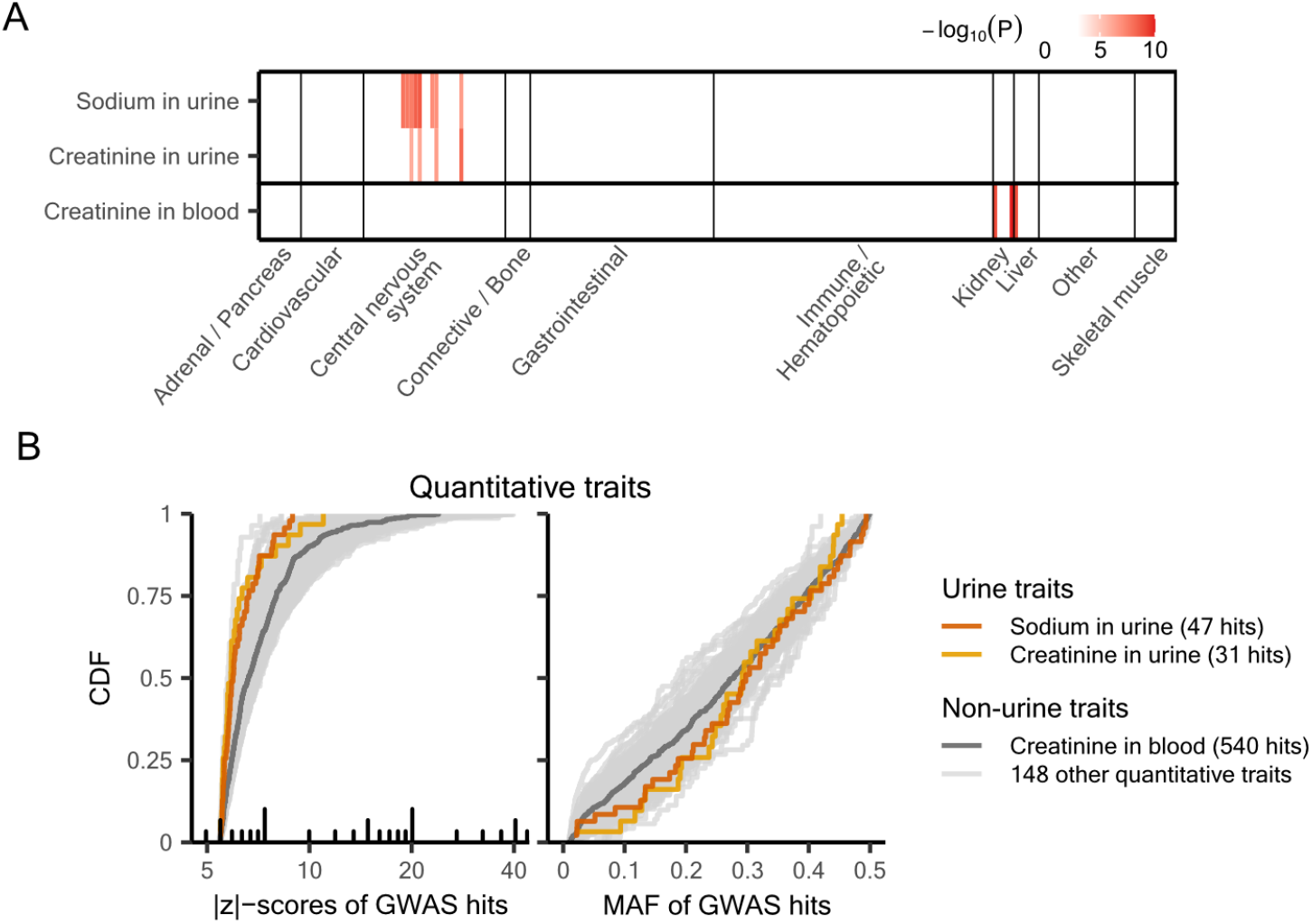
Contrasting urinary traits with serum creatinine (and 148 other quantitative traits). ***A)*** *S-LDSC enrichment results*. ***B)*** *CDF curves for GWAS hits. The full list of quantitative traits can be found in Supplementary Table 1. UK Biobank data fields are 30530 for sodium in urine, 30510 for creatinine in urine, and 30700 for creatinine in blood*.

**Figure S14:**
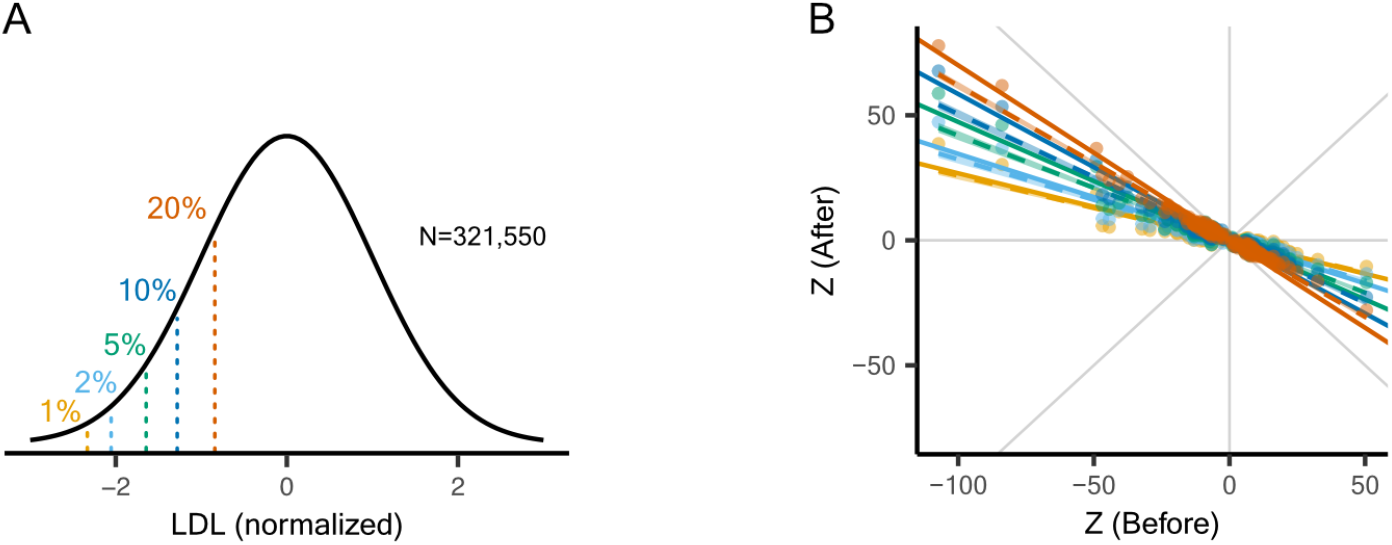
LDL GWAS power reduction after binarization. ***A)*** *Binarizing LDL levels with varying prevalence in the lower tail*. ***B)*** *Deflation of GWAS hit Z scores following lower-tail binarization of LDL. Different colors correspond to different prevalence used in panel A. Solid lines represent predicted slopes, and dashed lines indicate fitted slopes. Gray lines in the background are y* = 0, *x* = 0 *and y* = *±x. LDL was measured in unrelated White British individuals from the UK Biobank (data field 30780). The original phenotypic distribution was standardized using rank-based inverse normal transformation*.

**Figure S15:**
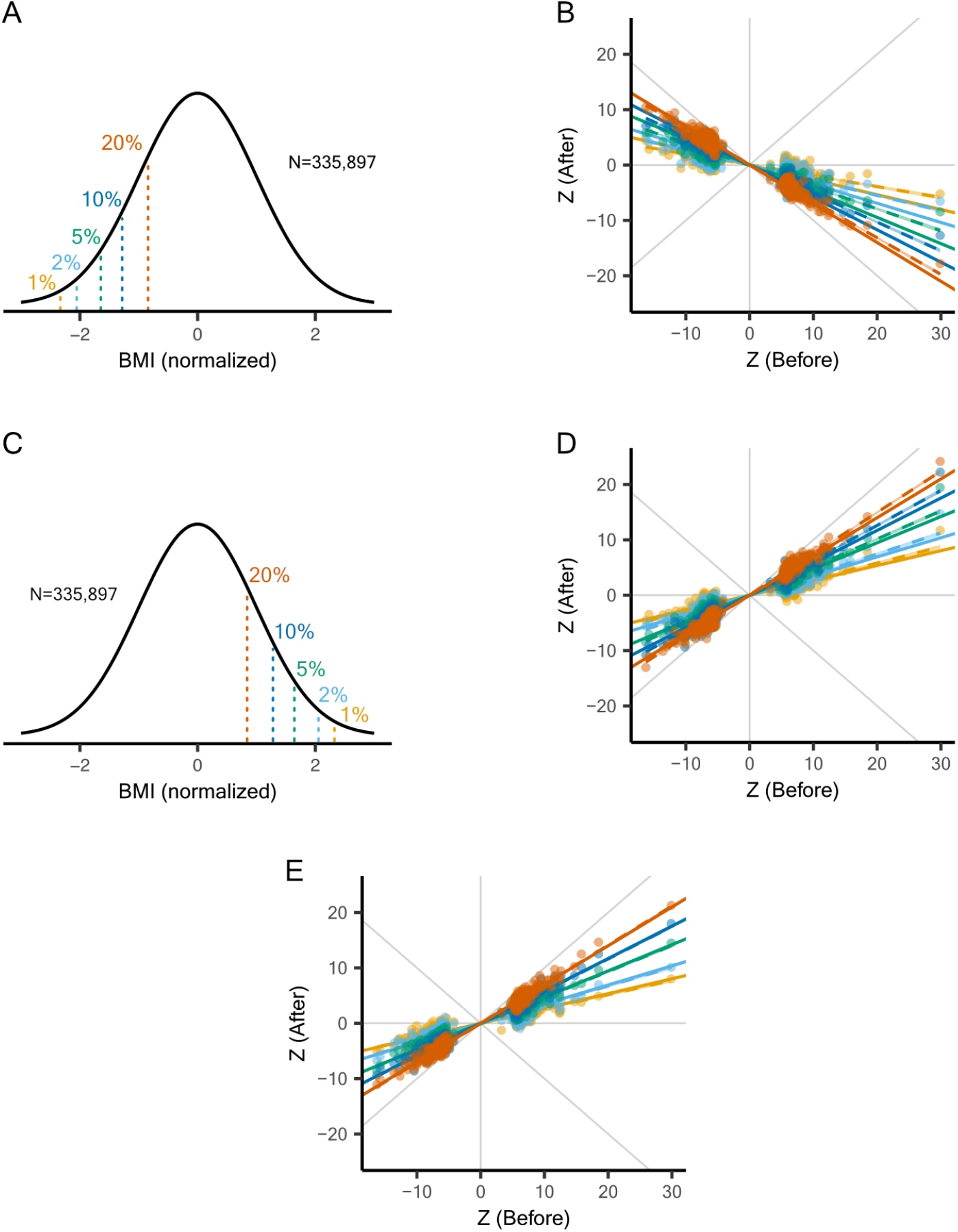
BMI GWAS power reduction after binarization or downsampling. *Binarizing BMI with varying prevalence in the* ***A)*** *lower tail or the* ***C)*** *upper tail. Deflation of GWAS hit Z scores after* ***B)*** *binarization in the lower tail*, ***D)*** *binarization in the upper tail, or* ***E)*** *downsampling BMI to matched sample sizes. In panels B, D and E, different colors correspond to different prevalence used in panels A and C. Solid lines represent predicted slopes, and dashed lines indicate fitted slopes. Gray lines in the background are y* = 0, *x* = 0 *and y* = *±x. BMI was measured in unrelated White British individuals from the UK Biobank (data field 21001). The original phenotypic distribution was standardized using rank-based inverse normal transformation*.

**Figure S16:**
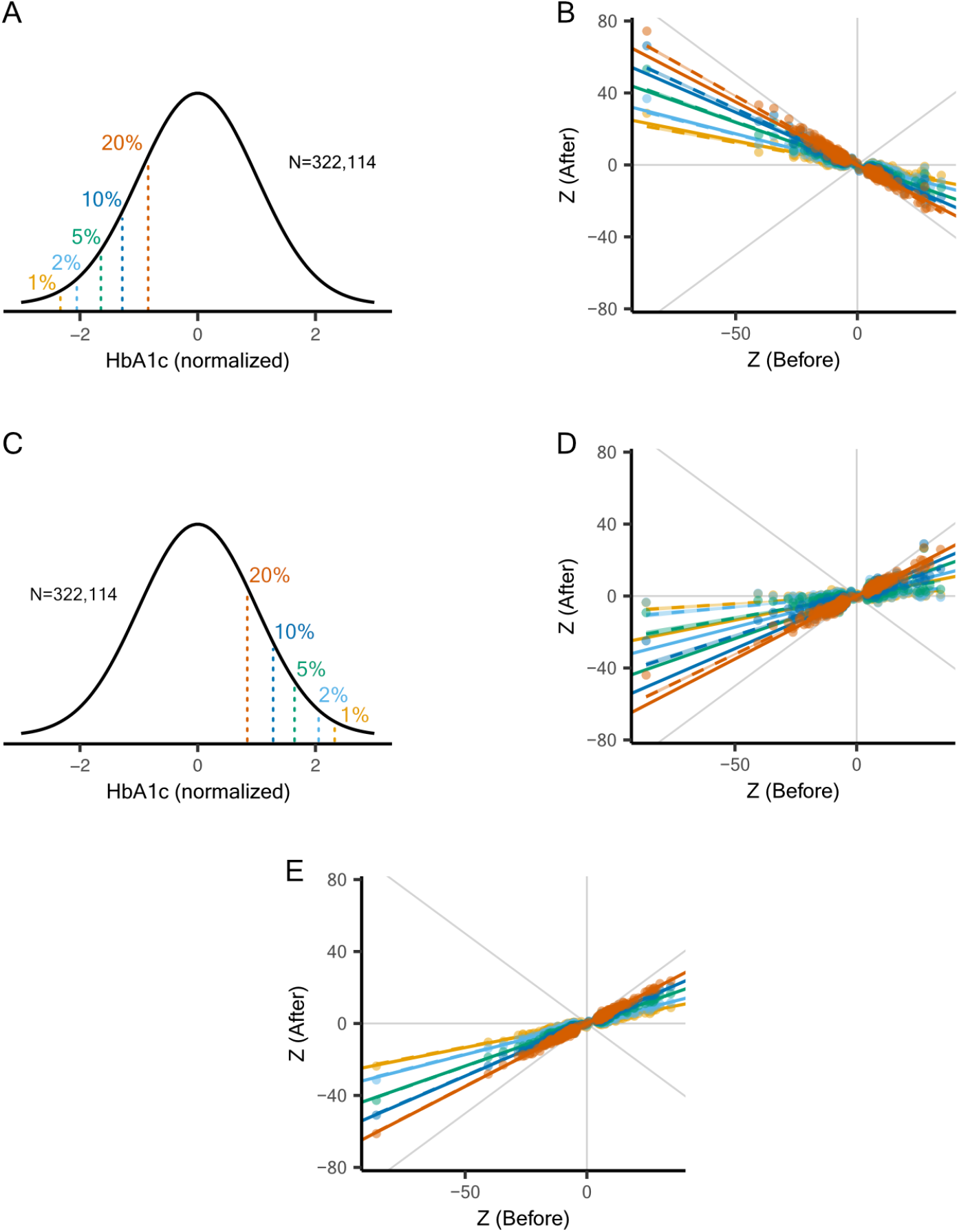
HbA1c GWAS power reduction after binarization or downsampling. *Binarizing hemoglobin A1C (HbA1c) with varying prevalence in the* ***A)*** *lower tail or the* ***C)*** *upper tail. Deflation of GWAS hit Z scores after* ***B)*** *binarization in the lower tail*, ***D)*** *binarization in the upper tail, or* ***E)*** *downsampling HbA1c to matched sample sizes. In panels B, D and E, different colors correspond to different prevalence used in panels A and C. Solid lines represent predicted slopes, and dashed lines indicate fitted slopes. Gray lines in the background are y* = 0, *x* = 0 *and y* = *±x. HbA1c was measured in unrelated White British individuals from the UK Biobank (data field 30750). The original phenotypic distribution was standardized using rank-based inverse normal transformation*.

**Figure S17:**
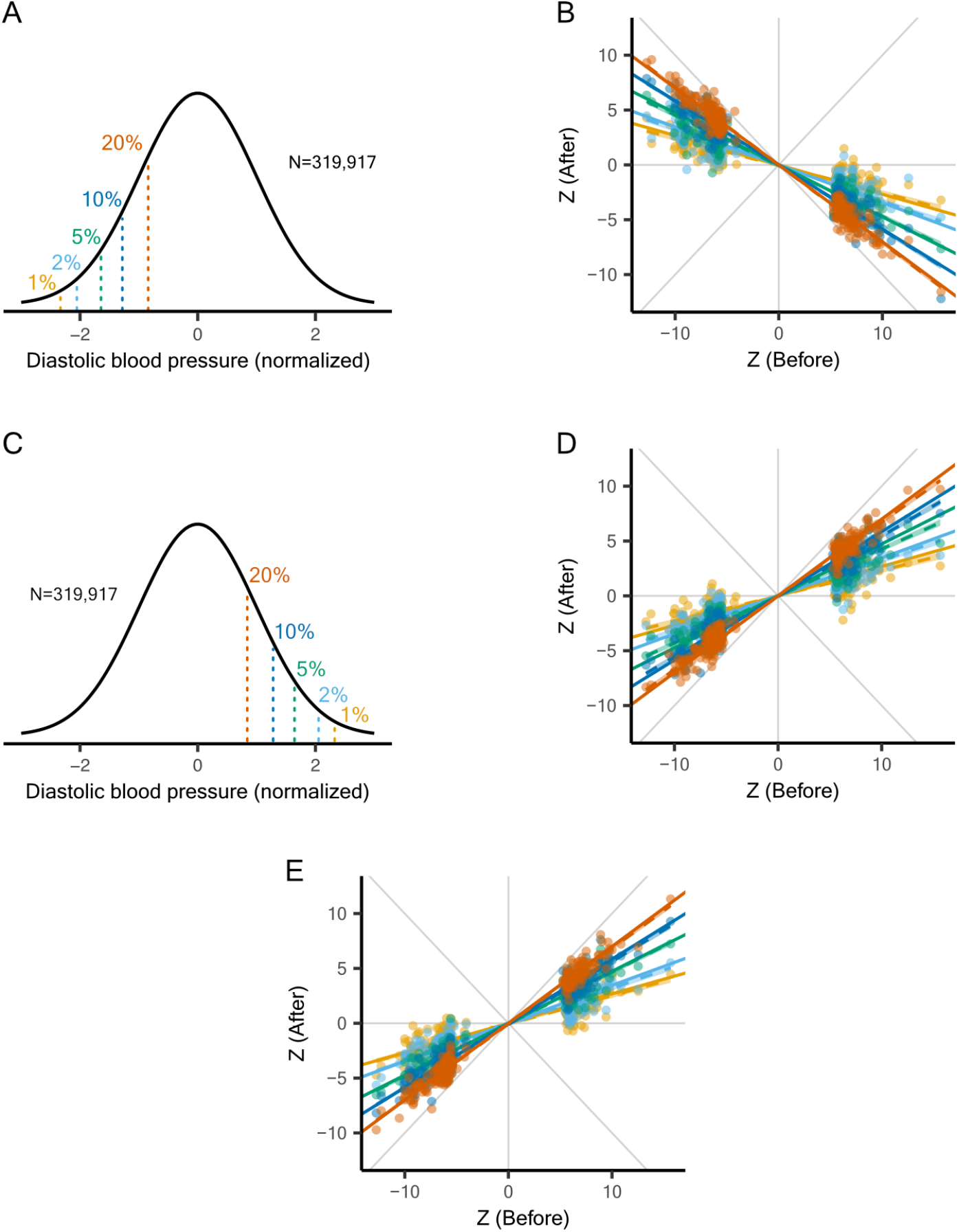
Diastolic blood pressure GWAS power reduction after binarization or downsampling. *Binarizing diastolic blood pressure with varying prevalence in the* ***A)*** *lower tail or the* ***C)*** *upper tail. Deflation of GWAS hit Z scores after* ***B)*** *binarization in the lower tail*, ***D)*** *binarization in the upper tail, or* ***E)*** *downsampling diastolic blood pressure to matched sample sizes. In panels B, D and E, different colors correspond to different prevalence used in panels A and C. Solid lines represent predicted slopes, and dashed lines indicate fitted slopes. Gray lines in the background are y* = 0, *x* = 0 *and y* = *±x. Diastolic blood pressure was measured in unrelated White British individuals from the UK Biobank (data field 4079). The original phenotypic distribution was standardized using rank-based inverse normal transformation*.

**Figure S18:**
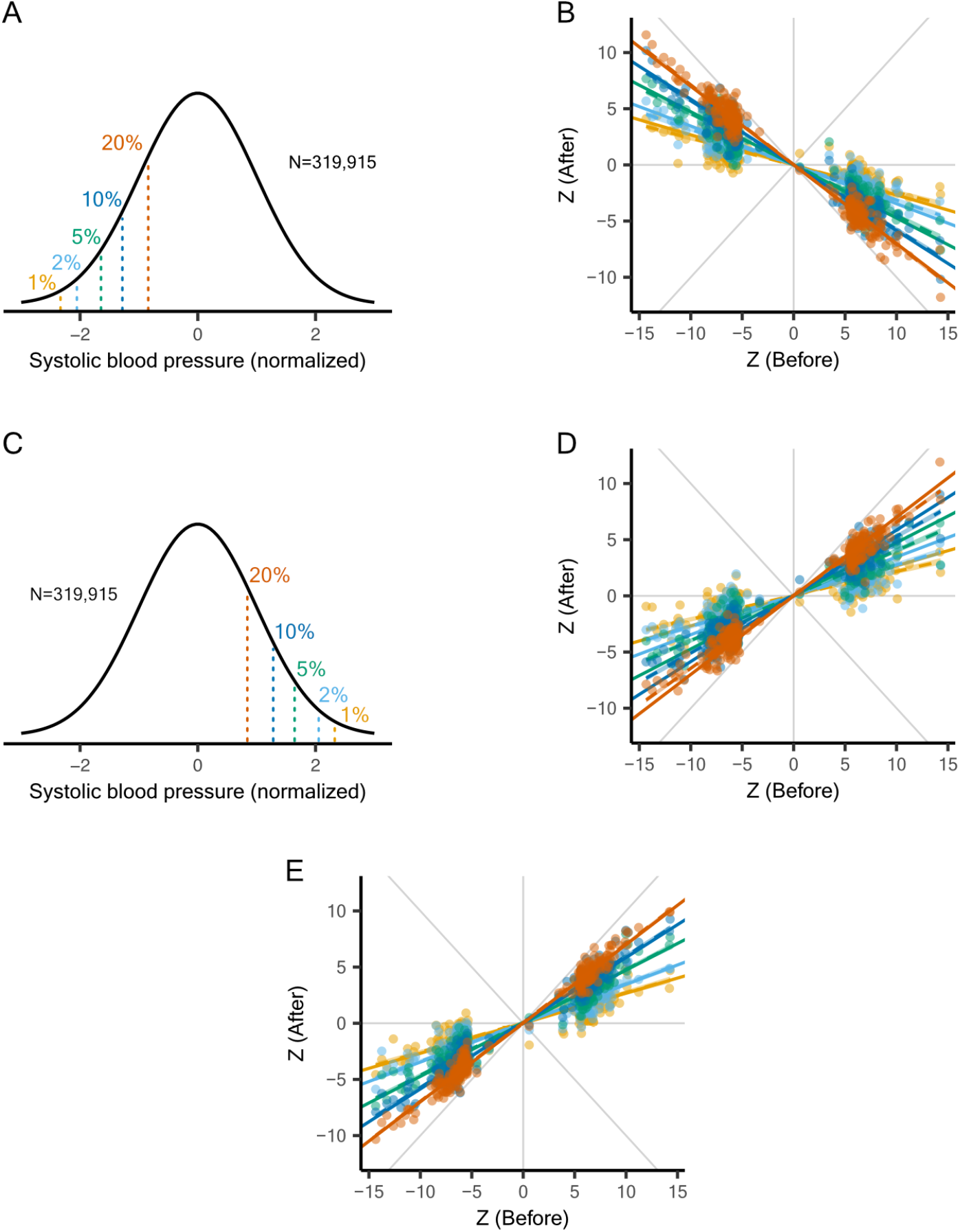
Systolic blood pressure GWAS power reduction after binarization or downsampling. *Binarizing systolic blood pressure with varying prevalence in the* ***A)*** *lower tail or the* ***C)*** *upper tail. Deflation of GWAS hit Z scores after* ***B)*** *binarization in the lower tail*, ***D)*** *binarization in the upper tail, or* ***E)*** *downsampling systolic blood pressure to matched sample sizes. In panels B, D and E, different colors correspond to different prevalence used in panels A and C. Solid lines represent predicted slopes, and dashed lines indicate fitted slopes. Gray lines in the background are y* = 0, *x* = 0 *and y* = *±x. Systolic blood pressure was measured in unrelated White British individuals from the UK Biobank (data field 4080). The original phenotypic distribution was standardized using rank-based inverse normal transformation*.

**Figure S19:**
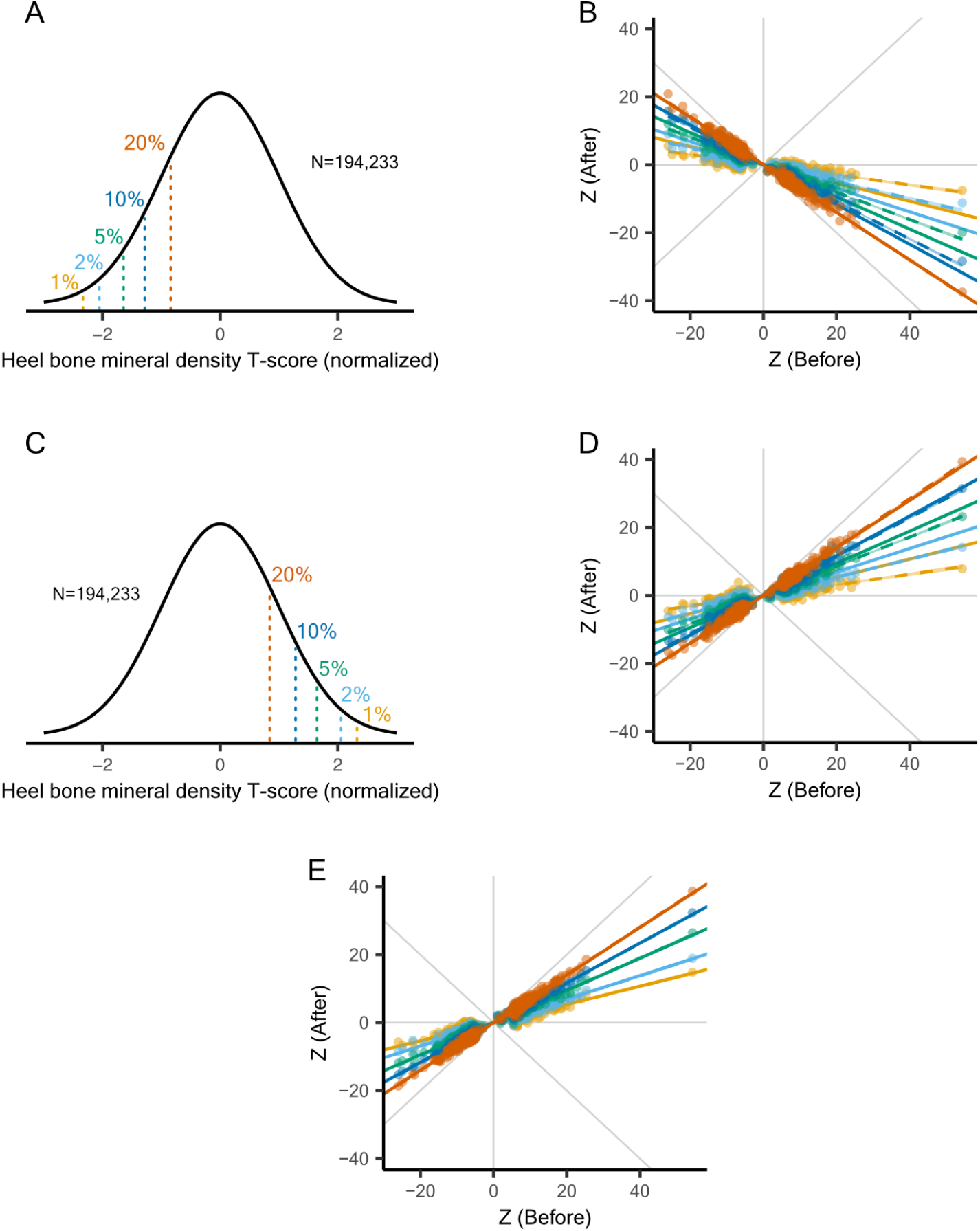
Heel bone mineral density T-score GWAS power reduction after binarization or downsampling. *Binarizing heel bone mineral density T-score with varying prevalence in the* ***A)*** *lower tail or the* ***C)*** *upper tail. Deflation of GWAS hit Z scores after* ***B)*** *binarization in the lower tail*, ***D)*** *binarization in the upper tail, or* ***E)*** *downsampling heel bone mineral density T-score to matched sample sizes. In panels B, D and E, different colors correspond to different prevalence used in panels A and C. Solid lines represent predicted slopes, and dashed lines indicate fitted slopes. Gray lines in the background are y* = 0, *x* = 0 *and y* = *±x. Heel bone mineral density T-score was measured in unrelated White British individuals from the UK Biobank (data field 78). The original phenotypic distribution was standardized using rank-based inverse normal transformation*.

**Figure S20:**
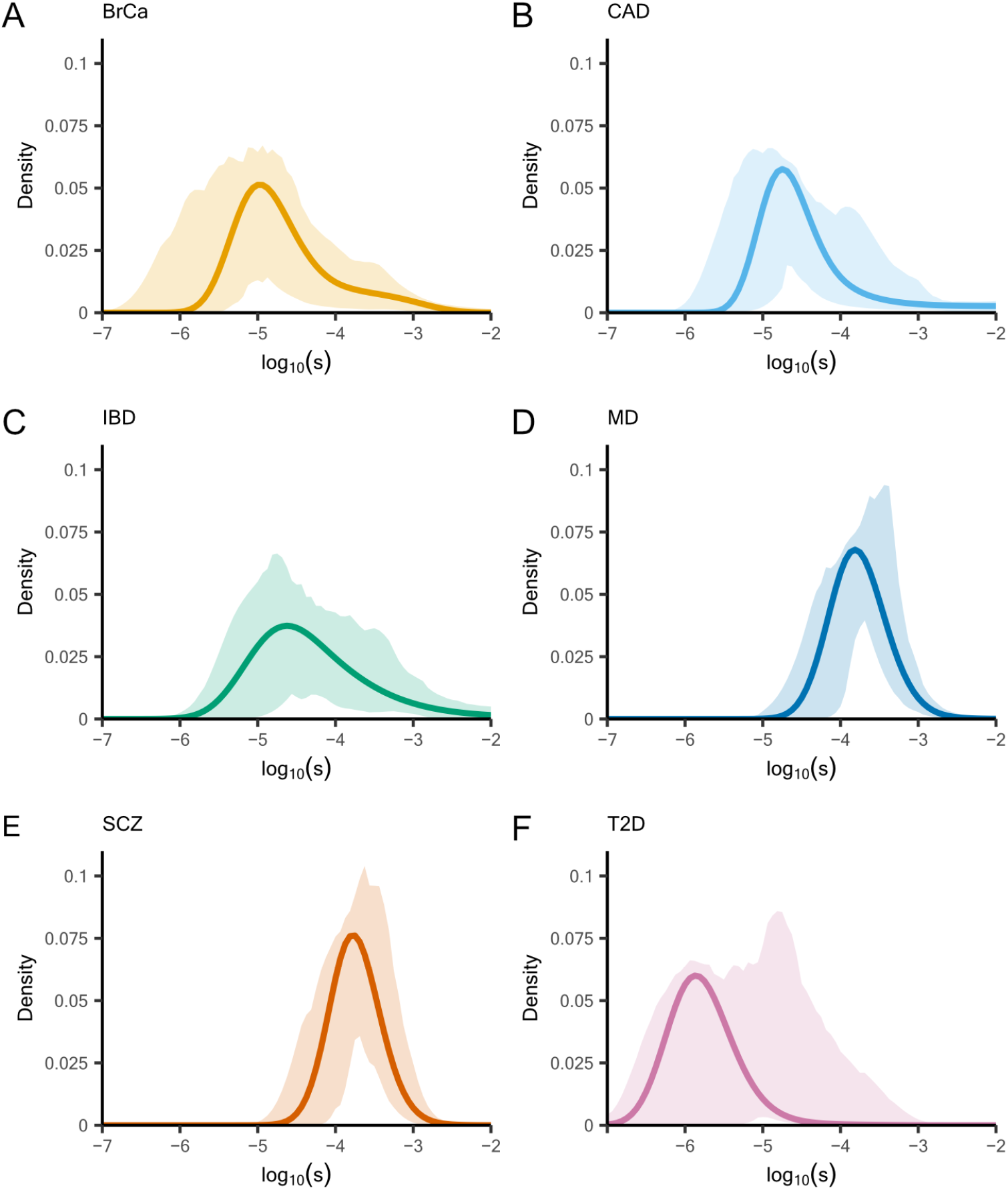
Trait-specific distributions of selection coefficients inferred from. ***A)*** *breast cancer* ***B)*** *coronary artery disease* ***C)*** *inflammatory bowel disease* ***D)*** *major depression* ***E)*** *schizophrenia* ***F)*** *type 2 diabetes GWAS hits, with 90% confidence envelopes shown*.

**Figure S21:**
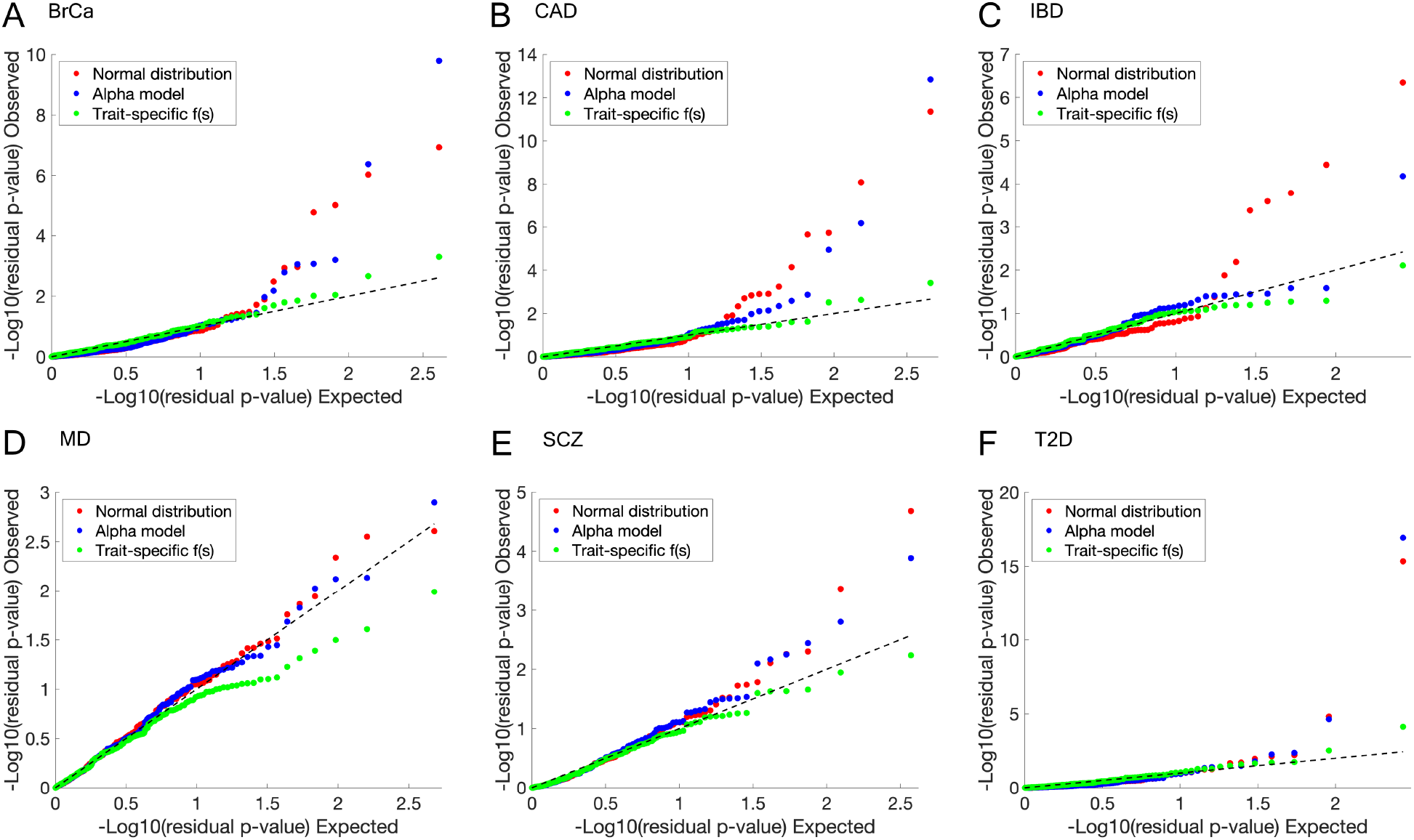
QQ plots of residual p-values for. ***A)*** *breast cancer* ***B)*** *coronary artery disease* ***C)*** *inflammatory bowel disease* ***D)*** *major depression* ***E)*** *schizophrenia* ***F)*** *type 2 diabetes*, **evaluated under three models: the normal distribution, the alpha model, and the adapted Simons model, which infers trait-specific** *f* (*s*).

**Figure S22:**
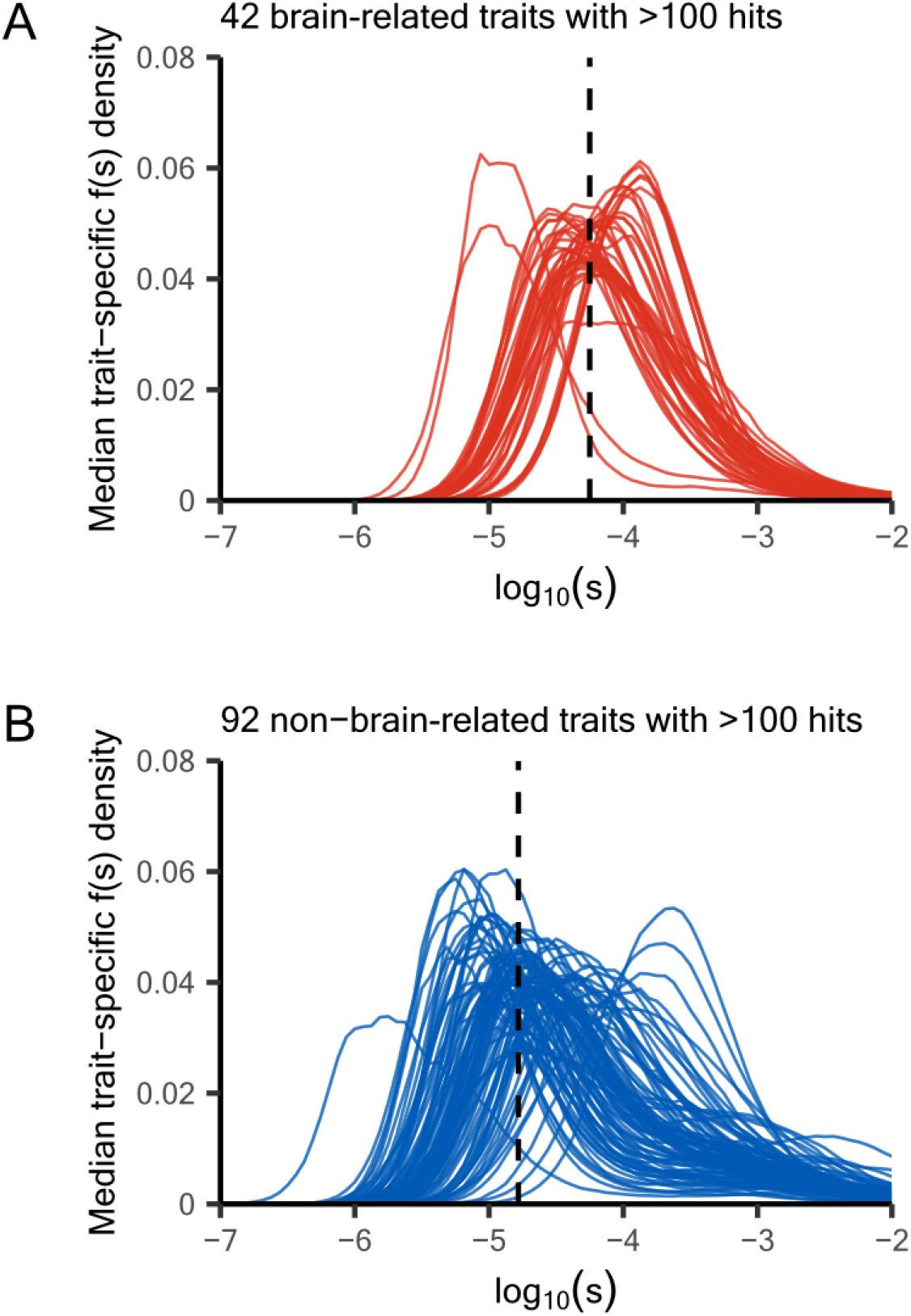
Median trait-specific distributions of selection coefficients inferred from 134 traits with at least 100 GWAS hits, with traits grouped into. ***A)*** *brain-related* **versus *B)*** *non-brain-related* **traits**. *Vertical line in each plot correspond to the median mode*.

**Figure S23:**
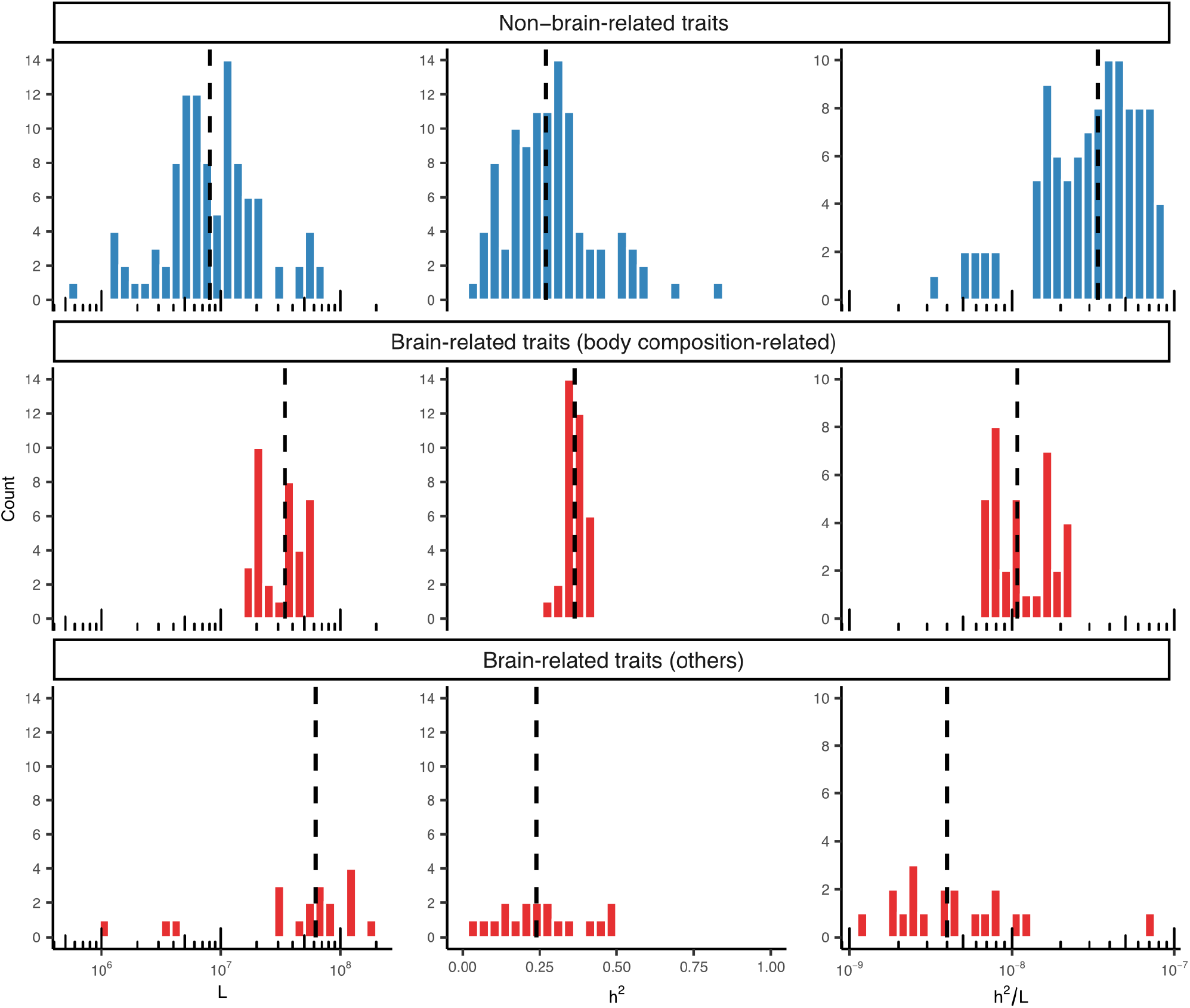
Re-plotting the estimates in Figure 4D as histograms to highlight distribution shifts. *Dashed vertical lines indicate the median values in each subplot. Brain-related traits are further divided into body composition-related traits and other brain-related traits. Body composition-related traits have L and h*^2^/*L estimates that fall between those of non–brain-related traits and traits primarily influenced by the CNS alone*.

**Figure S24:**
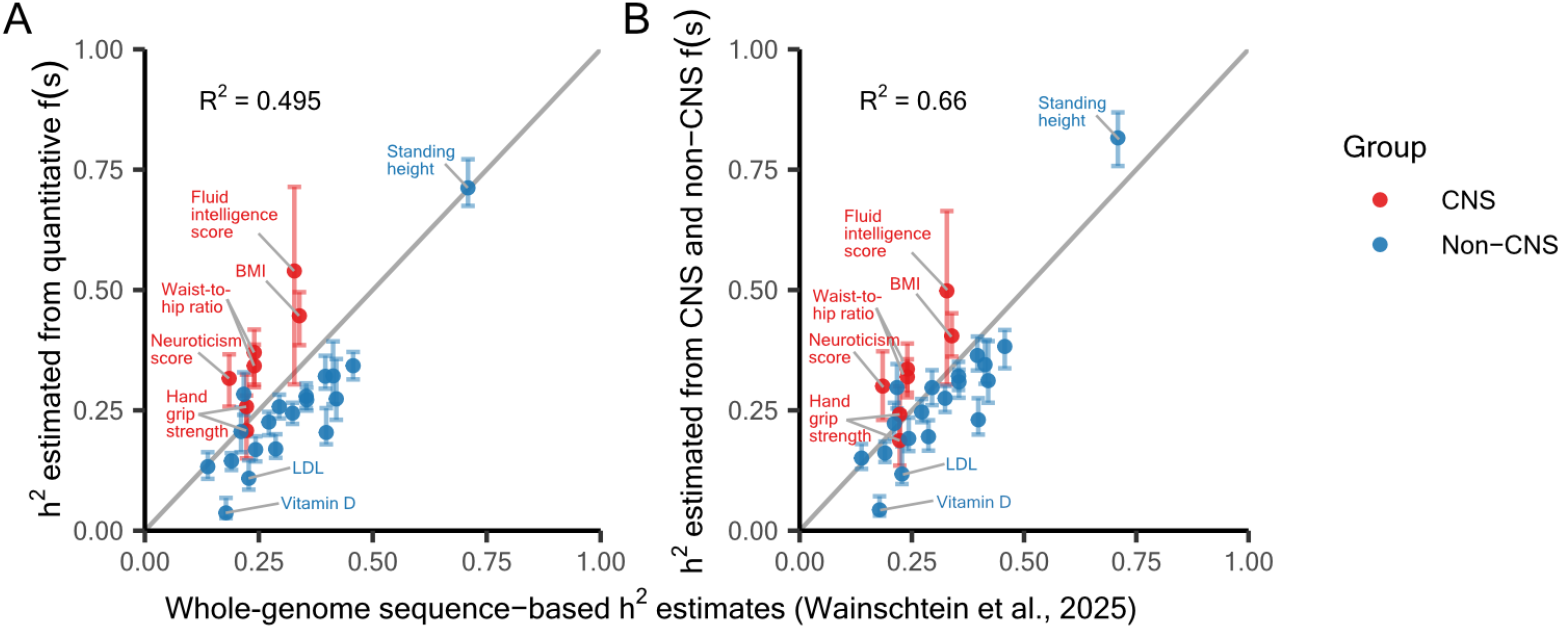
Comparing heritability estimates obtained in this study with recently reported whole-genome sequence-based heritability estimates 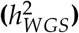. ***A)*** *Comparing* 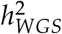 *with h*^2^ *obtained jointly with quantitative f* (*s*), *as shown in Figure 4B*. ***B)*** *Comparing* 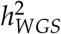 *with h*^2^ *obtained jointly with brain-related or non-brain-related f* (*s*), *as shown in Figure 4C. Wainschtein et al. [57] estimated* 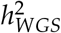 *in 347,630 unrelated individuals from the UK Biobank with GREML-LDMS* .

**Figure S25:**
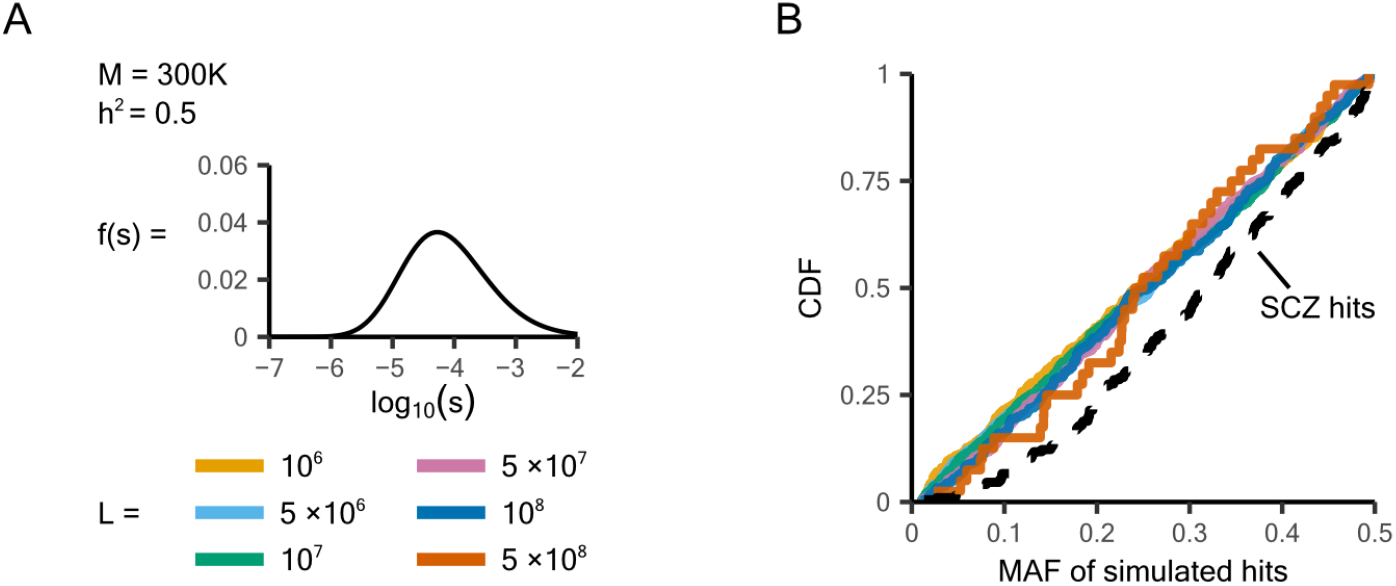
Variation in MAF distributions of GWAS hits cannot be recovered by changing *L*. ***A)*** *Same simulation setup as in Figure 5A*. ***B)*** *Varying L does not change the MAF distribution of GWAS hits. The black dashed line shows real schizophrenia GWAS hits for reference*.

**Figure S26:**
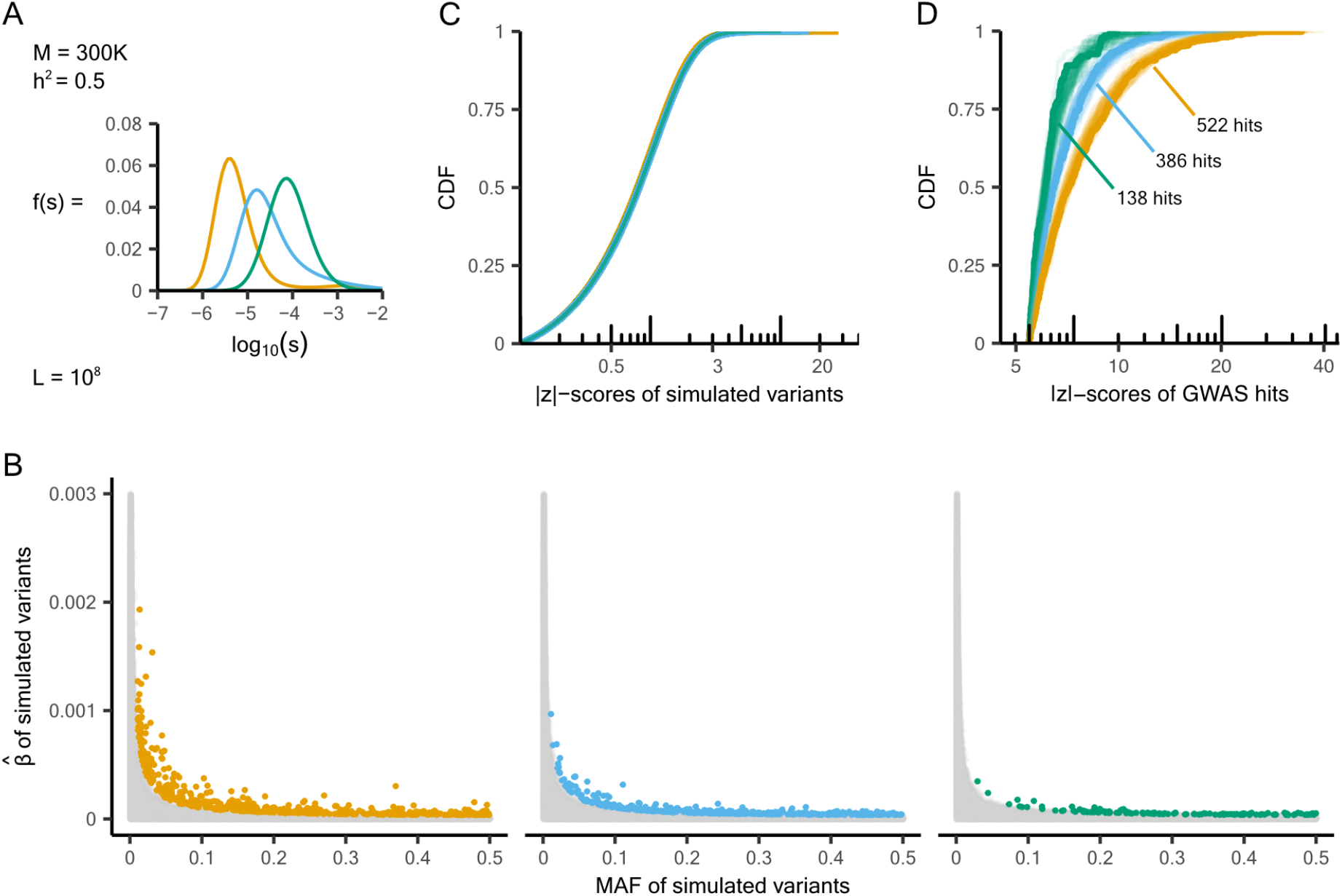
Impact of GWAS ascertainment on simulations under different *f* (*s*) distributions. ***A)*** *Same simulation setup as in Figure 5D*. ***B)*** *The GWAS ascertainment process. Variants with MAF* > 1% *and* |*Z*| > 5.45 *are discovered as hits and colored according to the f* (*s*) *distribution from which they were simulated. Unselected variants are shown in grey*. ***C)*** *Without GWAS ascertainment, the CDFs of* |*Z*| *scores for variants simulated under different f* (*s*) *distributions differ in the tails*. ***D)*** *After GWAS ascertainment, stronger f* (*s*) *produces steeper GWAS hit* |*Z*| *score CDFs and fewer hits*.

**Figure S27:**
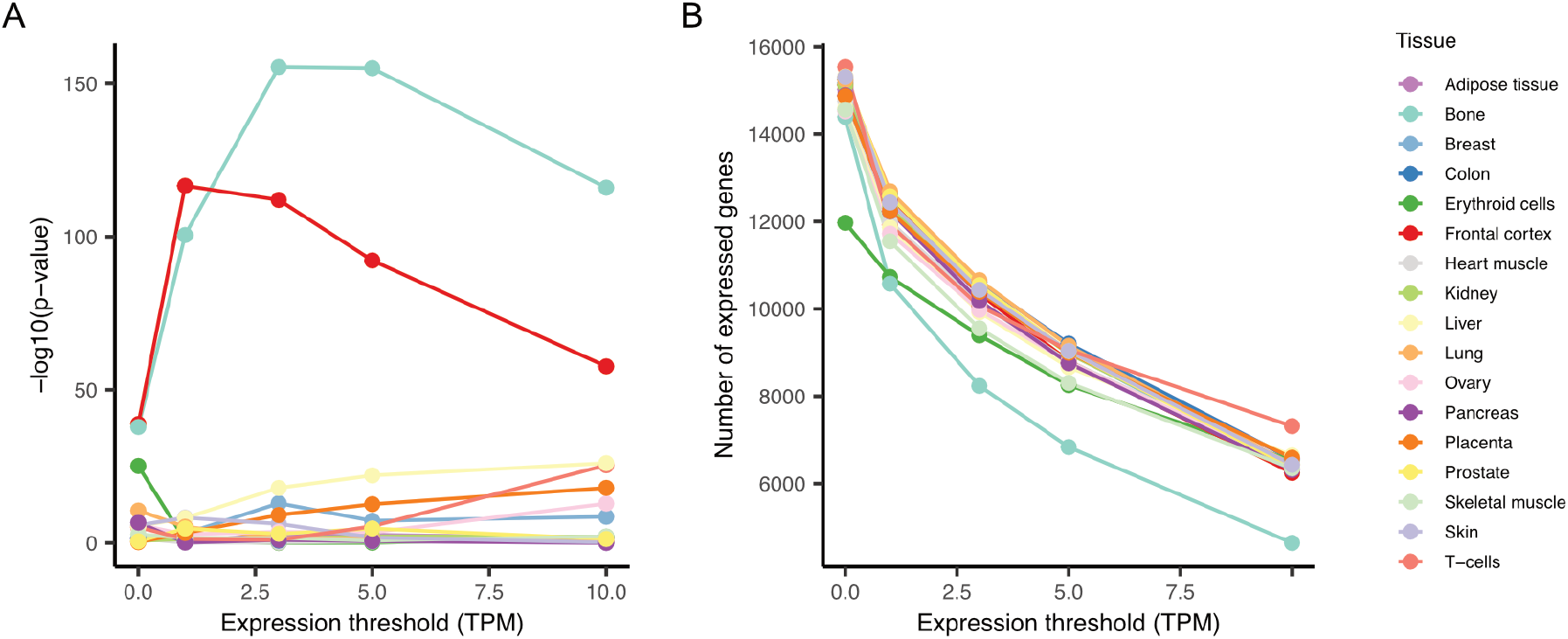
Brain expression status strongly predicts a gene’s constraint estimate, *s*_*het*_. ***A)*** *P-values from multiple regression models, where* log_10_(*s*_*het*_) *is regressed on expression status across 17 tissues. A gene is classified as “expressed” in a given tissue if its transcripts per million (TPM) values exceeds a threshold of 0, 1, 3, 5, or 10*. ***B)*** *While “frontal cortex” and “bone” appear to be the strongest predictors in panel A, genes are generally lowly expressed in the bone, and increasing the TPM cutoff rapidly excludes a large number of genes. In contrast, expression levels in the frontal cortex are more comparable to those in other tissues, indicating that its signal in panel A is not driven by a more rapid loss of genes at higher TPM thresholds. Gene expression values across tissues were curated in [39], with data extracted from the Human Protein Atlas [99]*.

**Figure S28:**
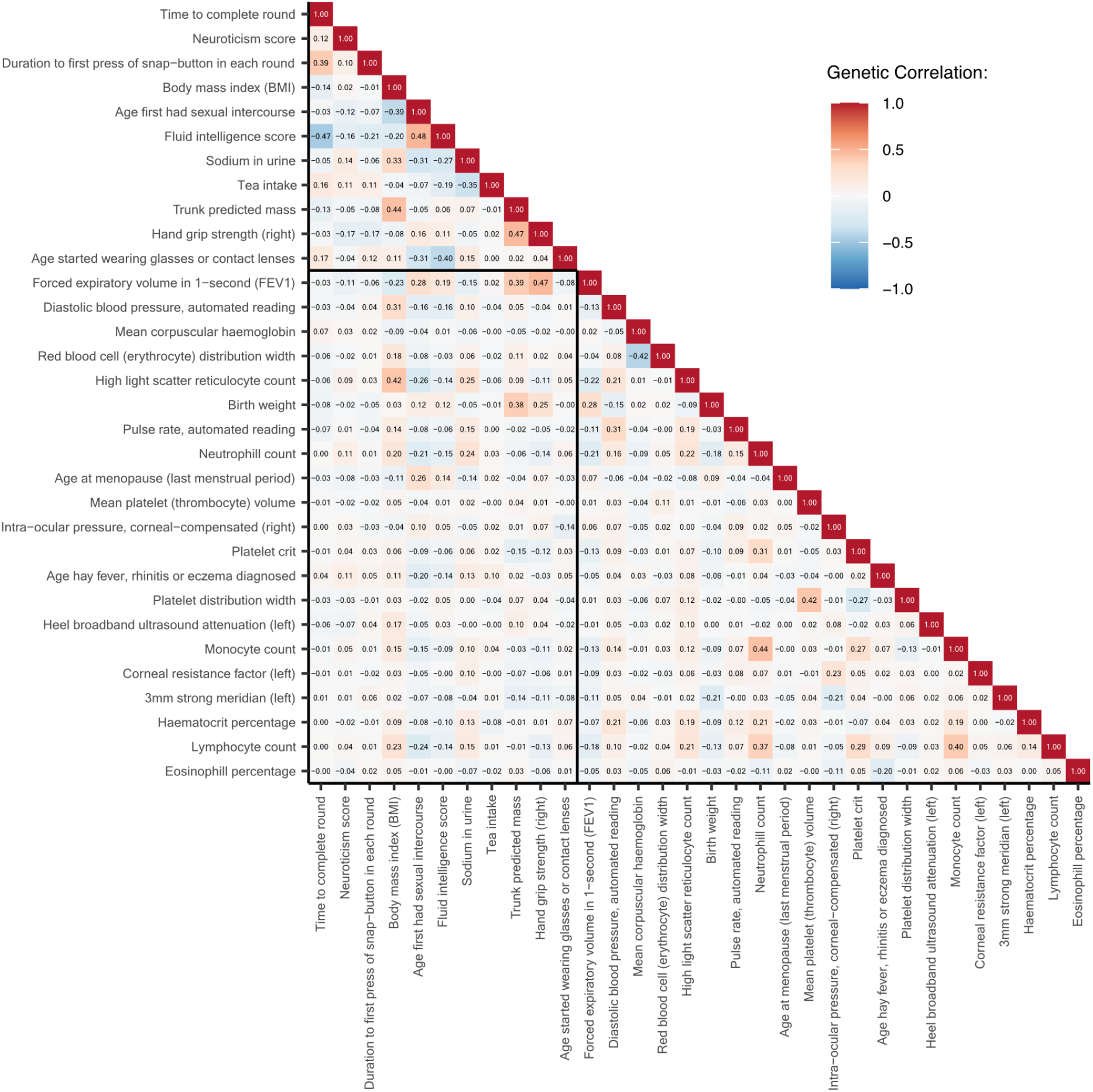
Heatmap of pairwise genetic correlations across 32 independent traits. *The top 11 traits are CNS traits*.

**Figure S29:**
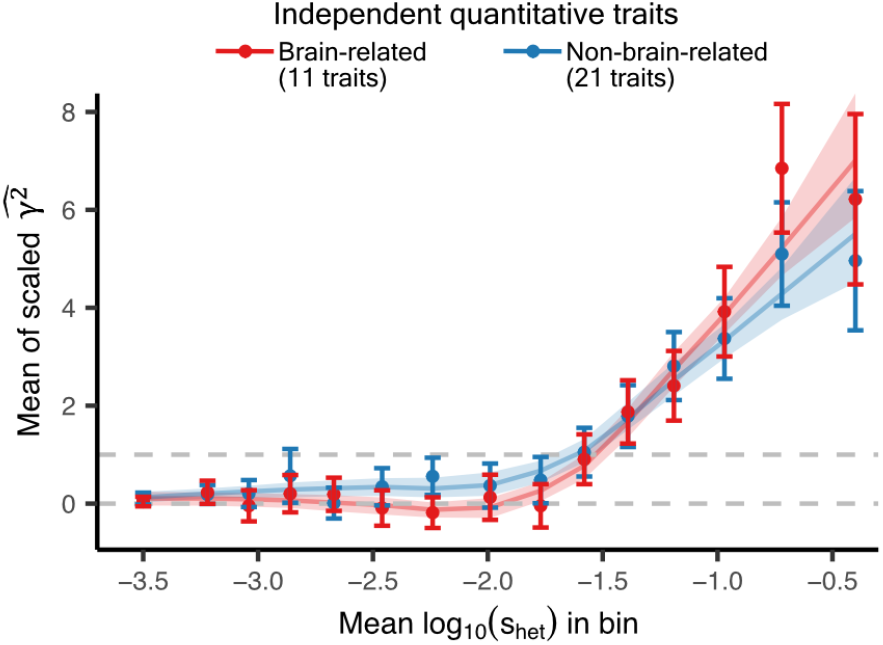
Mean squared genic effects on independent brain-related versus non-brain-related traits. *Genes are grouped into 15 quantiles of s*_*het*_ *estimates [58]. The estimated* 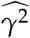 *values are scaled so that the mean across all gene-trait pairs in the tenth bin equals one. We applied a MAF* < 1% *filter. All estimates are shown with a* 95% *confidence interval. We bootstrapped genes at the trait level and showed the mean and 95% confidence intervals of the fitted LOESS curves*.

**Figure S30:**
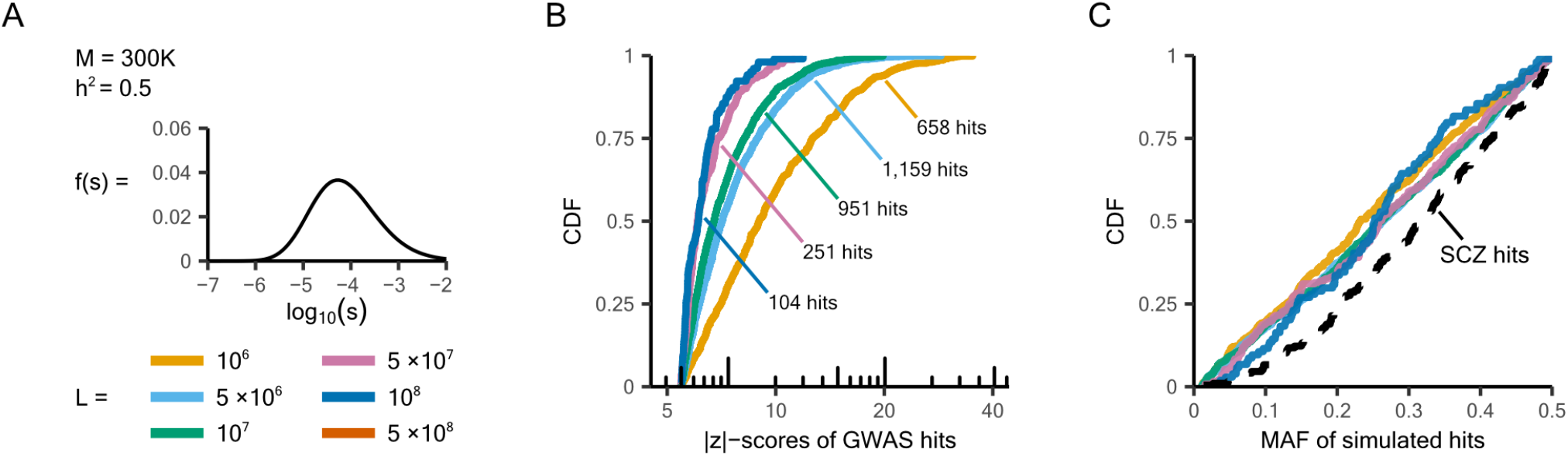
Assuming no pleiotropy for variant effects fails to recapitulate the observed variation in MAF distributions of GWAS hits. ***A)*** *Simulating under a modified Simons model that assumes no pleiotropy (Methods). The rest of the simulation set up is the same as in Figure 5A and S25*. ***B)*** *Larger L values lead to narrower distributions of GWAS hit* |*z*|*-scores. Simulating with a mutational target size of* 5 *×* 10^8^ *did not generate any GWAS hits*. ***C)*** *Varying L does not change the MAF distribution of GWAS hits. The black dashed line shows real schizophrenia GWAS hits for reference*.

## Notes

### Competing Interest Statement

The authors have declared no competing interest.

## References

[1] Galton F. Regression towards mediocrity in hereditary stature. The Journal of the Anthropo-logical Institute of Great Britain and Ireland. 1886;15:246–63.

[2] Falconer DS, Mackay TFC. Introduction to Quantitative Genetics. 4th ed. Harlow, Essex, England: Longman; 1996.

[3] Claussnitzer M, Cho JH, Collins R, Cox NJ, Dermitzakis ET, Hurles ME, et al. A brief history of human disease genetics. Nature. 2020;577(7789):179–89.

[4] Abdellaoui A, Yengo L, Verweij KJ, Visscher PM. 15 years of GWAS discovery: realizing the promise. The American Journal of Human Genetics. 2023;110(2):179–94.

[5] Canela-Xandri O, Rawlik K, Tenesa A. An atlas of genetic associations in UK Biobank. Nature genetics. 2018;50(11):1593–9.

[6] Kurki MI, Karjalainen J, Palta P, Sipilä TP, Kristiansson K, Donner KM, et al. FinnGen provides genetic insights from a well-phenotyped isolated population. Nature. 2023;613(7944):508–18.

[7] Koch E, Connally N, Baya N, Reeve M, Daly M, Neale B, et al. Genetic association data are broadly consistent with stabilizing selection shaping human common diseases and traits. bioRxiv. 2024:2024–06.

[8] Timpson NJ, Greenwood CM, Soranzo N, Lawson DJ, Richards JB. Genetic architecture: the shape of the genetic contribution to human traits and disease. Nature Reviews Genetics. 2018;19(2):110–24.

[9] Watanabe K, Stringer S, Frei O, Umićević Mirkov M, de Leeuw C, Polderman TJ, et al. A global overview of pleiotropy and genetic architecture in complex traits. Nature genetics. 2019;51(9):1339–48.

[10] Choi SW, Mak TSH, O’Reilly PF. Tutorial: a guide to performing polygenic risk score analy-ses. Nature protocols. 2020;15(9):2759–72.

[11] Wu T, Sham PC. On the transformation of genetic effect size from logit to liability scale. Behavior Genetics. 2021;51(3):215–22.

[12] Wang Y, Kanai M, Tan T, Kamariza M, Tsuo K, Yuan K, et al. Polygenic prediction across populations is influenced by ancestry, genetic architecture, and methodology. Cell Genomics. 2023;3(10).

[13] Sella G, Barton NH. Thinking about the evolution of complex traits in the era of genome-wide association studies. Annual review of genomics and human genetics. 2019;20(1):461–93.

[14] Simons YB, Bullaughey K, Hudson RR, Sella G. A population genetic interpretation of GWAS findings for human quantitative traits. PLoS biology. 2018;16(3):e2002985.

[15] Simons YB, Mostafavi H, Zhu H, Smith CJ, Pritchard JK, Sella G. Simple scaling laws control the genetic architectures of human complex traits. PLoS biology. 2025;23(10):e3003402.

[16] Johnson T, Barton N. Theoretical models of selection and mutation on quantitative traits. Philosophical Transactions of the Royal Society B: Biological Sciences. 2005;360(1459):1411–25.

[17] Walsh B, Lynch M. Evolution and selection of quantitative traits. Oxford University Press; 2018.

[18] Frei O, Holland D, Smeland OB, Shadrin AA, Fan CC, Maeland S, et al. Bivariate causal mix-ture model quantifies polygenic overlap between complex traits beyond genetic correlation. Nature communications. 2019;10(1):2417.

[19] Smith CJ, Sinnott-Armstrong N, Cichońska A, Julkunen H, Fauman EB, Würtz P, et al. In-tegrative analysis of metabolite GWAS illuminates the molecular basis of pleiotropy and genetic correlation. elife. 2022;11:e79348.

[20] Eyre-Walker A. Genetic architecture of a complex trait and its implications for fitness and genome-wide association studies. Proceedings of the National Academy of Sciences. 2010;107(Suppl_1):1752–6.

[21] Speed D, Hemani G, Johnson MR, Balding DJ. Improved heritability estimation from genome-wide SNPs. The American Journal of Human Genetics. 2012;91(6):1011–21.

[22] Agarwala V, Flannick J, Sunyaev S, Consortium G, Altshuler D. Evaluating empirical bounds on complex disease genetic architecture. Nature genetics. 2013;45(12):1418–27.

[23] Zeng J, De Vlaming R, Wu Y, Robinson MR, Lloyd-Jones LR, Yengo L, et al. Signatures of negative selection in the genetic architecture of human complex traits. Nature genetics. 2018;50(5):746–53.

[24] Schoech AP, Jordan DM, Loh PR, Gazal S, O’Connor LJ, Balick DJ, et al. Quantification of frequency-dependent genetic architectures in 25 UK Biobank traits reveals action of negative selection. Nature communications. 2019;10(1):790.

[25] Zeng J, Xue A, Jiang L, Lloyd-Jones LR, Wu Y, Wang H, et al. Widespread signatures of natural selection across human complex traits and functional genomic categories. Nature communications. 2021;12(1):1164.

[26] Finucane HK, Bulik-Sullivan B, Gusev A, Trynka G, Reshef Y, Loh PR, et al. Partitioning her-itability by functional annotation using genome-wide association summary statistics. Nature Genetics. 2015 Sep;47(11):1228–1235.

[27] Finucane HK, Reshef YA, Anttila V, Slowikowski K, Gusev A, Byrnes A, et al. Heritability enrichment of specifically expressed genes identifies disease-relevant tissues and cell types. Nature genetics. 2018;50(4):621–9.

[28] O’Connor LJ, Schoech AP, Hormozdiari F, Gazal S, Patterson N, Price AL. Extreme polygenicity of complex traits is explained by negative selection. The American Journal of Human Genetics. 2019;105(3):456–76.

[29] Omdahl AR, Weinstock JS, Keener R, Chhetri SB, Arvanitis M, Battle A. Sparse matrix factorization robust to sample sharing across GWASs reveals interpretable genetic components. The American Journal of Human Genetics. 2025;112(9):2178–97.

[30] O’Connor LJ, Sella G. Principled measures and estimates of trait polygenicity. bioRxiv. 2025:2025–07.

[31] Holland D, Frei O, Desikan R, Fan CC, Shadrin AA, Smeland OB, et al. Beyond SNP heritability: Polygenicity and discoverability of phenotypes estimated with a univariate Gaussian mixture model. PLoS Genetics. 2020;16(5):e1008612.

[32] Zhang Y, Qi G, Park JH, Chatterjee N. Estimation of complex effect-size distributions using summary-level statistics from genome-wide association studies across 32 complex traits. Nature genetics. 2018;50(9):1318–26.

[33] Patel RA, Weiß CL, Zhu H, Mostafavi H, Simons YB, Spence JP, et al. Characterizing selection on complex traits through conditional frequency spectra. Genetics. 2025;229(4):iyae210.

[34] Berg JJ, Li X, Riall K, Hayward LK, Sella G. Mutation–selection–drift balance models of complex diseases. Genetics. 2025:iyaf220.

[35] Neale lab - UK Biobank GWAS summary statistics; 2018. [Online; accessed 1-October-2022]. http://www.nealelab.is/uk-biobank/.

[36] Aragam KG, Jiang T, Goel A, Kanoni S, Wolford BN, Atri DS, et al. Discovery and system-atic characterization of risk variants and genes for coronary artery disease in over a million participants. Nature genetics. 2022;54(12):1803–15.

[37] Trubetskoy V, Pardiñas AF, Qi T, Panagiotaropoulou G, Awasthi S, Bigdeli TB, et al. Mapping genomic loci implicates genes and synaptic biology in schizophrenia. Nature. 2022;604(7906):502–8.

[38] Yang J, Ferreira T, Morris AP, Medland SE, of ANthropometric Traits (GIANT) Consortium GI, Replication DG, et al. Conditional and joint multiple-SNP analysis of GWAS summary statistics identifies additional variants influencing complex traits. Nature genetics. 2012;44(4):369–75.

[39] Spence JP, Mostafavi H, Ota M, Milind N, Gjorgjieva T, Smith CJ, et al. Specificity, length and luck drive gene rankings in association studies. Nature. 2025:1–8.

[40] Loos RJ, Yeo GS. The genetics of obesity: from discovery to biology. Nature Reviews Genetics. 2022;23(2):120–33.

[41] Liu Y, Chen S, Li Z, Morrison AC, Boerwinkle E, Lin X. ACAT: a fast and powerful p value combination method for rare-variant analysis in sequencing studies. The American Journal of Human Genetics. 2019;104(3):410–21.

[42] Howe LJ, Nivard MG, Morris TT, Hansen AF, Rasheed H, Cho Y, et al. Within-sibship genome-wide association analyses decrease bias in estimates of direct genetic effects. Nature genetics. 2022;54(5):581–92.

[43] Tan T, Jayashankar H, Guan J, Nehzati SM, Mir M, Bennett M, et al. Family-GWAS reveals effects of environment and mating on genetic associations. medRxiv. 2024:2024–10.

[44] Border R, Athanasiadis G, Buil A, Schork AJ, Cai N, Young AI, et al. Cross-trait assortative mating is widespread and inflates genetic correlation estimates. Science. 2022;378(6621):754–61.

[45] Davies NM, Hemani G, Neiderhiser JM, Martin HC, Mills MC, Visscher PM, et al. The importance of family-based sampling for biobanks. Nature. 2024;634(8035):795–803.

[46] Mostafavi H, Harpak A, Agarwal I, Conley D, Pritchard JK, Przeworski M. Variable prediction accuracy of polygenic scores within an ancestry group. elife. 2020;9:e48376.

[47] Hu S, Ferreira LA, Shi S, Hellenthal G, Marchini J, Lawson DJ, et al. Fine-scale population structure and widespread conservation of genetic effect sizes between human groups across traits. Nature Genetics. 2025:1–11.

[48] McMahon GM, Waikar SS. Biomarkers in nephrology. American journal of kidney diseases: the official journal of the National Kidney Foundation. 2013;62(1):165.

[49] de Groat WC, Griffiths D, Yoshimura N. Neural control of the lower urinary tract. Comprehensive Physiology. 2015;5(1):327.

[50] Falconer DS. The inheritance of liability to certain diseases, estimated from the incidence among relatives. Annals of human genetics. 1965;29(1):51–76.

[51] Mefford J, Smullen M, Zhang F, Sadowski M, Border R, Dahl A, et al. Beyond predictive R2: Quantile regression and non-equivalence tests reveal complex relationships of traits and polygenic scores. The American Journal of Human Genetics. 2025;112(6):1363–75.

[52] Speidel L, Forest M, Shi S, Myers SR. A method for genome-wide genealogy estimation for thousands of samples. Nature genetics. 2019;51(9):1321–9.

[53] Berisa T, Pickrell JK. Approximately independent linkage disequilibrium blocks in human populations. Bioinformatics. 2015;32(2):283.

[54] Ward LD, Kellis M. Evidence of abundant purifying selection in humans for recently acquired regulatory functions. Science. 2012;337(6102):1675–8.

[55] Rands CM, Meader S, Ponting CP, Lunter G. 8.2% of the human genome is constrained: variation in rates of turnover across functional element classes in the human lineage. PLoS genetics. 2014;10(7):e1004525.

[56] Christmas MJ, Kaplow IM, Genereux DP, Dong MX, Hughes GM, Li X, et al. Evolutionary constraint and innovation across hundreds of placental mammals. Science. 2023;380(6643):eabn3943.

[57] Wainschtein P, Zhang Y, Schwartzentruber J, Kassam I, Sidorenko J, Fiziev PP, et al. Estimation and mapping of the missing heritability of human phenotypes. Nature. 2025:1–9.

[58] Zeng T, Spence JP, Mostafavi H, Pritchard JK. Bayesian estimation of gene constraint from an evolutionary model with gene features. Nature Genetics. 2024 Jul;56(8):1632–1643.

[59] Uffelmann E, Huang QQ, Munung NS, De Vries J, Okada Y, Martin AR, et al. Genome-wide association studies. Nature reviews methods primers. 2021;1(1):59.

[60] Wray N, Pergadia M, Blackwood D, Penninx B, Gordon S, Nyholt D, et al. Genome-wide association study of major depressive disorder: new results, meta-analysis, and lessons learned. Molecular psychiatry. 2012;17(1):36–48.

[61] of the Psychiatric GWAS Consortium MDDWG, et al. A mega-analysis of genome-wide association studies for major depressive disorder. Molecular psychiatry. 2012;18(4):10–1038.

[62] Levinson DF, Mostafavi S, Milaneschi Y, Rivera M, Ripke S, Wray NR, et al. Genetic studies of major depressive disorder: why are there no genome-wide association study findings and what can we do about it? Biological psychiatry. 2014;76(7):510–2.

[63] Hyde CL, Nagle MW, Tian C, Chen X, Paciga SA, Wendland JR, et al. Identification of 15 genetic loci associated with risk of major depression in individuals of European descent. Nature genetics. 2016;48(9):1031–6.

[64] Howard DM, Adams MJ, Shirali M, Clarke TK, Marioni RE, Davies G, et al. Genome-wide association study of depression phenotypes in UK Biobank identifies variants in excitatory synaptic pathways. Nature communications. 2018;9(1):1470.

[65] Wray NR, Ripke S, Mattheisen M, Trzaskowski M, Byrne EM, Abdellaoui A, et al. Genomewide association analyses identify 44 risk variants and refine the genetic architecture of major depression. Nature genetics. 2018;50(5):668–81.

[66] Howard DM, Adams MJ, Clarke TK, Hafferty JD, Gibson J, Shirali M, et al. Genome-wide meta-analysis of depression identifies 102 independent variants and highlights the importance of the prefrontal brain regions. Nature neuroscience. 2019;22(3):343–52.

[67] Adams MJ, Streit F, Meng X, Awasthi S, Adey BN, Choi KW, et al. Trans-ancestry genomewide study of depression identifies 697 associations implicating cell types and pharmacotherapies. Cell. 2025;188(3):640–52.

[68] Schmitz-Abe K, Li Q, Greene S, Borrelli M, Luo S, Ramesh MC, et al. Unique signatures of highly constrained genes across publicly available genomic databases. Genetics in Medicine. 2025:101413.

[69] Consortium B, Anttila V, Bulik-Sullivan B, Finucane HK, Walters RK, Bras J, et al. Analysis of shared heritability in common disorders of the brain. Science. 2018;360(6395):eaap8757.

[70] Urbut SM, Wang G, Carbonetto P, Stephens M. Flexible statistical methods for estimating and testing effects in genomic studies with multiple conditions. Nature genetics. 2019;51(1):187–95.

[71] Boyle EA, Li YI, Pritchard JK. An expanded view of complex traits: from polygenic to omni-genic. Cell. 2017;169(7):1177–86.

[72] Liu X, Li YI, Pritchard JK. Trans effects on gene expression can drive omnigenic inheritance. Cell. 2019;177(4):1022–34.

[73] Liu JZ, Van Sommeren S, Huang H, Ng SC, Alberts R, Takahashi A, et al. Association analyses identify 38 susceptibility loci for inflammatory bowel disease and highlight shared genetic risk across populations. Nature genetics. 2015;47(9):979–86.

[74] Easton DF, Pooley KA, Dunning AM, Pharoah PD, Thompson D, Ballinger DG, et al. Genome-wide association study identifies novel breast cancer susceptibility loci. Nature. 2007;447(7148):1087–93.

[75] Voight BF, Scott LJ, Steinthorsdottir V, Morris AP, Dina C, Welch RP, et al. Twelve type 2 diabetes susceptibility loci identified through large-scale association analysis. Nature genetics. 2010;42(7):579–89.

[76] of Glucose MA, related traits Consortium (MAGIC) Investigators I, of ANthropometric Traits (GIANT) Consortium GI, Consortium AGENTDAT, Consortium SATDS, Shuldiner AR, et al. Large-scale association analysis provides insights into the genetic architecture and patho-physiology of type 2 diabetes. Nature genetics. 2012;44(9):981–90.

[77] A comprehensive 1000 Genomes–based genome-wide association meta-analysis of coronary artery disease. Nature genetics. 2015;47(10):1121–30.

[78] Pantelis C, Papadimitriou GN, Papiol S, Parkhomenko E, Pato MT, Paunio T, et al. Biological insights from 108 schizophrenia-associated genetic loci. Nature. 2014;511(7510):421–7.

[79] Fry A, Littlejohns TJ, Sudlow C, Doherty N, Adamska L, Sprosen T, et al. Comparison of sociodemographic and health-related characteristics of UK Biobank participants with those of the general population. American journal of epidemiology. 2017;186(9):1026–34.

[80] Bulik-Sullivan BK, Loh PR, Finucane HK, Ripke S, Yang J, of the Psychiatric Genomics Consortium SWG, et al. LD Score regression distinguishes confounding from polygenicity in genome-wide association studies. Nature genetics. 2015;47(3):291–5.

[81] Consortium GP, et al. A global reference for human genetic variation. Nature. 2015;526(7571):68.

[82] Chang CC, Chow CC, Tellier LC, Vattikuti S, Purcell SM, Lee JJ. Second-generation PLINK: rising to the challenge of larger and richer datasets. Gigascience. 2015;4(1):s13742–015.

[83] Milind N, Smith CJ, Zhu H, Spence JP, Pritchard JK. Buffering and non-monotonic behavior of gene dosage response curves for human complex traits. medRxiv. 2024 Nov. Available from: 10.1101/2024.11.11.24317065.

[84] Mbatchou J, Barnard L, Backman J, Marcketta A, Kosmicki JA, Ziyatdinov A, et al. Computationally efficient whole-genome regression for quantitative and binary traits. Nature genetics. 2021;53(7):1097–103.

[85] Zhu X, Stephens M. Bayesian large-scale multiple regression with summary statistics from genome-wide association studies. The annals of applied statistics. 2017;11(3):1561.

[86] Gillett AC, Vassos E, Lewis CM. Transforming summary statistics from logistic regression to the liability scale: application to genetic and environmental risk scores. Human Heredity. 2019;83(4):210–24.

[87] Bellenguez C, Küçükali F, Jansen IE, Kleineidam L, Moreno-Grau S, Amin N, et al. New insights into the genetic etiology of Alzheimer’s disease and related dementias. Nature genetics. 2022;54(4):412–36.

[88] Demontis D, Walters GB, Athanasiadis G, Walters R, Therrien K, Nielsen TT, et al. Genome-wide analyses of ADHD identify 27 risk loci, refine the genetic architecture and implicate several cognitive domains. Nature genetics. 2023;55(2):198–208.

[89] O’Connell KS, Koromina M, van der Veen T, Boltz T, David FS, Yang JMK, et al. Genomics yields biological and phenotypic insights into bipolar disorder. Nature. 2025:1–12.

[90] Zhang H, Ahearn TU, Lecarpentier J, Barnes D, Beesley J, Qi G, et al. Genome-wide asso-ciation study identifies 32 novel breast cancer susceptibility loci from overall and subtype-specific analyses. Nature genetics. 2020;52(6):572–81.

[91] De Lange KM, Moutsianas L, Lee JC, Lamb CA, Luo Y, Kennedy NA, et al. Genome-wide association study implicates immune activation of multiple integrin genes in inflammatory bowel disease. Nature genetics. 2017;49(2):256–61.

[92] Consortium*† IMSG, ANZgene, IIBDGC, WTCCC2. Multiple sclerosis genomic map implicates peripheral immune cells and microglia in susceptibility. Science. 2019;365(6460):eaav7188.

[93] Nalls MA, Blauwendraat C, Vallerga CL, Heilbron K, Bandres-Ciga S, Chang D, et al. Identi-fication of novel risk loci, causal insights, and heritable risk for Parkinson’s disease: a meta-analysis of genome-wide association studies. The Lancet Neurology. 2019;18(12):1091–102.

[94] Schumacher FR, Al Olama AA, Berndt SI, Benlloch S, Ahmed M, Saunders EJ, et al. Associ-ation analyses of more than 140,000 men identify 63 new prostate cancer susceptibility loci. Nature genetics. 2018;50(7):928–36.

[95] Ishigaki K, Sakaue S, Terao C, Luo Y, Sonehara K, Yamaguchi K, et al. Multi-ancestry genome-wide association analyses identify novel genetic mechanisms in rheumatoid arthritis. Nature genetics. 2022;54(11):1640–51.

[96] Xue A, Wu Y, Zhu Z, Zhang F, Kemper KE, Zheng Z, et al. Genome-wide association analyses identify 143 risk variants and putative regulatory mechanisms for type 2 diabetes. Nature communications. 2018;9(1):2941.

[97] Niu H, Álvarez-Álvarez I, Guillén-Grima F, Aguinaga-Ontoso I. Prevalence and inci-dence of Alzheimer’s disease in Europe: A meta-analysis. Neurología (English Edition). 2017;32(8):523–32.

[98] De Angelis R, Demuru E, Baili P, Troussard X, Katalinic A, Lopez MDC, et al. Complete cancer prevalence in Europe in 2020 by disease duration and country (EUROCARE-6): a population-based study. The Lancet Oncology. 2024;25(3):293–307.

[99] Uhlén M, Fagerberg L, Hallström BM, Lindskog C, Oksvold P, Mardinoglu A, et al. Tissue-based map of the human proteome. Science. 2015;347(6220):1260419.

